# Regulation of the histone H3K36 methyltransferase Set2 by the histone chaperone Spt6

**DOI:** 10.64898/2026.05.21.726836

**Authors:** Alexandra R. Elchert, Vanda Lux, James L. Warner, Tereza Nešporová, Václav Veverka, Fred Winston

## Abstract

Histone H3 lysine 36 methylation is a conserved histone modification that is critical for maintaining eukaryotic transcriptional fidelity and genomic stability. In *Saccharomyces cerevisiae*, this modification is catalyzed by Set2, an ortholog of the mammalian H3K36 methyltransferase SETD2. Previous genetic, biochemical, and structural studies showed that Set2 activity is repressed by a Set2 autoinhibitory domain (AID) and that activation requires the direct binding of the histone chaperone Spt6. To study the role of Spt6 and Set2 autoinhibition *in vivo*, we have isolated and analyzed multiple classes of Spt6 and Set2 mutants. Our results suggest an autoinhibited form of Set2 in which the catalytic domain is bound by the AID. In strong agreement with our genetic results, biophysical experiments demonstrate that the catalytic domain and AID physically interact, and that the autoinhibition mutants disrupt this interaction. Finally, RNA-seq and ChIP-seq studies show the importance of the Set2-Spt6 interaction for transcription and H3K36 methylation genome-wide. Taken together, our results support a model in which Set2 exists in an inactive, autoinhibited state *in vivo* through direct catalytic domain-AID interactions, with binding by Spt6 required to release this autoinhibited state.

## INTRODUCTION

During eukaryotic transcription, several different histone modifications occur that are required for normal chromatin structure and transcription (1). Among these, histone H3 lysine 36 (H3K36) methylation is a critical regulator of transcriptional fidelity and genome stability that is conserved from yeast to humans (2–15). While humans have several H3K36 methyltransferases, *Saccharomyces cerevisiae* has only one, Set2, which catalyzes all three possible forms of H3K36 methylation (mono-, di-, and tri-methylation) (2). Set2 is an ortholog of human SETD2 (7,16). There is evidence that most human H3K36 methyltransferases, including SETD2, act as tumor suppressors, demonstrating the importance of regulating H3K36 methylation for human health (3,7,17).

H3K36 methylation is deposited co-transcriptionally, resulting in enrichment along transcribed sequences (16,18–21). One primary function of H3K36 di- and trimethylation during transcription in *S. cerevisiae* is to promote the recruitment and activity of the Rpd3S histone deacetylase complex, which maintains low levels of histone acetylation within coding regions, ensuring a repressive environment (4,6,22–25). Consistent with this function, deletion of *SET2* results in widespread activation of aberrant intragenic transcription from within hundreds of genes (9,10,26).

Set2 has several conserved domains that are required for H3K36 methylation (7). The catalytic domain (CD) includes three smaller domains: the associated with SET (AWS), SET, and post-SET domains. The SET domain contains the catalytic pocket responsible for methylation of histone H3 lysine 36. Set2 also contains a conserved Set2-Rpb1 interaction (SRI) domain, which binds the Ser2- and Ser5-phosphorylated carboxy-terminal domain of Rpb1, the largest subunit of RNA polymerase II, during transcription elongation (18). Although the SRI domain is required for Set2 activity, it is not absolutely required for Set2 recruitment to chromatin (27). In addition to these domains, Set2 contains an autoinhibitory domain (AID) that negatively regulates its catalytic activity, as deletions and other mutations within the Set2 AID cause hyperactivity *in vitro* (28,29).

Previous work in both yeast and mammalian cells has suggested that Set2/SETD2 activity requires the histone chaperone Spt6. In yeast, several genetic studies showed that H3K36 methylation is undetectable in certain *spt6* mutants (4,27,30–32). Additionally, studies in mammalian cells indicated a role for Spt6 for normal levels of H3K36 methylation (33–35). A connection between yeast Set2 autoinhibition and Spt6 came from the isolation of Set2 AID mutants as suppressors of the *spt6* H3K36 methylation defect (28), suggesting that Spt6 is required to overcome Set2 autoinhibition. More recently, three studies used cryoEM and biochemical reconstitutions to show that H3K36 methylation *in vitro* requires a direct interaction between the Spt6 death like domain (DLD) and a specific region within Set2/SETD2, now termed the Spt6 Binding Region (SBR) (36–38). Together, those studies revealed an active conformation of Set2 and suggested that an alternative conformation existed for the autoinhibited state.

In this study, we have taken several approaches to gain new insights into the regulation of Set2 by Spt6. First, we assayed the requirement for each Spt6 domain for H3K36 methylation *in vivo* and found that, in agreement with the structural studies, the Spt6 DLD is required. We subsequently identified two residues in the DLD that are specifically required for H3K36 methylation *in vivo*. Second, we tested the requirement for the Set2 SBR for H3K36 methylation and found, surprisingly, that the SBR is partially required for Set2 autoinhibition. Third, to gain additional genetic evidence for an autoinhibited state of Set2, we identified several Set2 mutants with amino acid changes in the catalytic domain that bypass the requirement for Spt6. These mutants, along with previous Set2 AID mutants with the same phenotype, strongly support an AlphaFold 3-predicted model for an autoinhibited conformation of Set2 in which the Set2 autoinhibitory and catalytic domains directly interact. Fourth, using NMR, we demonstrated that these two domains do directly interact *in vitro* in a manner dependent upon the residues identified by our mutant analysis. Additional NMR studies showed that Set2 directly interacts with all four histones. Finally, RNA-seq and ChIP-seq experiments demonstrated strong mutant phenotypes when the Set2-Spt6 interaction is lost and additionally showed strong bypass of the requirement for Spt6 by the strongest Set2 suppressor mutant. Our results provide strong evidence for two states of Set2 *in vivo*: an autoinhibited conformation in which the Set2 CD binds the Set2 AID to repress Set2 activity, and an active state, dependent upon Spt6, in which the Set2 CD binds histone H3 to catalyze H3K36 methylation.

## MATERIALS AND METHODS

### Yeast strains

All *S. cerevisiae* strains in this study (Supplementary Table S1) were derived from S288C (39). Strains were generated through standard techniques, including transformations, crosses, two-step gene replacement (40), and CRISPR-Cas9 genome editing (41). All CRISPR-Cas9 edited strains were crossed to ensure 2:2 segregation of the mutant phenotype. The *S. pombe* strain used for the ChIP-seq spike-in control was FWP566 (28). To test for dominance, the suppressor strains were first crossed to a strain containing *spt6-RR* and the *FLO8-URA3* reporter, and the diploids were tested for growth on 5-FOA. For ChIP-seq experiments, *SET2* was fused to a 3HA epitope tag (42). Oligonucleotides and plasmids used for strain constructions are listed in Supplementary Tables S2 and S3.

### Growth media

*S. cerevisiae* were grown in liquid culture at 30°C in either YPD (1% yeast extract, 2% peptone, 2% glucose) or synthetic complete (SC) media lacking tryptophan (0.2% dropout mix, 0.145% yeast nitrogen base without amino acids and ammonium sulfate, 0.5% ammonium sulfate, 2% glucose), as indicated. For Spt6 depletion experiments, 3-indoleacetic acid (IAA; Sigma, I2886) was added to YPD to a final concentration of 25 μM. Solid media was created as follows: synthetic complete (SC) media lacking either lysine, uracil, or tryptophan (0.2% dropout mix, 0.145% yeast nitrogen base without amino acids and ammonium sulfate, 0.5% ammonium sulfate, 2% glucose, 2% agar); SC media with 0.005% uracil and 5-fluoroorotic acid (5-FOA, 1 mg/ml); or YPD with G418 (200 µg/ml), dimethyl sulfoxide (DMSO, 0.1%), or 1-naphthaleneacetic acid (NAA, 500 µM; Sigma, N0640). To test the suppression of the insertion mutation *lys2-128*δ (Spt^-^ phenotype (43,44)), cells were spotted on SC-lys. *S. pombe* liquid cultures were grown at 32°C in YES (0.5% yeast extract, 3% glucose, 225 mg/l each of adenine, histidine, leucine, uracil, and lysine).

### Western blotting and antibodies

Triplicate 10 ml cultures were grown to OD_600_ ≈ 0.6, pelleted, washed once with deionized (dH_2_O), and frozen at -70°C. Cell lysates were obtained as previously described (45). The treated cell extract was resuspended in 1X Laemmli Buffer (0.0625 M Tris-HCl pH 6.8, 2% SDS, 10% glycerol, 5% 2-mercaptoethanol, 0.01% bromophenol blue). The volume of culture to load on a gel was determined by normalizing to an OD_600_ equivalent of 12. Samples were loaded on 15% SDS-PAGE gels and transferred to PVDF Immobilon-FL (Millipore) membranes. The following antibodies were used for western blotting: 1:15,000 anti-H3 (generously provided by Karen Arndt), 1:2,500 anti-H3K36me2 (Abcam ab9049), 1:1,000 anti-H3K36me3 (Abcam ab9050), 1:8,000 anti-Pgk1 (Thermo 459250) 1:5,000 anti-HA (Abcam ab9110), 1:8,000 anti-Set2 (generously provided by Brian Strahl) 1:10,000 anti-Spt6 (generously provided by Tim Formosa), 1:5,000 anti-V5 (Thermo R960-25), and 1:10,000 anti-FLAG (Sigma M2). The following secondary antibodies were used: 1:10,000 IRDye 680RD goat anti-rabbit IgG (Licor) and 1:20,000 IRDye 800CW goat anti-mouse IgG (Licor). Quantification of western blots was done using ImageJ and plotted using GraphPad Prism.

### Yeast dilution plating (spot tests)

Yeast cultures were grown to saturation overnight from single colonies and normalized to OD_600_ ≈ 1, followed by five 10-fold dilutions, which were spotted onto solid media. Plates were incubated at 30°C unless noted otherwise. All experiments were performed in duplicate.

### Analysis of Spt6 domain deletions

To analyze the requirement for each Spt6 domain, plasmids were constructed that contained wild-type *SPT6* or the appropriate deletion mutation. The plasmids (Supplementary Table S3) were constructed by standard methods using PCR. The plasmids were used to transform yeast strain FY3544 and plated on SC-trp for two days at 30°C. After purification, a single colony of each transformant was inoculated individually into a flask with 25 ml of SC-trp. When cultures reached OD_600_ ≈ 0.6, an equal volume of 30°C media was added to dilute to OD_600_ ≈ 0.3. The cultures were split and treated with either DMSO (0.1%) or IAA (25 μM) and grown at 30°C for 90 minutes. Cells were pelleted, washed once with dH_2_O, and frozen at -70°C. Spt6 and H3K36 methylation protein levels were measured by western blotting. Phenotypes were assayed by spot tests on plates containing either DMSO (0.1%) or NAA (500 µM).

### AlphaFold 3 analysis

AlphaFold 3 (46) was used to predict the structures of the Set2 CD-AID (33–474) and Set2 CD-AID (33–474) bound to the Spt6 DLD (1019–1104). The top scoring models, with their accompanying pLDDT and PAE scores, are displayed. Structures were visualized in ChimeraX (47).

### Isolation and genetic analysis of *spt6-RR* suppressor mutations in *SET2*

To screen for new mutations in the 5’ portion (encoding the catalytic domain) of the *SET2* gene, several steps were taken. First, the 5’ portion of *SET2* was amplified from plasmid bARE10 (Supplementary Table S3) using the GeneMorph II Random Mutagenesis Kit (Agilent) and primers oARE306 and oARE307 (Supplementary Table S2). Second, the remaining sequence of bARE10, including the rest of *SET2* and the plasmid backbone, was amplified with Q5 High-Fidelity Polymerase (NEB) and purified by gel extraction. This amplification product was designed to overlap with the *SET2* amplification product by 60 base pairs at each end. Third, five independent cultures of yeast strain FY3590 were co-transformed with a mix of ∼300 ng of the mutagenized *SET2* 5’ region and 100 ng of purified backbone. Each transformation mix was equally split and plated on five SC-leu plates. Fourth, after two days of incubation at 30°C, the SC-leu plates (with a total of ∼8000 colonies) were replica plated to SC-leu 5-FOA plates and incubated at 30°C for two days to screen for suppressors (5-FOA resistant phenotype). Finally, 245 5-FOA resistant colonies were purified on SC-leu 5-FOA plates and of those, 23 plasmids were recovered into *E. coli*, purified, and sequenced (Plasmidsaurus). As each plasmid contained multiple mutations in *SET2* (Supplementary Table S4), candidate single mutations were created in strain FY3589 and verified by Sanger sequencing and genetic crosses.

### Protein expression and purification

The expression plasmids encoding different regions of *S. cerevisiae* Set2 (amino acids 33-260, 260-455, and 33-455), including both wild-type and mutant variants were derived from a pMCSG7 vector (T7 driven, ampicillin resistance). This vector encodes an N-terminal His6 affinity tag followed by the tobacco etch virus (TEV) protease recognition site. Upon TEV cleavage, the encoded proteins retained five amino acids (SNAAS) at their N-termini. Proteins were overexpressed in *Escherichia coli* BL21 (DE3) cells (NEB C2527I) harboring plasmid pRare2 for expression of tRNA molecules with rare codons (chloramphenicol resistance, 25 μg/ml). Transformants were plated and incubated overnight at 37°C. Colonies were used to inoculate cultures such that the starting OD_600_ was 0.05. Cultures were grown either using LB broth (Sigma) supplemented with 100 μg/ml ampicillin, 25 μg/ml chloramphenicol, and 0.5% glycerol, or for ^15^N or ^15^N/^13^C-labeled proteins, in minimal medium containing ^15^N-ammonium sulphate and ^13^C-glucose as the sole nitrogen and carbon sources and 100 μg/ml ampicillin and 25 μg/ml chloramphenicol. All cultures were grown at 37°C until induction of protein expression at OD_600_ ≈ 0.8 with 400 μM isopropyl-β-D-thiogalactopyranoside (IPTG). Protein expression was continued overnight at 18°C. Cells were harvested by centrifugation at 5000 g for 20 min and frozen before purification. Bacterial pellets were resuspended in lysis buffer (25 mM Tris–HCl, 1 M NaCl, 2 mM β-mercaptoethanol, 10 µM EDTA pH 7.5; 10 ml buffer per 1 g of pellet) containing protease inhibitors (cOmplete™, EDTA-free, Roche), DNase I (DN25 Sigma-Aldrich,2 units/ ml) and lysed by Emulsiflex C3 (ATA Scientific). The lysate was cleared by centrifugation (25000 g, 4°C, 30 min) and loaded on HIS-Select^®^ Nickel Affinity Gel resin (Sigma). The bound protein was eluted by imidazole buffer (25 mM Tris-HCl, 1 M NaCl, 250 mM imidazole, 2 mM β-mercaptoethanol, 10 µM EDTA, pH 7.5). The fractions containing purified protein were dialyzed in the presence of TEV protease (for His_6_ tag removal) in lysis buffer overnight. The second chromatography on HIS-Select^®^ Nickel Affinity Gel resin served for the removal of the TEV protease and proteins non-specifically interacting with the resin. The purified protein present in the flow-through fraction from HIS-Select^®^ Nickel Affinity gel resin was concentrated using a Vivaspin centrifugal filtration unit with 10 kDa cutoff. The final purification step was size exclusion chromatography Superdex 75 Increase 10/300 GL (Cytiva) for Set2 CD or AID and Superdex 200 Increase 10/300 GL (Cytiva) for Set2 (33–455) variants in buffer 25 mM Tris-HCl, 150 mM NaCl, 1 mM TCEP, 10 μM EDTA pH 7.5. The purest fractions were concentrated as before. For hydrogen-deuterium exchange experiments, the proteins were additionally purified using MonoQ 5/50 GL (Cytiva) chromatography in 25 mM Tris-HCl, 0.15 - 1M NaCl, 1 mM TCEP, 10 μM EDTA pH 7.5 buffer. The final concentration of the purified protein was determined on a NanoDrop spectrophotometer, and its purity was evaluated by SDS-PAGE followed by Coomassie Brilliant Blue. The protein was shock-frozen in liquid nitrogen and stored at -80°C. The Set2 mutants used were as follows: for the catalytic domain (residues 33-260) – wild type, I89L, N90A, L92F, L92H, L82I, N208Y, G260D, I89L/N208Y, and I89L/G260D; for the autoinhibitory domain (260–455) – wild type; and for the combined CD and AID domains (33–455) – wild type, G260D and I89L/G260D.

*Xenopus laevis* histone proteins (H2A, H2B, H3, H4) were expressed in *E. coli* BL21 CodonPlus (DE3) RIL cells (Agilent technologies 230245) as previously described (48,49). All cultures were grown at 37°C until induction of protein expression at OD_600_ ≈ 0.5 with 400 μM IPTG. Protein expression into inclusion bodies continued for 3 hours at 37°C. To briefly summarize the isolation procedure, after extraction from the inclusion bodies, histones were purified on an anion exchange column (HiPrep^®^ Q FF 16/10, Cytiva) and a cation exchange column (HiPrep^®^ SP FF 16/10, Cytiva) using anion (7 M deionized urea, 20 mM Tris–HCl, 0.2-1 M NaCl, 1 mM EDTA, 5 mM β-mercaptoethanol, pH 7.5) and cation (7 M deionized urea, 20 mM Na-acetate, 0.1-1 M NaCl, 1 mM EDTA, 5 mM β-mercaptoethanol, pH 5.2) exchange buffers. Pure fractions were pooled and dialyzed against 2 mM β-mercaptoethanol in MilliQ water, flash-frozen, lyophilized and stored at −80°C.

Histone dimers H3/H4 or H2A/H2B were refolded as previously described for octamers (48,50). Equimolar amounts of individual histones were resuspended in unfolding buffer (6 M GuHCl, 20 mM Tris–HCl, 5 mM DTT, pH 7.5), mixed in final concentration of 1 mg/ml and refolded during dialysis into 10 mM Tris–HCl, 2 M NaCl, 1 mM EDTA, 5 mM β-mercaptoethanol, pH 7.5 (dialysis repeated 3X). The sample was concentrated and applied to a Superdex^®^ 200 Increase 10/300 GL column (Cytiva) that was equilibrated in 25 mM Tris-HCl, 200 mM NaCl, 10 µM EDTA and 2 mM TCEP buffer pH 7.5. Histone dimer-containing fractions were pooled, concentrated, flash frozen and stored at -80°C.

### NMR spectroscopy

NMR spectra were acquired at 25°C on an 850 MHz Bruker Avance spectrometer using a triple-resonance (^15^N/^13^C/^1^H) cryoprobe. Samples used for backbone resonance assignment of the individual Set2 domains were prepared at 0.2-0.4 mM in a volume of 0.35 mL, in buffer (25 mM Tris-HCl pH 7.5, 150 mM NaCl, 1 mM TCEP, 10 μM EDTA, and 5% D2O/95% H2O) in a 5 mm Shigemi tube. A series of double- and triple-resonance spectra (50,51) were recorded to obtain sequence-specific backbone resonance assignments of Set2 constructs (catalytic domain residues 33-260, autoinhibitory domain 260-455). For the catalytic domain, only the region spanning residues 57–115 yielded sufficient signal quality in the triple-resonance spectra for backbone assignment. The backbone assignment of the autoinhibitory domain was nearly complete.

To monitor interaction between the two Set2 domains (CD or AID, wild-type or mutant forms), or the binding of Set2 domains to H2A/H2B or H3/H4 dimers, 2D ^15^N/^1^H SOFAST-HMQC spectra were recorded. Binding experiments were performed using 40 μM ^15^N-labeled protein titrated with unlabeled binding partner at molar ratios of 1:0.5, 1:1, 1:2, and 1:4 in a 3 mm NMR tube (0.16 mL sample volumes) using a modified buffer with 10% D_2_O/90% H_2_O. Because the affinities of the Set2 catalytic domain and autoinhibitory domain for histone dimers were higher, histone dimers were titrated at lower molar ratios of 1:0.2, 1:0.4, 1:0.6, and 1:0.8. NMR experiments with H2A/H2B dimers were performed in the same buffer supplemented with 200 mM NaCl, whereas experiments with H3/H4 dimers were performed with 300 mM NaCl to prevent histone precipitation.

NMR spectra were analyzed using NMRFAM-SPARKY (52). Peak intensities were exported together with their signal-to-noise ratio (SNR) values as SPARKY peak lists and analyzed using R. Residue-specific intensity ratios were calculated as the ratio of the peak intensity in the bound spectrum to that in the free spectrum. For each residue, the intensity ratio was calculated as: 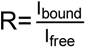. The uncertainty of each ratio was estimated by propagating the errors from the two peak measurements. The relative uncertainty of the ratio was calculated using the Pythagorean error-propagation approach: 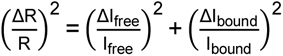. Because the uncertainty of each peak measurement was estimated from its signal-to-noise ratio, the relative ratio error was calculated as: 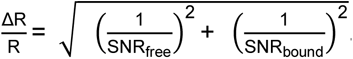. The absolute error of the ratio was then obtained by multiplying the relative uncertainty by the calculated ratio: 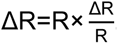. A minimum relative error threshold of 3% was applied to all ratios. Chemical shift perturbations (CSPs) were calculated using the weighted formula: 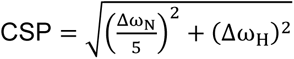 where Δω_N_ and Δω_H_ correspond to the chemical shift differences in the indirect (^15^N) and direct (^1^H) dimensions, respectively. Calculated intensity ratios, ratio errors, missing-peak annotations, and chemical shift perturbations were exported to Excel files for further inspection. Intensity ratio and CSP plots were generated using GraphPad Prism.

### Hydrogen deuterium exchange

Proteins set2 (33–455) wild type, G260D, and I89L/G260D were diluted in HEPES buffer (25 mM HEPES, 150 mM NaCl, 1 mM TCEP, pH 7.5) to a concentration of 8 µM. Samples were diluted with D_2_O HEPES buffer (25 mM HEPES, 150 mM NaCl, 1 mM TCEP, pD 7.5) to create a 90% deuterium labeling mixture. Labeling was conducted over multiple timepoints in 20°C (15 seconds; 2 minutes; 20 minutes) and 70 µl aliquots were quenched 1:1 (v/v) with 500 mM glycine, 6 M urea buffer (pH 2.3) and then flash frozen in liquid nitrogen. Frozen samples were thawed immediately prior to HDX-MS analysis. Samples were prepared in quadruplicate for each timepoint. Samples for sequence mapping were processed in the same way, except D_2_O was replaced with H_2_O in the labeling phase.

### Data collection for hydrogen-deuterium exchange

Data acquisition was carried out on SELECT SERIES Cyclic IMS instrument coupled to an M-class nanoACQUITY UPLC system and an HDX manager (Waters Corporation, MA, USA). All samples were prepared using LEAP PAL HDX autosampler and injected into the cold compartment of the autosampler set to 4°C for LC-MS analysis. The injected proteins were on-line digested on a protease column separated from the main cold LC compartment and set to 20°C (Nepenthesin-2/Pepsin, AP-PC-006, AffiPro), trapped on PepMap™ 100 C18 HPLC Trap Cartridge (C.N.: 160434, Thermo Scientific) and separated on Thermo Hypersil GOLD™ C18 Selectivity HPLC Column 1 mm x 50 mm (25002-051030, Thermo Scientific) using a 13.5 minute 5% to 35% solvent B gradient. Solvent A was 0.4% FA in H_2_O and solvent B was 0.4% FA in ACN. The flowrate was 100 µl/min for loading, digestion, and desalting, and 50 µl/min for separation. Data were collected in MSe mode for mapping and deuterated samples, using a standard ESI source with Source Temperature set to 100°C, desolvation gas temperature set to 350°C, desolvation flow to 800 l/hour, Cone Voltage 45V, Capillary voltage 3.15kV and Source Offset 20V. Manual quadrupole profile optimized for peptides was used (250;500;1200). The mass range was set to m/z 100–1200 and single pass ion mobility method with 5 ms of injection time and 4 ms of separation time was used. Detailed instrument setup parameters have been uploaded alongside raw data to the PRIDE data repository.

### Data analysis for hydrogen-deuterium exchange

Non-deuterated peptides were identified using PLGS, version 3.0.3 (Waters) using replicates of non-deuterated control samples. Processing parameters were as follows: automatic chromatographic peak width, automatic MS TOF resolution, lock mass window 0.25 Da, low energy threshold 150.0 counts and elevated energy threshold 30.0 counts. Workflow parameters were set to automatic peptide tolerance, 3 minimum fragment ion matches per peptide, 7 minimum fragment ion matches per protein, and the false discovery rate was set to 100 as instructed by the manufacturer for single protein datasets. Raw MS data were imported into Chronect^TM^ Studio – HDExaminerTMPRO, version 0.10.0.151 (Trajan Scientific and Medical) and the automatic validation of data with HDX-DIA processing was used. Corrected final deuteration values for wild-type and mutant proteins were further processed in R. Replicate measurements were grouped according to start and stop positions, sequence, labeling time and condition, and summarized as mean deuteration and standard deviation (SD). For each comparison, only peptides shared between wild type and the respective mutant were included. Residue-level weighted deuteration values and weighted SDs were then calculated and used to generate residue-level difference profiles. Residues covered by at least one significantly different peptide were included in plots generated in GraphPad Prism.

### RNA sequencing (RNA-seq)

RNA was purified from 50 ml cultures of cells grown to OD_600_ ≈ 0.6 in YPD using an approach adapted from a previously described method (53). To allow for later spike-in normalization, *S. pombe* cells were added at 10% of final cell concentration as measured using a hemocytometer. Briefly, cells were pelleted and then resuspended in 450 µl sodium acetate buffer (50 mM sodium acetate, 10 mM EDTA pH 5). Next, 50 µl of 10% SDS was added to each sample, followed by 500 µl of AE-equilibrated phenol (AE-Phenol). After vortexing, samples were incubated at 65°C for 4 minutes. The tubes were immediately transferred to a dry ice-ethanol bath for 1 minute. Samples were then centrifuged at 12,500 rpm for 10 minutes. The top aqueous layer was transferred to a fresh tube and combined with 300 µl AE-phenol and 300 µl chloroform and vortexed. Samples were spun at 12,500 rpm for 2 minutes, and the aqueous layer was transferred to an Eppendorf tube with 50 µl 3M Sodium acetate pH 5.3. After vortexing, 1.25 ml of 100% ethanol was added to each tube and vortexed. RNA was precipitated by incubating samples at -20°C for at least 2 hours. After incubation, samples were centrifuged at 12,500 rpm for 15 minutes. The pellet was washed with 500 µl 70% ethanol and centrifuged at 12,500 rpm for 5 minutes. The supernatant was removed and the pellet was dried for 10 minutes. The pellet was resuspended in 75 µl dH2O. RNA concentrations and quality were determined by TapeStation D1000. RNA-seq was performed by Plasmidsaurus (plasmidsaurus.com), using their Ultrafast RNA-seq service. The sequencing uses a 3’ end counting approach and is described in greater detail at https://plasmidsaurus.com/technical-documentation/rna.

### RNA-seq analysis

Sequencing results were aligned to the experimental (*S. cerevisiae*) and spike-in (*S. pombe*) genomes using Bowtie2 (54) and sorted using samtools (55,56). Spike-in normalization was performed using a custom python script. Spike-in normalized coverage tracks of reads mapping to each strand were generated using a custom python script and deepTools (57). Spearman correlation coefficients between all libraries were also calculated using deeptools. RNA-seq experiments were performed in biological triplicate. Overall, replicates correlated well with one another (Supplementary Figure S8A,B). Coverage tracks from the three replicates were averaged using deepTools. A table of all non-overlapping, protein-coding transcripts for each strand was generated using bedtools (58). DeepTools was used to extract the RNA-seq coverage for reads that were antisense to these annotations. These coverage tracks were merged and used to generate plots using a custom python script. Average antisense transcript abundance was defined as the mean coverage from 250 bp upstream of the TSS to the end of the transcript. The log_2_ fold-change in antisense transcript abundance for each transcript in each mutant over wild type was calculated and used to generate violin plots, scatter plots, and tables using a custom python script (see Data and code availability). Individual genomic loci were visualized using IGV (59).

### Chromatin immunoprecipitation and sequencing (ChIP-Seq)

Preparation of ChIP-seq libraries was performed as previously described (60), including a 10% *S. pombe* spike in at the step prior to immunoprecipitation, with the following changes: A saturated overnight culture inoculated from a single colony was used to grow each culture to OD_600_ ≈ 0.6 in YPD. For the immunoprecipitations (IPs), 300 µg of chromatin was used for histones, and 500 µg of chromatin was used for HA and Rpb1. The following antibody amounts were used for each IP reaction: 4 µl of anti-H3 (ab1791), 4 µl of anti-H3K36me2 (ab9049), 4 µl of anti-H3K36me3 (ab9050), 5 µl anti-HA (ab9110) (for Set2-HA), and 10 µl anti-Rpb1 (8WG16; Millipore 05-952). Libraries were built using the NEBNext Ultra II DNA library Prep Kit for Illumina (NEB) following manufacturer specifications. Libraries were indexed using NEBNext Multiplex Oligos for Illumina (NEB). Libraries were sequenced on a NovaSeq X Plus at the Harvard Medical School BPF NGS Genomics Core Facility.

### ChIP-seq analysis

Sequencing results were analyzed as previously described (60). Briefly, sequences were aligned to experimental (*S. cerevisiae*) and spike-in (*S. pombe*) genomes using Bowtie2 (54) and sorted using samtools (55,56). Spike-in normalization was performed using a custom python script and coverage tracks were generated using deepTools (57). Spearman correlation coefficients between all libraries were also calculated using deepTools. ChIP experiments were performed in biological triplicate, and replicates correlated well with one another (Supplementary Figure S9A). Coverage tracks from the three replicates were averaged using deepTools, and the averaged tracks were used to generate heatmap plots using a custom python script (see Data and code availability). Individual genomic loci were visualized using IGV (59).

## RESULTS

Our goal in these experiments was to study the regulation of Set2 autoinhibition. First, we set out to study the regulation of Set2 by Spt6 by identifying the regions of both proteins that are required for Set2 activity *in vivo*. Second, we sought to gain new understanding of the putative Set2 autoinhibited conformation, using both genetic and biophysical approaches.

### The Spt6 DLD is required for H3K36 methylation *in vivo*

To determine the requirement for individual Spt6 domains for H3K36 methylation *in vivo*, we first assessed the requirement for each domain for growth, as Spt6 is essential for viability. To do this, we used yeast strains in which endogenous Spt6 could be depleted using an auxin-inducible degron while Spt6 mutants were expressed from plasmids (Figure 1A). Growth was assayed by spot tests on rich media containing either 500 µM NAA to cause Spt6 depletion, or 0.1% DMSO as a non-depletion control.

**Figure 1.**
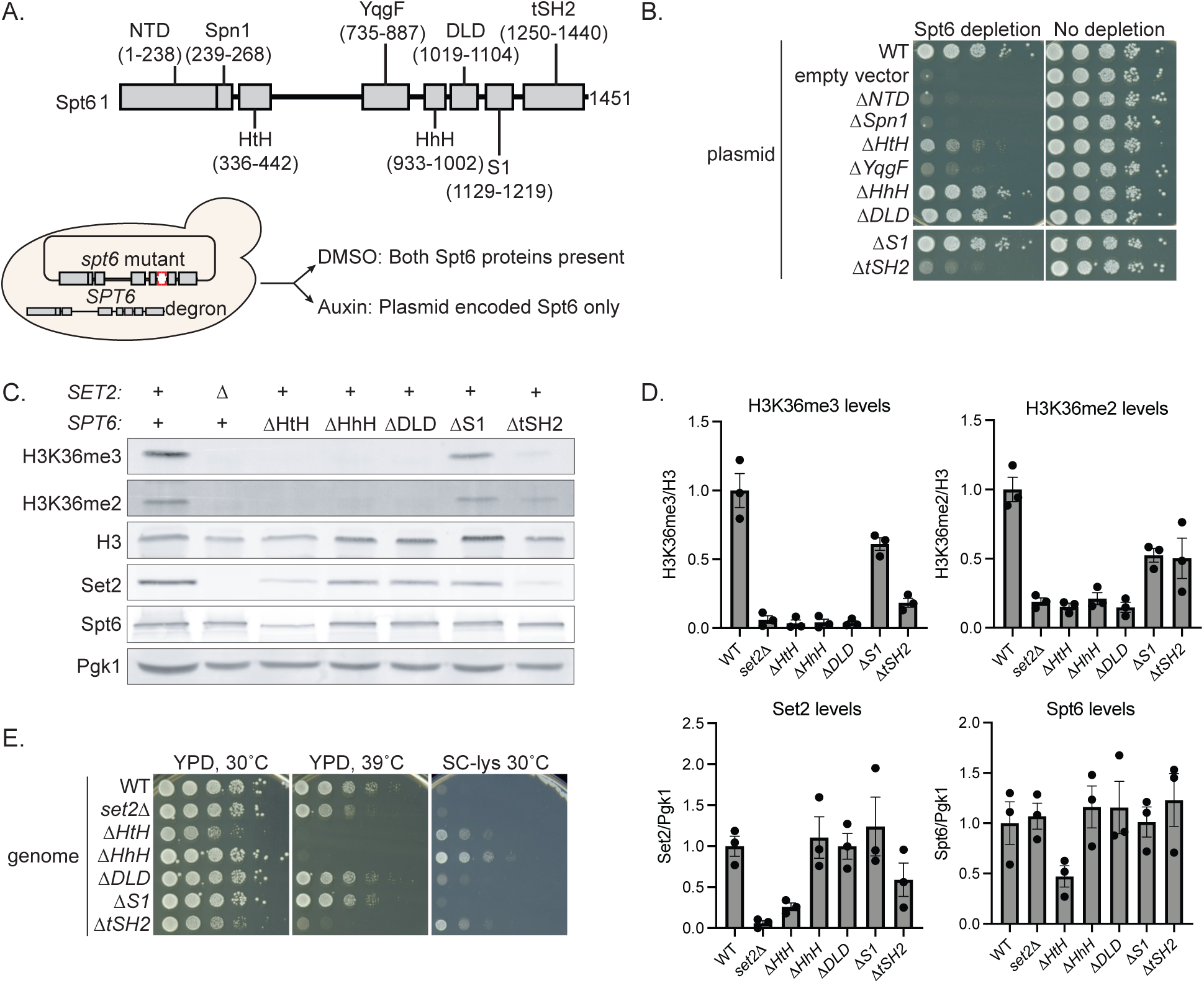
The Spt6 DLD is required for H3K36 methylation *in vivo*. **(A)** The top shows a diagram of Spt6 domains in *S. cerevisiae*. The numbers are amino acid positions (79). The bottom is a schematic of the depletion-complementation system used to determine the requirement for each Spt6 domain for viability. **(B)** Spot tests of the depletion-complementation strains. Shown are plates incubated for two days at 30°C. **(C)** Representative western blots showing histone H3K36 methylation levels and protein levels in *spt6* domain deletion mutants. **(D)** Quantification of three independent western blot experiments in (C). Each dot represents a single replicate. The error bars indicate the standard error of the mean. **(E)** Spot tests analyzing the temperature-sensitive and Spt^-^ phenotypes. Plates were incubated for two days at the indicated temperatures. Growth on SC-lys shows expression of the *lys2-128*δ reporter.

Our results showed that Spt6 domains were differentially required for growth (Figure 1B). Two mutants, *spt6ΔNTD* (as previously described (60)) and *spt6ΔSpn1* failed to grow under depletion conditions, similar to the empty vector control, while the *spt6ΔYqgF*, *spt6ΔtSH2,* and *spt6ΔHtH* mutants exhibited severely impaired growth. The results for *spt6ΔtSH2* are the same as previously reported (61–65). In contrast, the *spt6ΔHhH, spt6ΔDLD,* and *spt6ΔS1* mutants grew similarly to cells expressing wild type Spt6. We reasoned that if an Spt6 domain is primarily required for Set2 activity, that domain deletion would grow well, as the H3K36 methyltransferase Set2 is not required for normal growth on rich media (2).

To study the viable *spt6* mutants further, we integrated them into the genome, replacing wildtype *SPT6,* and performed several tests. First, we measured the levels of H3K36me2 and H3K36me3 by western blots. Our results showed that four of the mutants, *spt6ΔtSH2, spt6ΔHtH, spt6ΔHhH,* and *spt6ΔDLD*, had greatly reduced levels of both H3K36me2 and H3K36me3, similar to the *set2Δ* control (Figure 1C,D). Two of these mutants, *spt6ΔtSH2* and *spt6ΔHtH,* are unlikely to be specifically defective for Set2 activity as the H3K36 methylation defect in *spt6ΔtSH2* is likely caused by the decreased chromatin association of the Spt6ΔtSH2 mutant protein (64,66), and the H3K36 methylation defect in *spt6ΔHtH* is likely due to greatly reduced levels of both Spt6 and Set2 protein (Figure 1C,D). In contrast, the *spt6ΔHhH* and *spt6ΔDLD* mutants are more specifically defective for Set2 activity, as they have no detectable H3K36me2 or H3K36me3, yet both have wild-type levels of Spt6 and Set2 proteins.

To characterize the mutants in greater detail, we tested them for both temperature sensitivity and an Spt^-^ phenotype. This revealed that *spt6ΔHhH* is both Spt^-^ and temperature-sensitive, while *spt6ΔDLD* has only a very weak Spt^-^ phenotype (Figure 1E). The strong similarity between the *spt6ΔDLD* and *set2Δ* mutant phenotypes suggests that the Spt6 DLD domain is more specifically required for H3K36 methylation than the Spt6 HhH domain. Our *in vivo* results, taken together with recent structural and biochemical studies (36–38), strongly suggest that the Spt6 DLD directly interacts with and activates Set2 *in vivo*.

### DLD residues D1035 and E1038 are required for H3K36 methylation

While there is strong evidence for Spt6 DLD-Set2 interactions (36–38), other studies suggest that the Spt6 DLD also binds to the histone chaperone FACT (67) and the histone variant H2A.Z (68). This leaves open the possibility that the observed phenotypes are due to disruption of other DLD interactions. To gain additional evidence that the *spt6ΔDLD* phenotypes we observed are caused by effects on Spt6-Set2 binding, we used AlphaFold 3 to predict the Spt6-Set2 interaction. From this information, we predicted that two Spt6 residues, D1035 and E1038, were oriented to interact with Set2 (Figure 2A).

**Figure 2.**
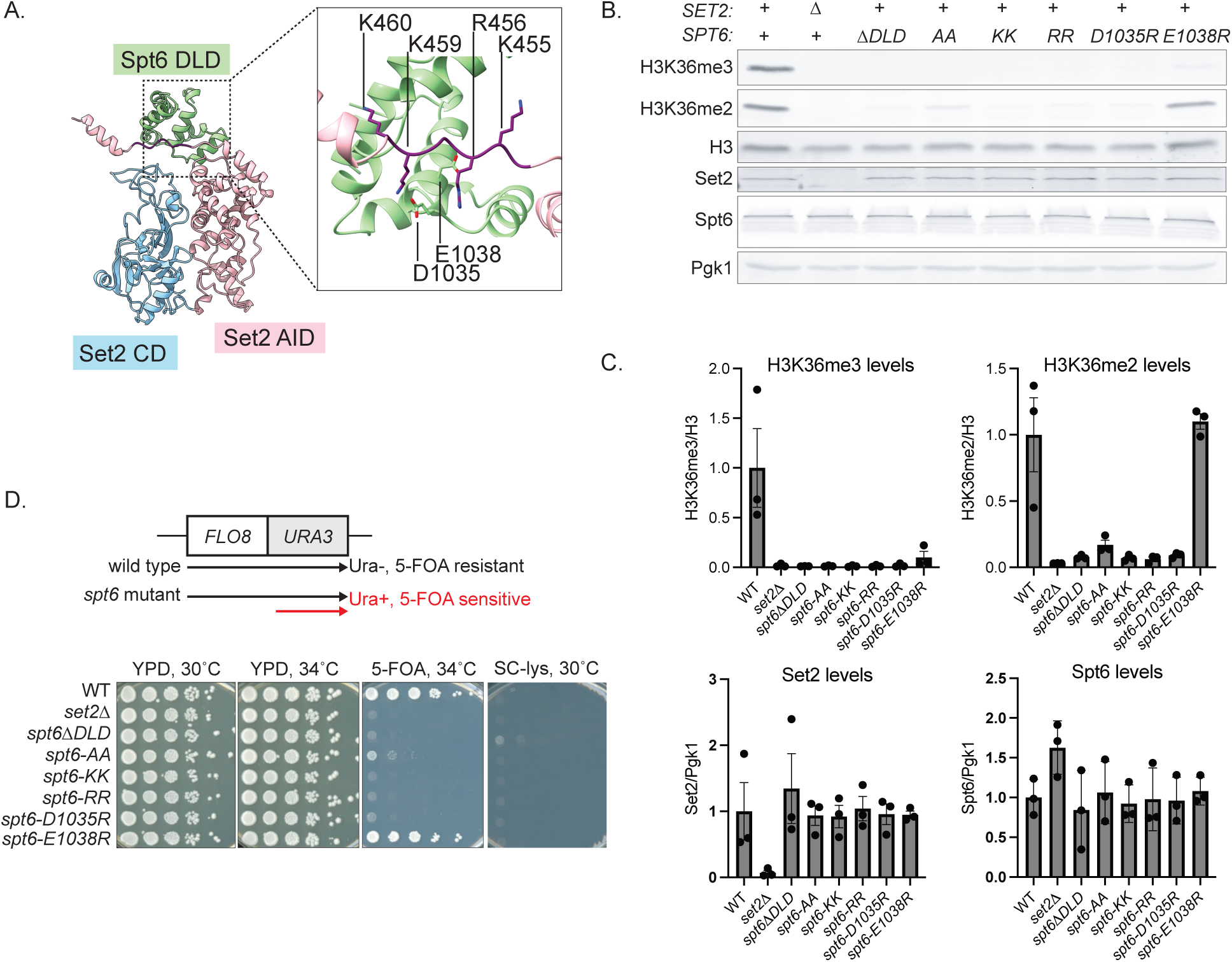
Spt6 DLD residues D1035 and E1038 are required for H3K36 methylation. **(A)** The AlphaFold3-predicted structure of the Set2 catalytic domain (CD) and autoinhibitory domain (AID) bound to the Spt6 DLD. Histone H3 was included during the generation of this structure but it is excluded from this image for clarity. The residues that comprise the Set2-Spt6 interaction are detailed in the inset. **(B)** Representative western blots showing H3K36 methylation levels and protein levels in the Spt6-DLD point mutants. **(C)** Quantification of three independent replicates of western blot images in (B). Each dot represents a single replicate. The error bars indicate the standard error of the mean. **(D)** On the top is shown a schematic of the *FLO8-URA3* cryptic intragenic transcription reporter. On the bottom are spot tests showing the Spt^-^ and cryptic intragenic transcription phenotypes. 5-FOA plates were incubated at 34°C as reported previously (28). Images of plates were taken after three days of incubation at 34°C.

To test if these Spt6 residues are important for H3K36 methylation, we constructed *spt6* mutants that changed one or both residues to arginine, lysine, or alanine. The choices of arginine and lysine were based on evidence that the interaction between Set2 and the Spt6 DLD is mediated by electrostatic interactions (36,37,69). Our results demonstrated that when D1035 and E1038 were both changed to arginine or lysine, the phenotypes were similar to *spt6ΔDLD*, as they both abolished detectable H3K36 methylation, while the changes to alanine had slightly less severe effects (Figure 2B,C). When we tested the individual amino acid changes (*spt6-D1035R* and s*pt6-E1038R*) we found that *spt6-D1035R* had no detectable H3K36me2 or H3K36me3, while *spt6-E1038R* had a partial defect, with reduced H3K36me3, but normal levels of H3K36me2 (Figure 2B,C).

We also compared these single and double amino acid changes to *spt6ΔDLD* by two sensitive genetic tests. First, we found that the amino acid changes caused no detectable Spt^-^ phenotype, making them more similar to a *set2Δ* mutant, compared to the weak but detectable Spt^-^phenotype caused by *spt6ΔDLD* (Figure 2D). We also assayed how the different DLD mutations affect transcription from a cryptic intragenic promoter with the *FLO8* gene, using a previously described *FLO8-URA3* fusion reporter (Figure 2D, (28)). The *spt6-RR, spt6-KK,* and *spt6-D1035R* mutants were all 5-FOA-sensitive, indicating expression of cryptic transcription. In line with the low levels of H3K36me2 present in *spt6-AA*, this mutant exhibited weak growth on 5-FOA. The *spt6-E1038R* mutant grew similarly to wild type on 5-FOA, indicating no cryptic transcription, consistent with a previous report that the *FLO8* cryptic intragenic promoter can be repressed by H3K36me2 (22). Together, these results have shown that specific Spt6 amino acids predicted to interact with Set2 are required for H3K36 methylation. Furthermore, a change of one of these amino acids, D1035, causes mutant phenotypes identical to loss of Set2.

### Analysis of the Spt6 binding region within Set2

Recent studies showed that the Spt6 DLD binds to a conserved 5-10 amino acid sequence in Set2/SETD2 (the Spt6 binding region, or SBR) that is required for H3K36me3 *in vitro* (36–38). One of these studies also showed that a mutation in the *S. cerevisiae* Set2 SBR, changing five consecutive amino acids to alanines, impaired H3K36me3 *in vivo* (38). To further study the Set2 SBR *in vivo* and the possible role of electrostatic interactions between Spt6 and Set2, we constructed a Set2 SBR mutant that changed four positively charged SBR residues (K455, R456, K459, K460 (Figure 2A)), each to negatively charged glutamic acid (*set2-4E*) and assayed the mutant by westerns for H3K36me2/me3 levels and by spot tests for cryptic transcription using our *FLO8-URA3* reporter. Our results showed that the *set2-4E* mutant had no detectable H3K36me3, but still had near-normal levels of H3K36me2 (Supplementary Figure S1C-E).

As the Spt6 DLD-Set2 SBR interaction is believed to be electrostatic (38), we sought to obtain evidence for this *in vivo* by combining the *set2-4E* mutant (basic to acidic changes) with reciprocal *spt6* DLD mutations (acidic to basic changes) to test whether they might confer mutual suppression. We combined the *set2-4E* mutation with three different *spt6* mutations that changed acidic residues in the Spt6 DLD to either lysine or arginine. As a control, we also combined the *set2-4E* mutation with *spt6ΔDLD*, which, by our model, should be unable to restore a Set2-Spt6 interaction by electrostatic interactions. Analysis of these mutants yielded unexpected results. Combining *set2-4E* with *spt6-KK*, *spt6-RR,* or *spt6-1035R* resulted in a greatly reduced mutant phenotype compared to the *spt6* mutants, approximately the same as the *set2-4E* mutant phenotype (Supplementary Figure S1C-E). However, the *set2-4E spt6ΔDLD* double mutant also had a weaker mutant phenotype, a result inconsistent with suppression caused by restoration of electrostatic interactions. While these results were uninformative with respect to the nature of the Set2 SBR-Spt6 DLD interactions, the ability of *set2-4E* to suppress *spt6ΔDLD* suggests that the Set2 SBR region functions in a more complex way than simply by binding to the Spt6 DLD and is likely required for Set2 autoinhibition, as discussed later.

### Genetic evidence that the Set2 catalytic and autoinhibitory domains directly interact to repress Set2 activity

Previous studies provided evidence for two conformations of Set2: an active conformation, when Set2 interacts with Spt6 and a nucleosome to methylate H3K36, and an autoinhibited conformation, in the absence of Spt6 and a nucleosome, in which the Set2 autoinhibitory domain (AID) represses Set2 activity by directly interacting with the Set2 catalytic domain (CD). While there is substantial structural and biochemical data in support of the active structure (36–38), there is primarily genetic evidence for the putative autoinhibited structure (28,29,37). To gain additional evidence for this autoinhibited conformation, we modeled Set2 using AlphaFold 3, which predicted direct interactions between the CD and AID. Interestingly, the previously isolated Set2 AID mutants that bypass the requirement for Spt6 (28) are located at this predicted interface (Figure 3C; Supplementary Figure S3A,B). However, the mutant hunt that discovered the AID mutants did not identify any mutants on the opposing side of the interface, in the CD. The isolation of such mutants would strengthen support for the AlphaFold 3 model. We note that such mutants are expected to be much rarer than those in the AID, as amino acid changes in the CD might impair Set2 catalytic activity and Set2-histone interactions.

**Figure 3.**
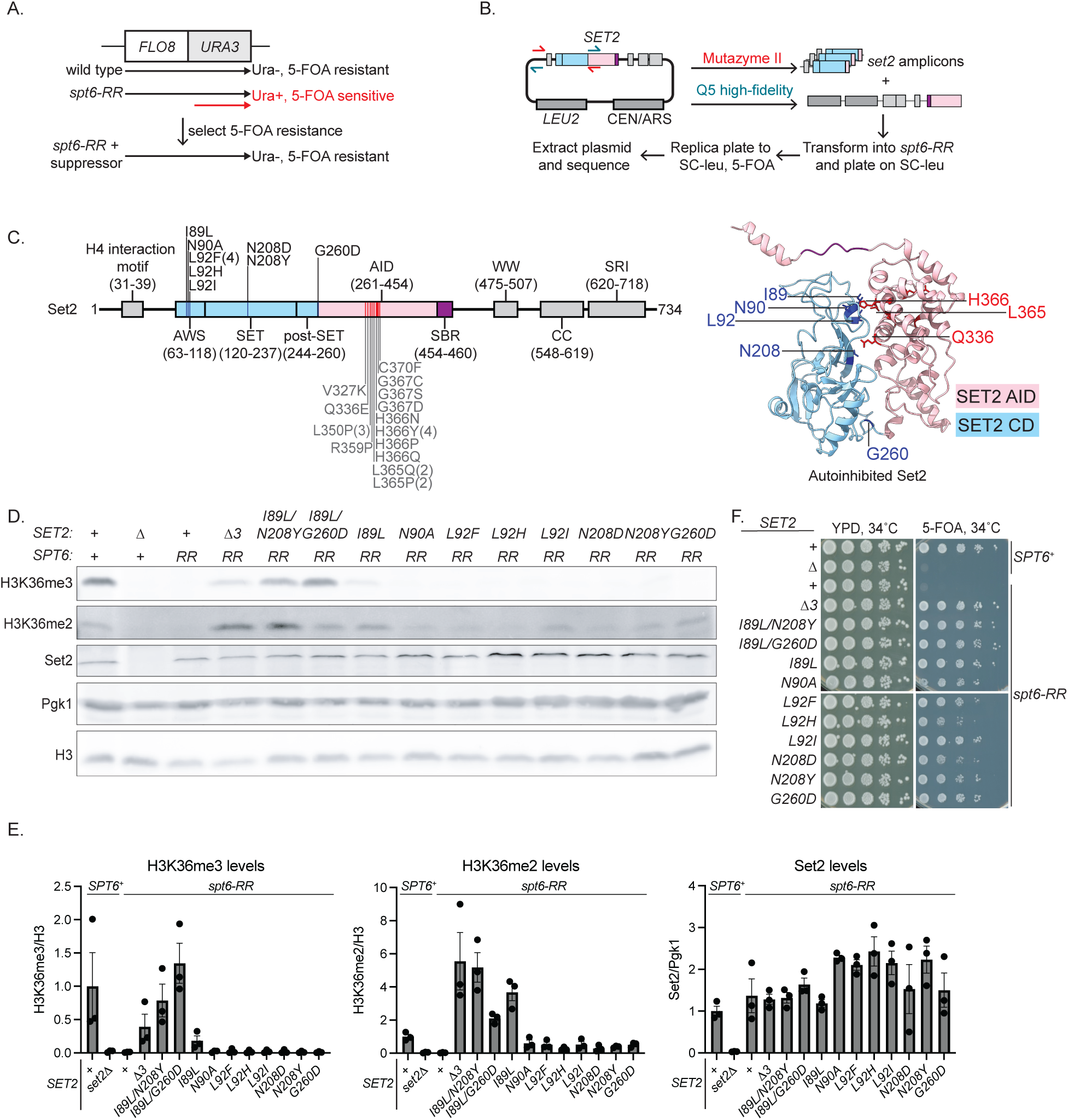
Identification of Set2 catalytic domain mutations that suppress the requirement for Spt6. **(A)** The diagram shows the reporter used to identify mutants. **(B)** Overview of the plasmid-mutagenesis screen used to isolate suppressor mutations within the Set2 CD. The small arrows indicate the positions of primers for amplification of the *SET2* catalytic domain (red) or the remainder of the plasmid (blue). **(C)** On the left is a Set2 domain diagram indicating the amino acid changes within the catalytic domain (in dark blue) that were isolated in the suppressor selection. The mutations within the AID (in red) were previously reported (28). On the right is an AlphaFold3-predicted structure of Set2 CD-AID (33–474) alone (autoinhibited), showing the positions of the mutations. **(D)** Representative western blots showing H3K36 methylation levels and protein levels in the *SET2* suppressors of *spt6-RR*. **(E)** Quantification of three independent western blot images in (D). Each dot represents a single replicate. The error bars indicate the standard error of the mean. **(F)** Spot test analysis of *FLO8-URA3* expression in the *spt6-RR* suppressors. Plates were incubated for two days at 34°C.

To identify *set2* mutations in the CD that bypass the requirement for Spt6, we took two approaches. First, using AlphaFold 3 as a guide, we constructed two *set2* mutants to test the amino acid substitutions N90A and T93A, which seemed likely to impair the interaction between the Set2 CD and AID. We tested whether these two mutations could suppress a mutation in the Spt6 DLD, *spt6-RR,* by assaying the *FLO8-URA3* intragenic transcription reporter (reporter shown in Figure 3A). Our results showed that while *set2-T93A* failed to suppress *spt6-RR, set2-N90A* was a suppressor (Supplementary Figure S2). The identification of the *set2-N90A* mutant as a suppressor of *spt6-RR* suggested that additional suppressors in the CD might be possible.

To identify additional *spt6-RR* suppressors in the Set2 CD, we performed an unbiased mutant screen of the Set2 CD. To do this, we used a plasmid-encoded copy of *SET2* to specifically target the region of *SET2* 5’ of the AID for PCR mutagenesis (Figure 3B). From a screen of approximately 8,000 transformants, we identified 23 candidates, from which we extracted and sequenced the mutant plasmids. While each plasmid carried multiple mutations in *SET2* (Supplementary Table S4), we identified seven different mutants with substitutions at four amino acid positions that we speculated to be causative, as all four amino acid positions are at or near the predicted interface of the Set2 CD and AID. (These residues are distal from the Set2 residues (Y149, F174, M177, F234, and Y236) required for catalysis (70).) Furthermore, three of the four amino acid positions identified by our mutations (I89, L92, N208) are predicted to directly face the positions previously identified by the AID mutants that bypass the requirement for Spt6 (Figure 3C) (71) and I89, N90, L92, and G260 are conserved in human SETD2 (Supplementary Figure S3C). As described below, these mutations are dominant, so we refer to the mutations using *SET2* nomenclature.

We took several steps to further characterize the new *SET2* suppressor mutations. We first constructed yeast strains with each single *SET2* mutation and demonstrated that suppression of *spt6-RR* segregates 2:2 in crosses, confirming that each single *SET2* mutation causes suppression. Next, we measured the degree of suppression of *spt6-RR* by western blots to assay the level of H3K36me2 and H3K36me3. We found that all of the *SET2* suppressors restored H3K36me2 to a detectable degree. The strongest suppressor, *SET2-I89L*, was comparable in strength to the strongest suppressors in the Set2 AID domain (for example, *SET2*Δ*3* (28); Figure 3D-F). We then tested the *SET2* suppressor mutations in a wild type *SPT6^+^* background to test for possible effects on H3K36 methylation levels (Supplementary Figure S3D-F). While most of the *SET2* mutants had levels of H3K36me3 equal to or greater than wild type, one mutant, *SET2-N90A*, had reduced levels of H3K36me3, likely accounting for its apparent weak suppression of *spt6-RR* as assayed by westerns. As described later, this may be caused by reduced Set2-histone interactions in this mutant. Finally, we tested three of the suppressor mutants for dominance and found that they were all dominant, ranging from modest to strong dominance (Supplementary Figure S3G). Together, these results provided strong evidence that the autoinhibited conformation predicted by AlphaFold 3 occurs *in vivo* and is required for the Spt6 dependence of Set2 activity.

Our strongest mutant, *SET2-I89L*, had a conservative amino acid change, suggesting that only certain amino acids in this position might confer suppression without impairing Set2 function. To test this, we constructed three additional amino acid changes at this position, to another aliphatic amino acid (valine) and to two aromatic amino acids (tyrosine and tryptophan). We then tested these mutants for H3K36 methylation levels in both *spt6-RR* and *SPT6^+^* backgrounds (Supplementary Figure S4A,B) and found that only the original mutant, *SET2-I89L*, conferred strong suppression. These data suggest that residue I89 likely serves a critical role in the regulation and activity of Set2.

Finally, as our new suppressors caused amino acid changes in both the AWS and SET regions within the CD, we tested whether combining mutations from these two classes would provide even stronger suppression. Our results showed that two double mutants, *SET2-I89L/N208Y and SET2-I89L/G260D,* are stronger suppressors than any of the single mutants, with the *SET2-I89L/G260D spt6-RR* mutant having nearly wild-type H3K36 methylation levels (Figure 3D-F).

These results, taken with our previously isolated Set2 AID mutants and AlphaFold 3 modeling, provide strong support for a Set2 conformation in which the AID directly contacts the CD to inhibit catalytic activity.

### Direct binding of the Set2 CD to the Set2 AID is disrupted by *SET2* suppressor mutations

Our *SET2* suppressor and AlphaFold 3 analyses, combined with previous genetic and biochemical results (28,29,37), suggested that the Set2 AID and CDs directly interact. To test this, we used nuclear magnetic resonance (NMR) spectroscopy to measure binding of the Set2 CD (amino acids 33-260) to the Set2 AID (amino acids 260-455). In the first set of experiments, we found that the CD bound to ^15^N-labeled AID, with the interactions largely coinciding with the amino acid positions identified by the previously isolated AID mutants (Figure 4A,B). Importantly, this binding was disrupted when we assayed the I89L mutant CD (Figure 4C,D). In a complementary approach, we observed that the AID bound to ^15^N-labeled CD (Figure 4E,F,H), again with the binding disrupted by the I89L mutant CD (Figure 4E,G,H). In addition, we found that the CD-AID binding was disrupted by most of the other CD suppressor mutants that were tested (Supplementary Figure S5). This strong convergence of the NMR results with our genetic results have established that the Set2 AID and Set2 CD directly interact and that the Set2 CD amino acids identified by our mutants are required for that interaction.

**Figure 4.**
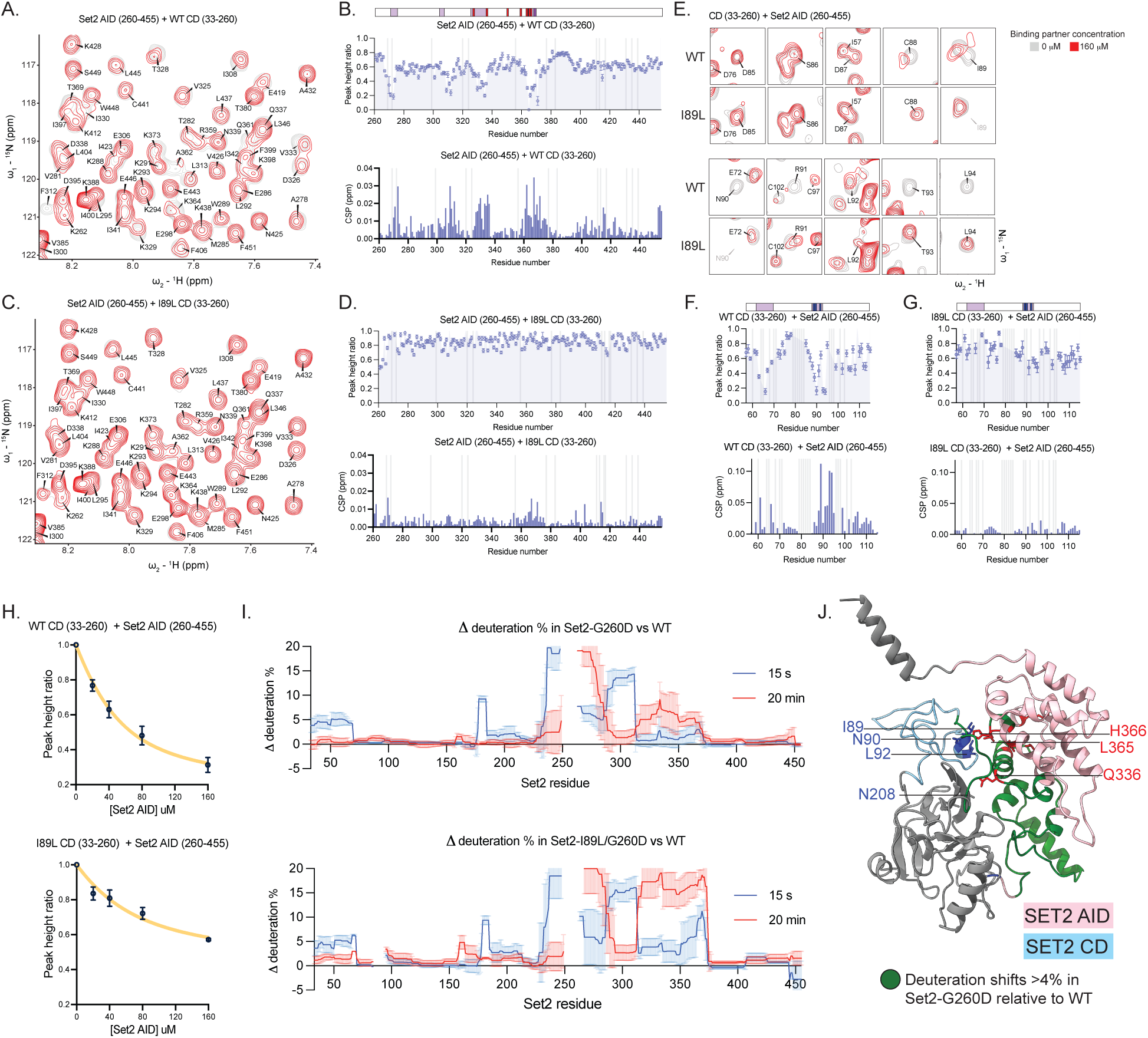
Analysis of Set2 CD-AID interactions using NMR and hydrogen-deuterium exchange coupled to mass spectrometry. **(A)** Overlaid 2D ^15^N/^1^H SOFAST-HMQC spectra of ^15^N labeled Set2 AID (260–455) in the absence (gray) or presence (red) of a four-fold molar excess of unlabeled Set2 CD (33–260). **(B)** The color bar above the plots shows the location within the Set2 AID of previously identified suppressor mutations (red) and AlphaFold-predicted AID-CD interface residues (purple). The top plot shows the backbone amide NMR signal intensity ratios obtained for Set2 AID upon titration with wild type Set2 CD. A lower peak height ratio indicates evidence of binding. The bottom plot shows quantification of backbone amide chemical shift perturbations (CSPs) in the Set2 AID induced upon addition of Set2 CD (33–260). A higher chemical shift perturbation indicates evidence of binding. The data obtained using four-fold molar excess of Set2 CD are shown. Residues that could not be assigned are shown as gray bars. **(C)** Plot as in (A) but with I89L-containing Set2 CD. **(D)** Plot as in (B) but with I89L-containing Set2 CD. **(E)** Overlaid 2D ^15^N/^1^H SOFAST-HMQC spectra of ^15^N labeled Set2 CD (33–260) in the absence (gray) or presence (red) of a four-fold molar excess of unlabeled Set2 AID (260–455). A subset of signals, for amino acid residues 85-94, are shown, as we were able to obtain a comprehensive assignment only for the AWS region of the Set2 CD. **(F)** The color bar above the plots shows the location within the Set2 CD of suppressor mutations described in this paper (blue) and AlphaFold-predicted CD-AID interface residues (purple). The top plot shows the backbone amide NMR signal intensity ratios obtained for wild type Set2 CD upon titration with the wild type Set2 AID. The bottom plot shows quantification of backbone amide chemical shift perturbations (CSPs) in the Set2 CD (only assigned residues) induced upon addition of Set2 AID (260–455). The data obtained using four-fold molar excess of Set2 AID are shown. Residues that could not be assigned are shown in gray. **(G)** Plots as in (F) but with I89L-containing Set2 CD. **(H)** Normalized and averaged intensities of 10 most perturbed peaks of Set2 CD wild type (left) and I89L-containing Set2 CD (right) upon titration with Set2 AID (260–455). **(I)** The top blot shows the difference in percent deuteration between Set2-CD-AID (33–455) G260D and wild type protein at 15 seconds and 20 minutes. The bottom blot is the same as top, but for Set2-CD-AID (33–455) I89L/G260D. Positive values indicate higher deuterium uptake and increased solvent accessibility relative to wild type. Gaps in each graph indicate regions for which directly equivalent peptides were not detected in the respective mutant and wild type pair. **(J)** AlphaFold3 predicted structure of Set2-CD-AID (33–474) with CD (33–260) in blue and AID (261–474) in pink with the residues with deuteration difference >4% in Set2-G260D relative to wild type are colored in red. The portions that are not present (456–474) or not assigned (33–56, 116–259) in NMR plots are colored gray.

One suppressor mutant of interest was Set2-G260D, as G260 was not predicted to be part of the CD-AID interface (Figure 3C). Given its position at the very end of the CD and AID fragments used for NMR, we were unable to assay the effects of G260D by this approach. Therefore, we used hydrogen-deuterium exchange coupled to mass spectrometry (HDX-MS) to analyze a Set2 construct containing both the CD and AID (33–455), comparing deuterium uptake in wild-type Set2 to two mutants: Set2-G260D and Set2-I89L/G260D. Our results showed significant differences in the percent deuterium uptake within both fast-exchange loops and beta sheets within the AWS-SET and AID, and within the slow-exchange helices within the AID (Figure 4I; Supplementary Figure S6). Consistent with the genetic results, there were greater differences for the Set2-I89L/G260D double mutant compared to the Set2-G260D single mutant (Figure 4I). These solvent-accessible regions in Set2-G260D and Set2-I89L/G260D indicate a large conformational change relative to wild type (Figure 4J), consistent with reduced autoinhibition.

### Evidence that the Set2 CD binds both Set2 AID and histone H3

Our NMR results showed that some of the weaker suppressors, including Set2-L92F and Set2-N90A, abolished detectable AID-CD interactions (Supplementary Figure S5). This apparent discrepancy, between a modest suppression phenotype and a strong loss-of-binding phenotype, might be caused by the mutants impairing interactions with both Set2 AID and with histone H3. Indeed, several of the amino acids identified by our suppressors are predicted to also interact with both the AID and the histone H3 αN helix (Figure 5A). This possibility is consistent with both the NMR results and reduced levels of H3K36me2 or H3K36me3 observed for several of these mutants in an *SPT6^+^*background (Supplementary Figure S3D-E).

**Figure 5.**
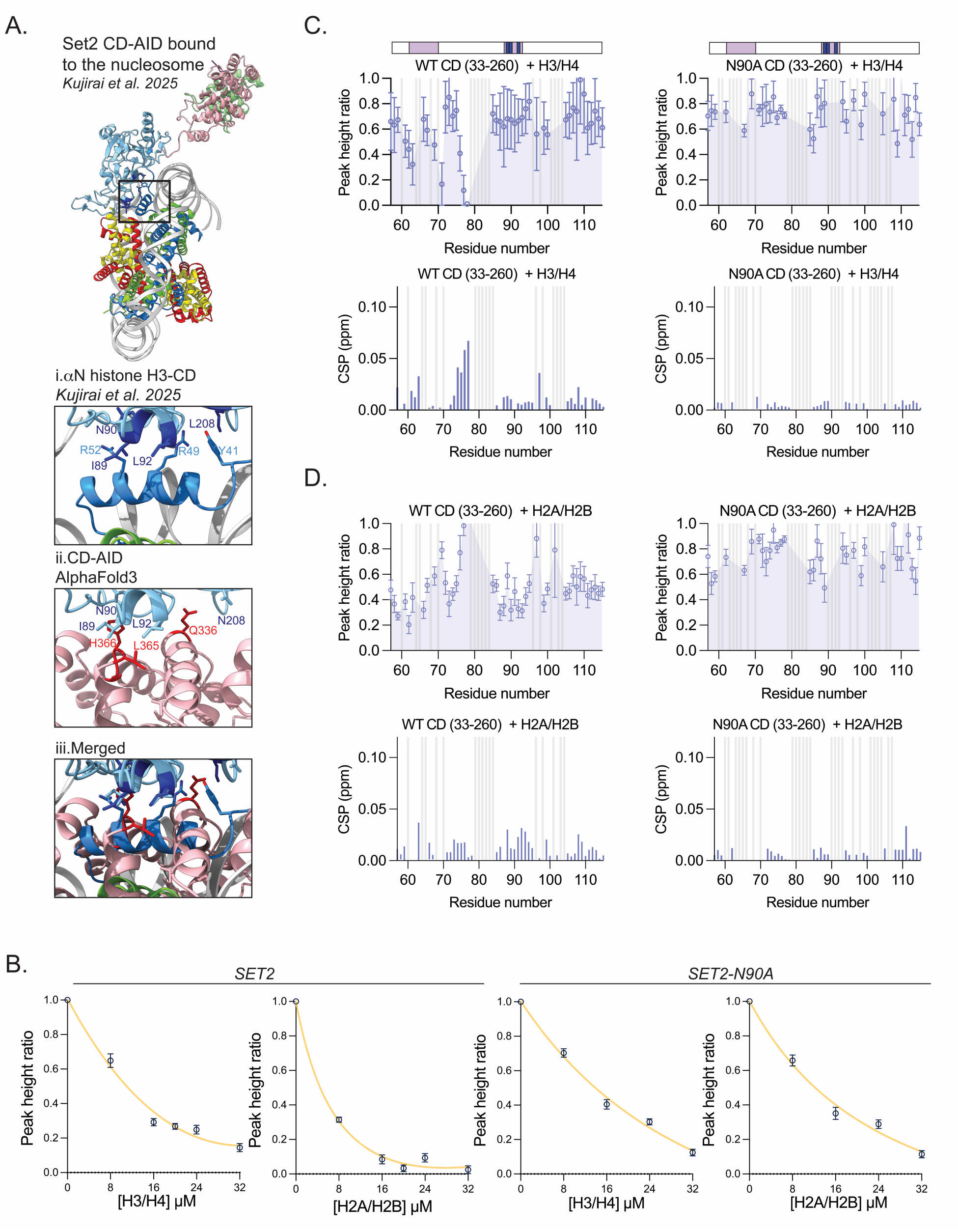
Analysis of Set2 CD interactions with core histones. **(A)** Structure of Set2 (CD in blue, AID in pink) and the Spt6-DLD (green) bound to the nucleosome with the elongation complex PDB:9WMS (38). Other proteins in the elongation complex are hidden for clarity. Detailed view of i. αN histone H3 and Set2 CD interactions in structure (PDB: 9WMS) ii. Predicted CD-AID interactions by AlphaFold3 and iii. Structures i and ii merged. **(B)** Normalized and averaged intensities of 10 most perturbed peaks of Set2 CD wild type (left) and N90A-containing Set2 CD (right) upon titration with either histone H3/H4 or H2A/H2B dimer. **(C)** The color bar above the plots shows the location within the Set2 CD of previously identified suppressor mutations (blue) and AlphaFold-predicted AID-CD interface residues (purple). The top plots show the NMR signal intensity ratios obtained for wild type or N90A-containing Set2 CD mutant upon titration with the histone H3/H4 dimer. A lower peak height ratio indicates evidence of binding. The bottom plots show quantification of backbone amide chemical shift perturbations (CSPs) in the Set2 CD induced upon addition of histone H3/H4 dimer. A higher chemical shift perturbation indicates evidence of binding. Data obtained at a 1:0.8 molar ratio of Set2 CD to histone dimers (40 µM CD and 32 µM histone dimers) are shown. Residues that could not be assigned are shown as gray bars. A subset of signals, 57-115, are shown, as we were able to obtain a comprehensive assignment only for AWS of the Set2 CD. **(D)** As in (C), but with H2A/H2B dimer.

To investigate direct Set2-histone interactions, we used NMR. In these experiments, we tested both the Set2 CD and Set2 AID for interactions with histone H3/H4 and H2A/H2B dimers. Our results showed that both Set2 domains interacted with both classes of histone dimers (Figure 5B-D, Supplementary Figure S7). This result supports earlier genetic results showing that histones H2A and H4 are required for H3K36 methylation (72,73). Furthermore, when we assayed the Set2-N90A mutant protein, it displayed weaker binding to both H2A/H2B and H3/H4 (Figure 5B-D, Supplementary Figure S7). Structural studies as Set2 N90 is located near residue R52 of the αN histone H3 helix, which is required for SETD2 activity *in vitro* (70) and H3K36me3 *in vivo* (74) (Figure 5A). Together, these results provide additional evidence that an important aspect of Set2 regulation is the dual role of specific residues in the Set2 CD in either conferring autoinhibition by interaction with Set2 AID or activity by interaction with its substrate, histone H3.

### Control of cryptic antisense transcription by Spt6 and Set2

A prominent phenotype caused by the loss of Set2 in yeast and SETD2 in mammalian cells is elevated levels of cryptic transcription in both the sense and antisense directions (9,10,30,75,76). If Spt6 is required for Set2 activity, then we would expect that *spt6-RR* and *set2Δ* would have similar effects on cryptic transcription. To test this, and to assay the degree of *spt6-RR* suppression by our strongest *SET2* suppressor, we measured the level of cryptic antisense transcripts by oligo-dT-primed RNA-seq. This was done in five strains: wild type, *set2Δ*, *spt6-RR*, *spt6-RR SET2-I89L/G260D*, and *SET2-I89L/G260D*. All measurements were performed in triplicate and the results were spike-in normalized (Supplementary Figure S8A,B).

Our results showed that the *set2Δ* and *spt6-RR* mutants had similar levels of derepressed antisense transcripts, with elevated levels measured for 284 and 296 genes, respectively, and with an overlap of 167 genes (p-value < 10^-50^) (Figure 6A-C, Supplementary Figure S8C, Supplementary Table S5). Furthermore, when we compared the *spt6-RR* mutant to the *spt6-RR SET2-I89L/G260D* mutant, the latter had a greatly decreased level of antisense transcripts, with only 86 genes with an elevated level of antisense, showing strong suppression of the antisense transcription phenotype by the *SET2-I89L/G260D* mutant (Figure 6C, Supplementary Figure S8D). These effects were also clear when looking at the single gene level (Figure 6D,E). Taken together, these results provide strong evidence that the Spt6 DLD is required for the same function as Set2, namely H3K36 methylation.

**Figure 6.**
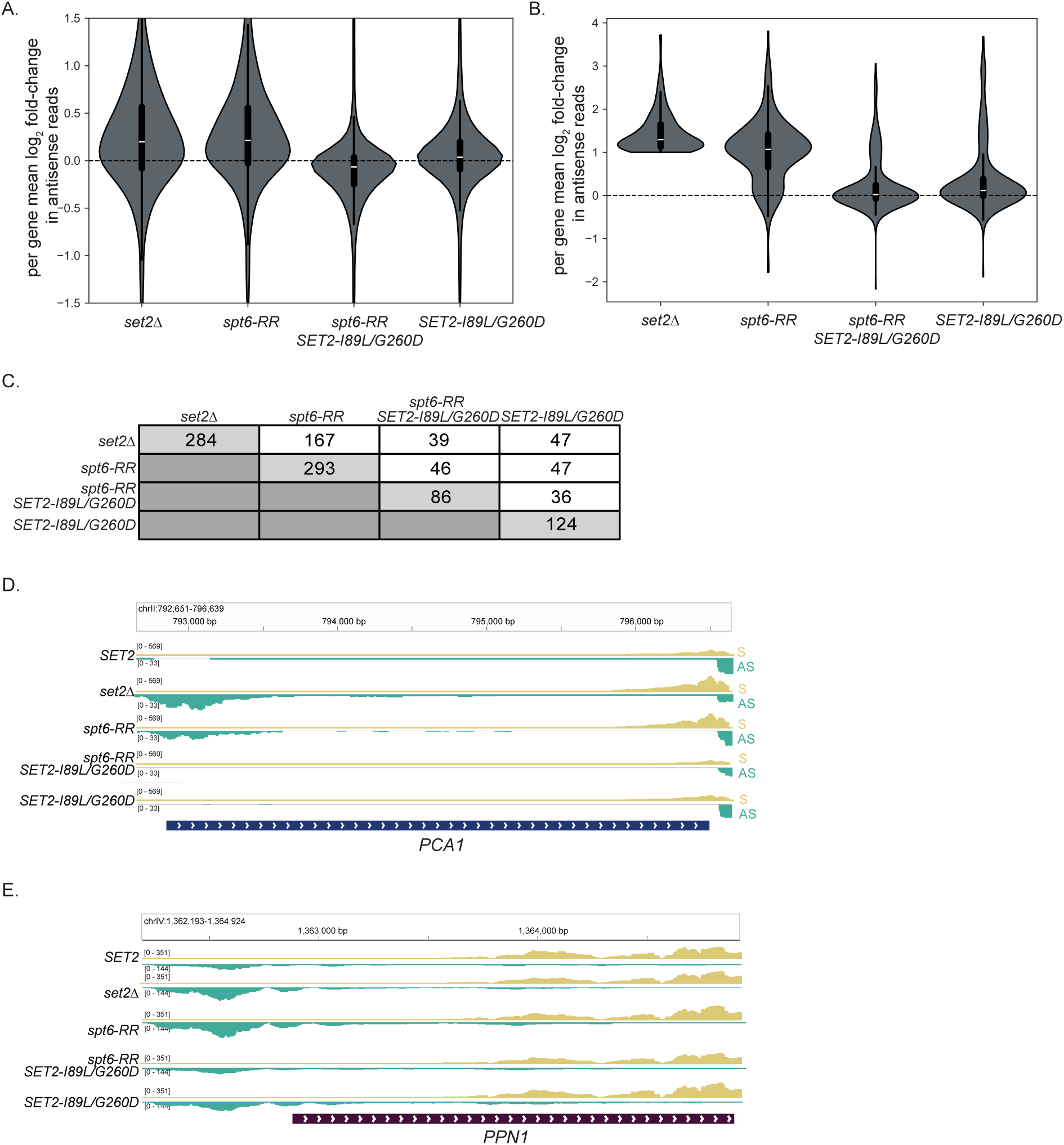
Analysis of antisense transcription. **(A)** Plots showing the log_2_ fold change of average antisense signal coverage in 3,087 non-overlapping protein-coding genes of the indicated strain relative to wild type. **(B)** Plots showing the average antisense signal coverage for each gene containing log_2_ fold-change >1 in *set2Δ* relative to wild type. Shown in (A) and (B) are the average of three replicates (see Supplementary Figure S8A,B). **(C)** Values represent the number of genes with a log_2_ fold-change >1 in average antisense signal coverage per gene relative to wild type. The numbers in the intersections show the number of genes in common between the two indicated mutants. **(D)** Genome browser view showing the distribution of 3’ antisense RNA-seq reads across *PCA1*, a gene previously shown to contain Set2 repressed antisense transcripts. **(E)** As in (D) but with *PPN1*.

### ChIP-seq analysis of *spt6-RR* and bypass of Spt6 by *set2* suppressors

To address the functional role of the Spt6-Set2 interaction *in vivo*, we performed ChIP-seq experiments. For these experiments, we assayed Set2 (using an HA epitope tag), Rpb1, H3K36me2, H3K36me3, and total H3 association across the yeast genome, in four different strains: wild type (*SPT6^+^ SET2-HA*), *spt6-RR SET2-HA*, *spt6-RR SET2-I89L/G260D-HA*, and *SET2-I89L/G260D-HA*. All assays were performed in triplicate and the results were spike-in normalized to exogenously added *S. pombe* chromatin (Supplementary Figure S9A). In the wild-type strain, our results showed strong association for Set2, H3K36me2, and H3K36me3 across gene bodies starting shortly after the TSS (Supplementary Figure S9B). When we compared the mutant strains to wild type, our results showed strong effects genome-wide for both H3K36me2 and H3K36me3: loss of both modifications in *spt6-RR*, with essentially complete suppression in the *spt6-RR SET2-I89L/G260D* mutant (Figure 7A,B, Supplementary Figure S9B-D). In the *SET2-I89L/G260D* mutant (with wild type *SPT6^+^)*, we observed elevated levels of H3K36me3 and decreased levels of H3K36me2 over gene bodies, similar to what was previously observed for *SET2-H366N*, a hyperactive Set2 AID mutant (Figure 7A,B) (28). We also observed H3K36me2 over the 5’ regions of genes, indicating that a hyperactive Set2 methylates H3K36 in locations not observed in wild type (Figure 7B, Supplementary Figure S9B).

**Figure 7.**
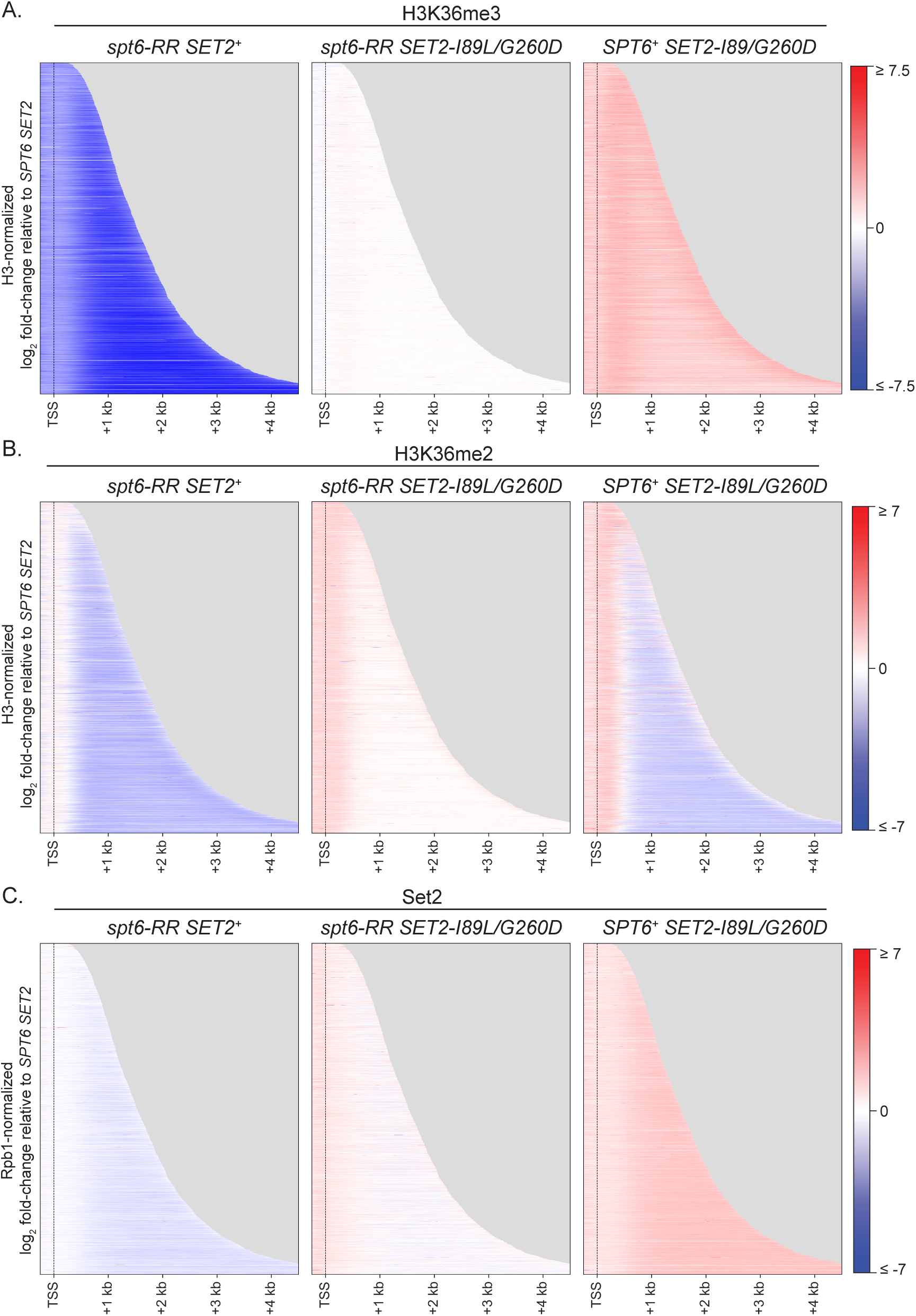
*SET2-I89L/G260D* restores H3K36 methylation genome-wide. Heatmaps of ChIP-seq results, showing the fold-change of **(A)** H3K36me3, **(B)** H3K36me2, or **(C)** Set2 relative to wild type between the indicated strains *spt6-RR, spt6-RR SET2-I89L/G260D*, or *SET2-I89L/G260D*. (A) and (B) are normalized to H3, and (C) is normalized to Rpb1. Shown are 3,087 non-overlapping protein-coding genes ordered by length.

The *spt6-RR* mutant provided an opportunity to test whether Spt6-Set2 interactions are required for Set2 recruitment *in vivo.* From our ChIP-seq data, we observed that there was a modestly reduced level of Set2 associated with chromatin in the *spt6-RR* mutant compared to wild type (Figure 7C), while Set2 levels were close to wild type levels in the *spt6-RR SET2-I89L/G260D* mutant, consistent with suppression, and modestly elevated above wild type levels in the *SET2-I89L/G260D* mutant. We note that there are approximately wild-type levels of Set2 protein in these strains (Figure 3E), ruling out altered Set2 abundance as underlying the ChIP-seq results. These results reveal a modest role for Spt6 in Set2 recruitment but suggest that Spt6 is primarily required for activation of Set2 in order to achieve the correct level of H3K36 methylation.

## DISCUSSION

Our work has provided several new insights into the regulation of the H3K36 methyltransferase Set2 by the histone chaperone Spt6. First, we have shown that the Spt6 DLD is required for detectable H3K36 methylation by Set2 *in vivo,* based on both western and ChIP-seq experiments. The strong phenotypic similarity between Spt6 DLD mutants and the complete loss of Set2 suggests that the primary function of the Spt6 DLD is to enable H3K36 methylation by Set2. This is consistent with the observation that the DLD is not conserved in the related bacterial protein Tex, even though the two flanking Spt6 domains (HhH and S1) are conserved (79–81). Our results extend recent structural and biochemical studies (36–38) with respect to the regulation of Set2 by Spt6.

Our results have also provided novel information regarding the mechanism for Set2 autoinhibition. First, we identified specific residues in the Set2 catalytic domain (CD) that are required for Set2 autoinhibition. Most of these amino acids are located along a predicted interface with the Set2 AID, directly opposite previously-identified Set2 AID mutants of the same phenotype (28), providing strong genetic evidence that a Set2 CD-AID direct interaction confers Set2 autoinhibition. We went on to use protein NMR and hydrogen-deuterium exchange to demonstrate a direct Set2 CD-AID interaction, and showed that this interaction is dependent upon amino acids identified by our mutants. Together, our results support a model in which Set2 exists in two conformations *in vivo*: an autoinhibited state, mediated by direct contacts between the CD and AID, and an activated state that requires binding to the Spt6 DLD and allows Set2 CD-histone H3 interactions.

While genetic, biochemical, and structural data all support that the binding of the Spt6 DLD to the Set2 SBR is required for overcoming Set2 autoinhibition, the mechanism by which the Spt6 DLD releases Set2 autoinhibition remains an open question. Based on structural predictions, it is unlikely that the DLD-SBR binding sterically blocks Set2 CD-AID interactions (Figure 2A), suggesting that the mechanism requires the participation of other proteins, such as histone H3. We unexpectedly observed that a four amino acid change in the SBR, *set2-4E*, was able to partially bypass the requirement for the Spt6 DLD, providing evidence that the SBR is normally important for autoinhibition. As we also observed that *set2-4E* caused a partial loss of H3K36 methylation in a wild type *SPT6^+^* background, we propose that this region normally serves two roles: in Set2 autoinhibition in the absence of Spt6 and in Set2 activation when Set2 SBR and Spt6 DLD are bound to each other.

Previous studies showed that the Set2 AID domain adopts a TND domain fold (amino acids 365-450). This domain has been identified in many eukaryotic transcription elongation factors (82) and it has been shown to interact with an unstructured TND-interacting motif (TIM) (82).

However, in contrast to the other TND-containing elongation factors, there is no evidence that this region of the Set2 AID recognizes a TIM and, consistent with this, the Set2 CD is not TIM-like in either structure or amino acid composition. Thus, the demonstration of the CD-AID interaction shows that TNDs have a broader range of interaction than was previously recognized.

Our analysis of Spt6 identified the HhH domain as essential for H3K36 methylation, similar to the Spt6 DLD. Indeed, the initial studies that demonstrated a requirement for Spt6 in H3K36 methylation all analyzed *spt6-1004*, an HhH deletion (4,27,30,31). However, there is no evidence for a direct Spt6 HhH-Set2 interaction based on AlphaFold 3 or structural studies (36–38). Furthermore, the *spt6ΔHhH* mutant is more pleiotropic than *spt6ΔDLD,* suggesting a broader role for the HhH domain than for the DLD. Thus, our data, combined with previously published structural data, suggest that the DLD is the only Spt6 domain directly interacting with Set2. As the HhH domain is directly adjacent to the DLD, the most likely explanation for the loss of H3K36 methylation in *spt6ΔHhH* is that the HhH domain is required for the correct position of the DLD in Spt6 relative to Set2.

Our ChIP-seq results showed that the level of chromatin-associated Set2 is partially dependent upon Spt6, as we observed a modestly decreased level of Set2 on chromatin in an *spt6-RR* mutant compared to wild type. This result, combined with earlier analysis (27), suggests that multiple Set2 domains contribute to Set2 recruitment to chromatin. The decreased level that we observed in *spt6-RR* is most likely caused by the autoinhibited conformation of Set2 rather than the lost interaction with Spt6 as there were near wild type levels of chromatin-associated Set2 in the *spt6-RR SET2-I89L/G260D* mutant, where Set2-Spt6 interactions would be equally impaired. These results are consistent with previous analysis of the *spt6-1004* mutant and its suppressor, *SET2-H366N* (28). While these effects on chromatin-associated Set2 were modest in the mutants that we tested, the changes in H3K36me2 and H3K36me3 were drastic. Those results, combined with biochemical results that showed that Spt6 directly activates SETD2 *in vitro* (36,37), strongly suggest that the requirement for Spt6 in Set2 function occurs primarily at a post-recruitment step for Set2.

Why H3K36 methylation needs to be regulated remains an open area of investigation, but several results suggest that such regulation is important. In several yeast mutant backgrounds, H3K36 methylation is toxic, including in many histone chaperone mutants (83–86) as well as mutants for Bur1, a kinase that phosphorylates the Rpb1 CTD and the Spt5 CTR (6,87). The tight regulation of histone methylation appears to be generally important, as previous studies have demonstrated autoinhibition for the H3K4 methyltransferase Set1 (88) and the H3K9 methyltransferase Clr4 (89). For Set2, autoinhibition ensures that H3K36 methylation is deposited in the right context—on nucleosomes during transcription—and at the right level.

Possibly, dynamic regulation of Set2 is required during rapid cellular changes, such as during changes in nutrients or during stress conditions (90,91). In humans, where most histone methyltransferases have been implicated in cancer (92–95), regulation of H3K36 methylation may be critical during development and in different cell types.

## DATA AND CODE AVAILABILITY

The next-generation sequencing data (unprocessed read files and processed coverage files) can be accessed at GEO (https://www.ncbi.nlm.nih.gov/geo/) with accession numbers GSE331502 (RNA-seq) and GSE332555 (ChIP-seq). The proteomics data have been deposited to the ProteomeXchange Consortium via the PRIDE partner repository (96) with the dataset identifier PXD078353. The scripts used to analyze next-generation sequencing data and an explanation of how spike-in normalization factors were calculated can be accessed at https://github.com/winston-lab.

## SUPPLEMENTARY DATA

Supplementary Data are available at NAR Online.

## ACKNOWLEDGMENTS

We thank Karen Arndt and Rajaraman Gopalakrishnan for helpful comments on the manuscript. We are grateful to Milan Fábry, Pavel Srb, Veronika Krejčiříková, Barbora Konečná and Magdaléna Hořejší for support with NMR sample preparation and data acquisition. We also thank František Filandr for support with mass spectrometry experiments.

## AUTHOR CONTRIBUTIONS

A.R.E. and F.W. wrote the manuscript, with input from all authors. NMR experiments and NMR data analysis were performed by V.L. and V.V. HDX experiments and data analysis were performed by T.N. and V.V. All other experiments were performed by A.R.E. Analyses of and data representation of RNA-seq and ChIP-seq were performed by J.L.W.

