## Supplementary Figures and Tables for "Regulation of the histone H3K36 methyltransferase Set2 by the histone chaperone Spt6"

Supplementary Figure S1

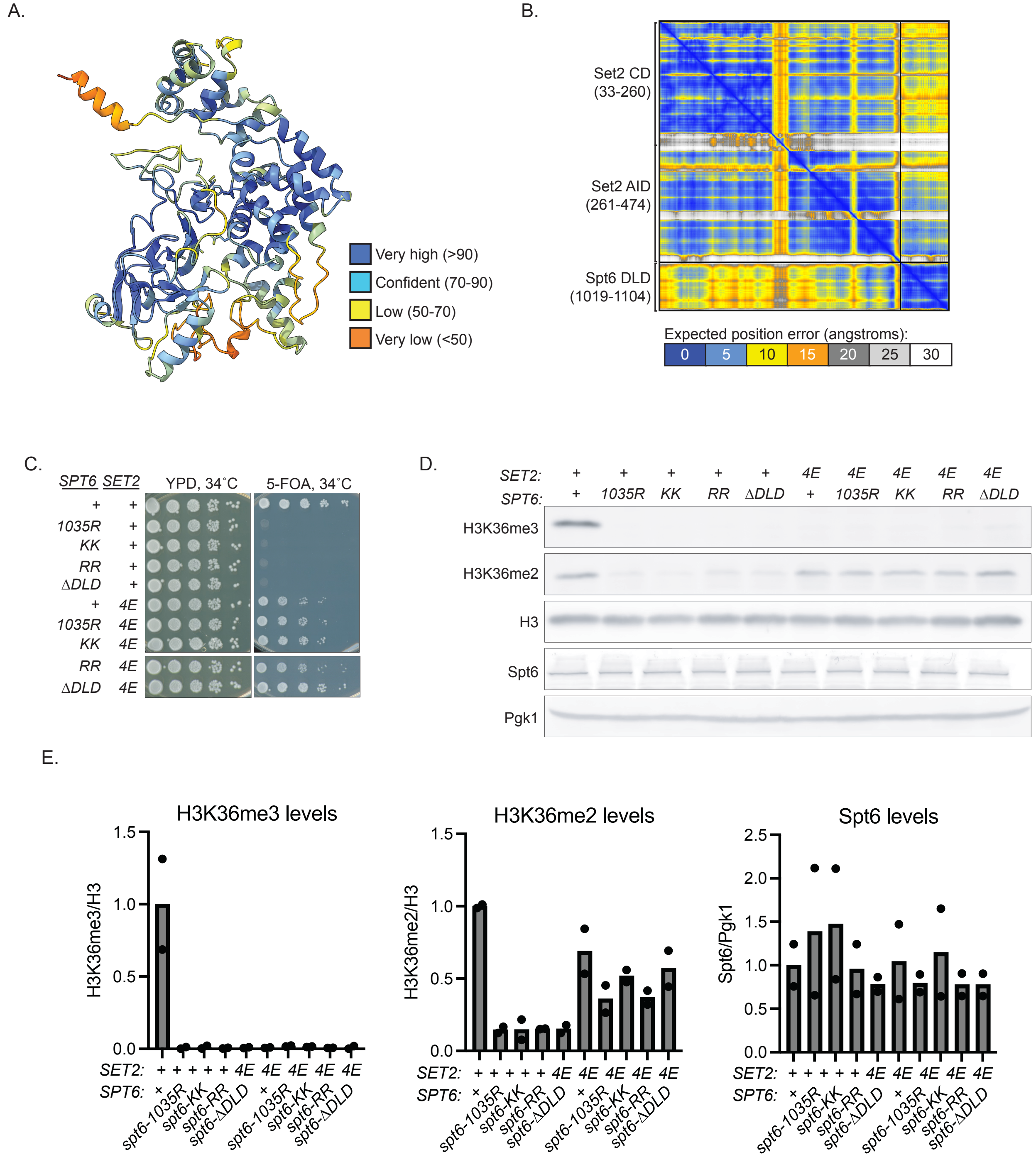

**Supplementary Figure S1. Mutant phenotypes of the *set2-4E* mutant.**

(A) AlphaFold3 model predicting an interaction between Set2 CD-AID (33-474) and Spt6 DLD (1019-1104). The model is colored by pLDDT scores which measure the per-residue confidence of the predicted model.

(B) Predicted Alignment Error plot for the AlphaFold 3 model in (B).

(C) Spot tests showing expression of the *FLO8-URA3* intragenic transcription reporter in *set2-4E* mutant. Plate images were taken after two days of incubation at 34˚C.

(D) Representative western blot analysis of H3K36 methylation of the mutants in (A).

(E) Quantification of three independent western blots in (B). Shown is the average of two independent replicates.

Supplementary Figure S2

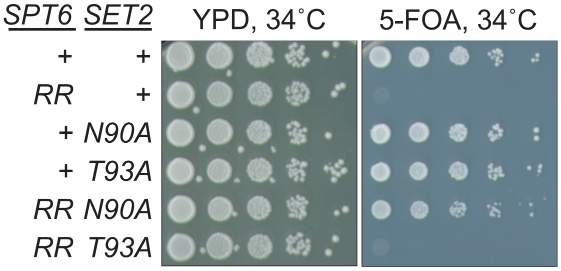


**Supplementary Figure S2. Analysis of the ability of *Set2-T93A* and *-N90A* to suppress the requirement for Spt6 for H3K36 methylation.**

Spot tests showing expression of the *FLO8-URA3* intragenic transcription reporter in the *set2-N90A* and *set2-T93A* mutants with *SPT6*$^{+}$ or *spt6-RR*. Plate images were taken after two days of incubation at 34˚C.

Supplementary Figure S3

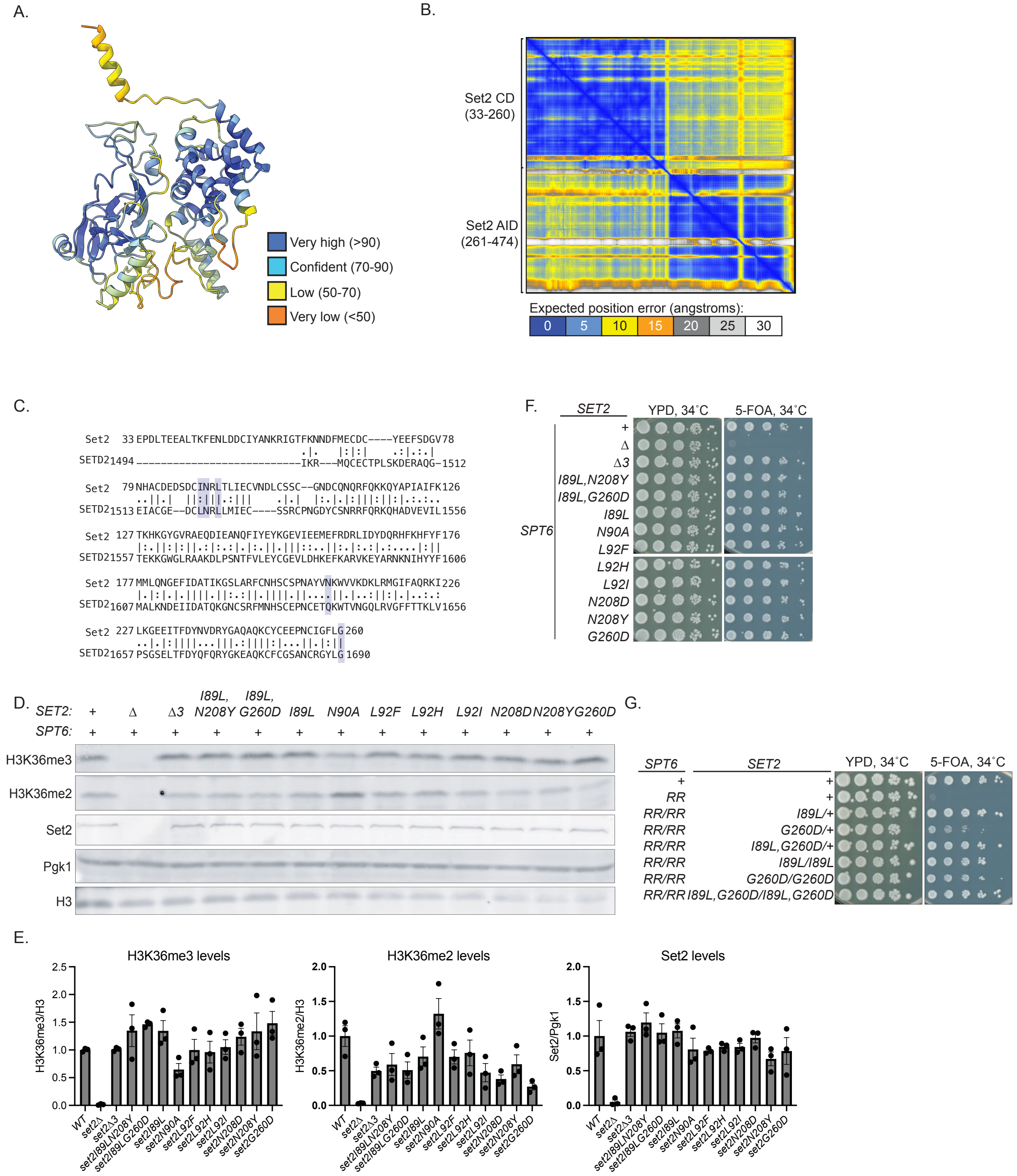

**Supplementary Figure S3. Measuring the effect of Set2 suppressor mutations on H3K36me2 and H3K36me3 levels.**

(A) AlphaFold3 model of Set2 CD and Set2 AID (33-474). The model is colored by pLDDT scores which measure the per-residue confidence of the predicted model.

(B) Predicted Alignment Error plot for the AlphaFold 3 model in (F).

(C) Alignment of Set2 CD and SETD2 CD. Highlighted in lavender are the residues that were mutated in the suppressor selection presented in this paper.

(D) Representative western blot analysis of each Set2 suppressor mutation on H3K36 methylation in a wild-type *SPT6*$^{+}$ background.

(E) Quantification of three independent western blots in (A). Each dot represents a single replicate. The error bars indicate the standard error of the mean.

(F) Spot tests analysis of the level of intragenic transcription in the Set2 suppressor mutants. Plates were incubated for two days at 34˚C.

(G) Spot test analysis of the level of intragenic cryptic transcription in diploids that are either homozygous or heterozygous for the indicated mutation in *SET2*. Plate images shown are after two days of incubation at 34˚C.

Supplementary Figure S4

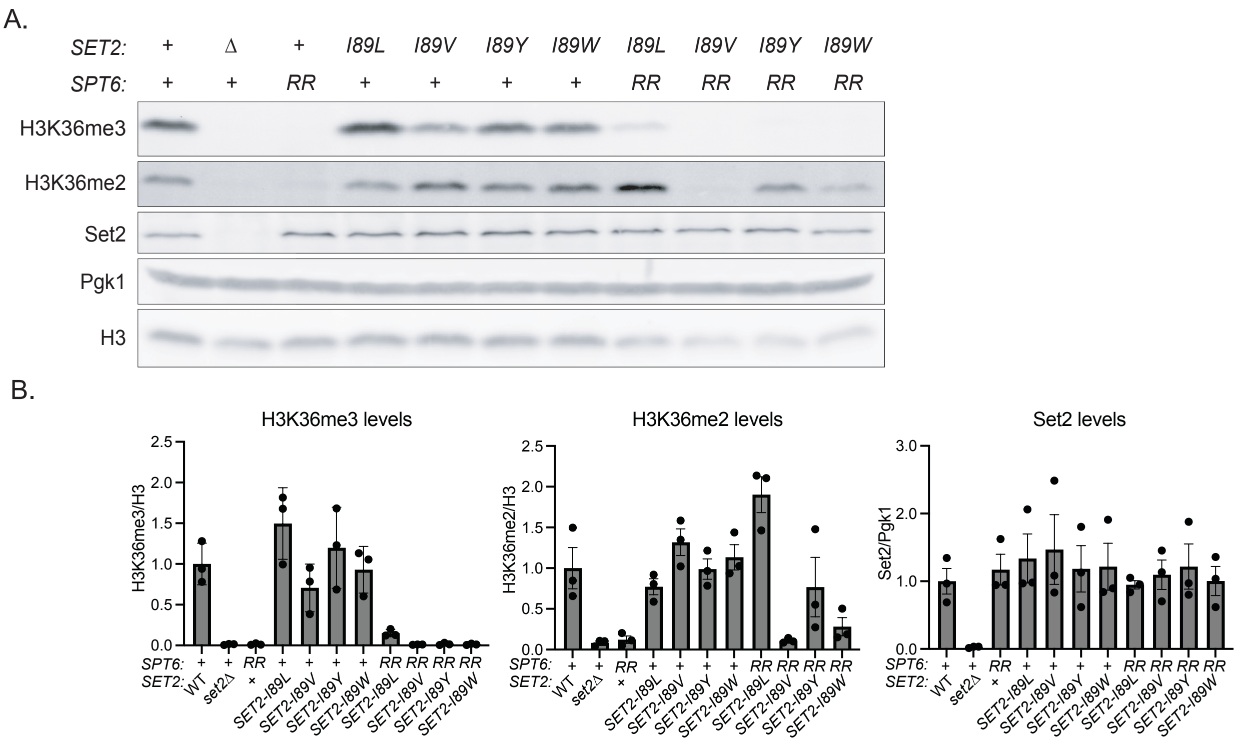


**Supplementary Figure S4. Allele specificity for the suppression of *spt6-RR* at Set2 I89*.***

(A) Representative western blots of *set2-I89* mutants in *SPT6*$^{+}$ and *spt6-RR* backgrounds.

(B) Quantification of three independent western blot experiments in (A). Each dot represents a single replicate. The error bars indicate the standard error of the mean.

### Supplementary Figure S5

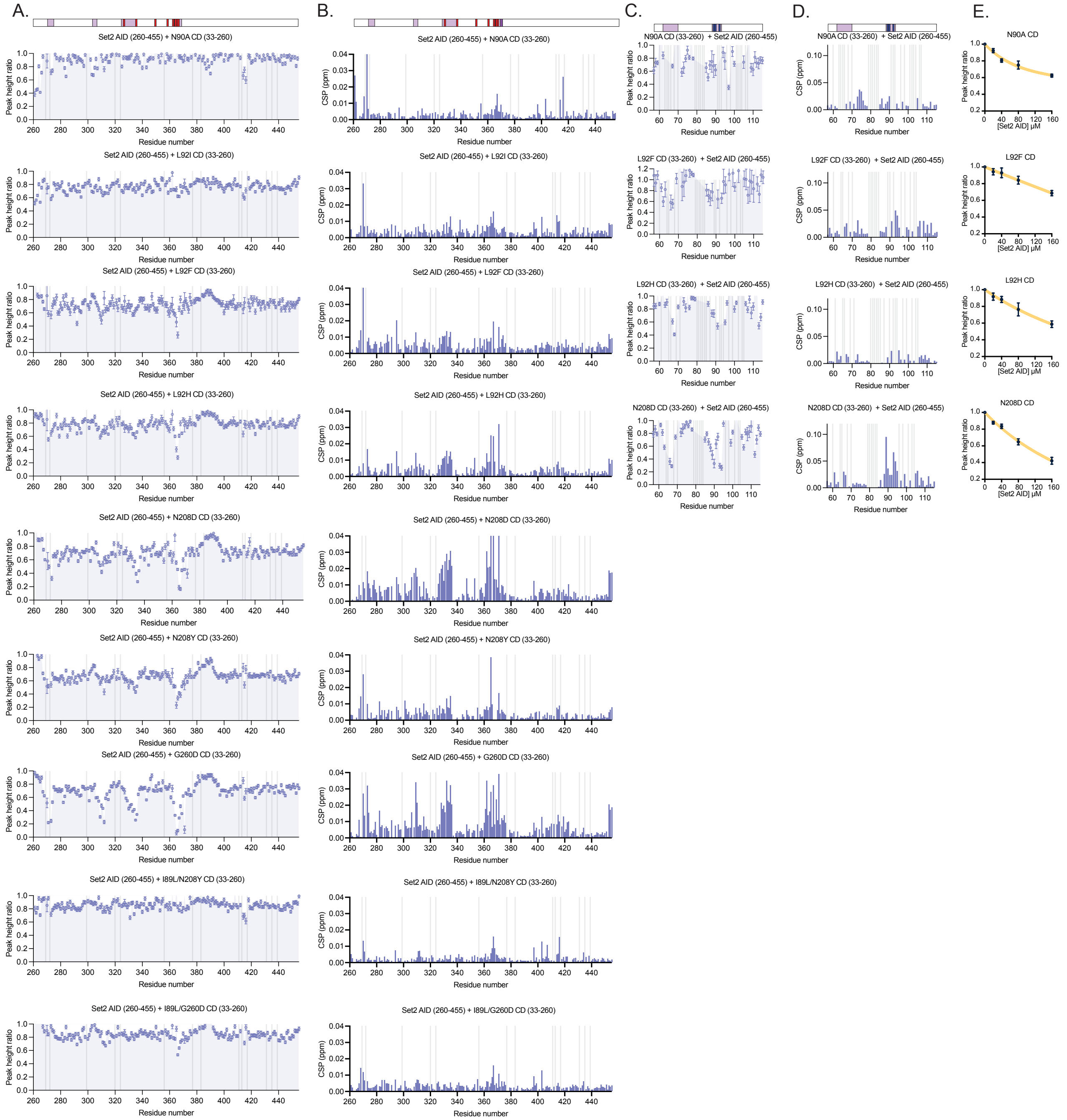

**Supplementary Figure S5. Further analysis of Set2 CD-AID interactions.**

(A) Plot of the NMR signal intensity ratios obtained for Set2 AID upon titration with the indicated Set2 CD mutant. Residues that could not be assigned are indicated as gray bars. The data obtained using four-fold molar excess of Set2 CD are shown. A lower peak height ratio indicates evidence of binding.

(B) Quantification of backbone amide chemical shift perturbations (CSPs) in the Set2 AID induced upon addition of Set2 CD (33-260). The data obtained using four-fold molar excess of Set2 CD are shown. Residues that could not be assigned are shown as gray bars. The color bar above the plots shows the location within the Set2 AID of previously identified suppressor mutations (red) and AlphaFold-predicted AID-CD interface residues (purple).

(C) As in (A) but obtained for the indicated Set2 CD mutant upon titration with Set2 AID. A subset of signals, 57-115, are shown, as we were able to obtain a comprehensive assignment only for AWS of the Set2 CD.

(D) As in (B) but for CSPs in the Set2 CD induced upon addition of Set2 AID (260-455). A subset of signals, 57-115, are shown, as we were able to obtain a comprehensive assignment only for AWS of the Set2 CD.

(E) Normalized and averaged intensities of 10 most perturbed peaks of the indicated mutant Set2 CD upon titration of the Set2 AID (260-455).

Supplementary Figure S6

A.

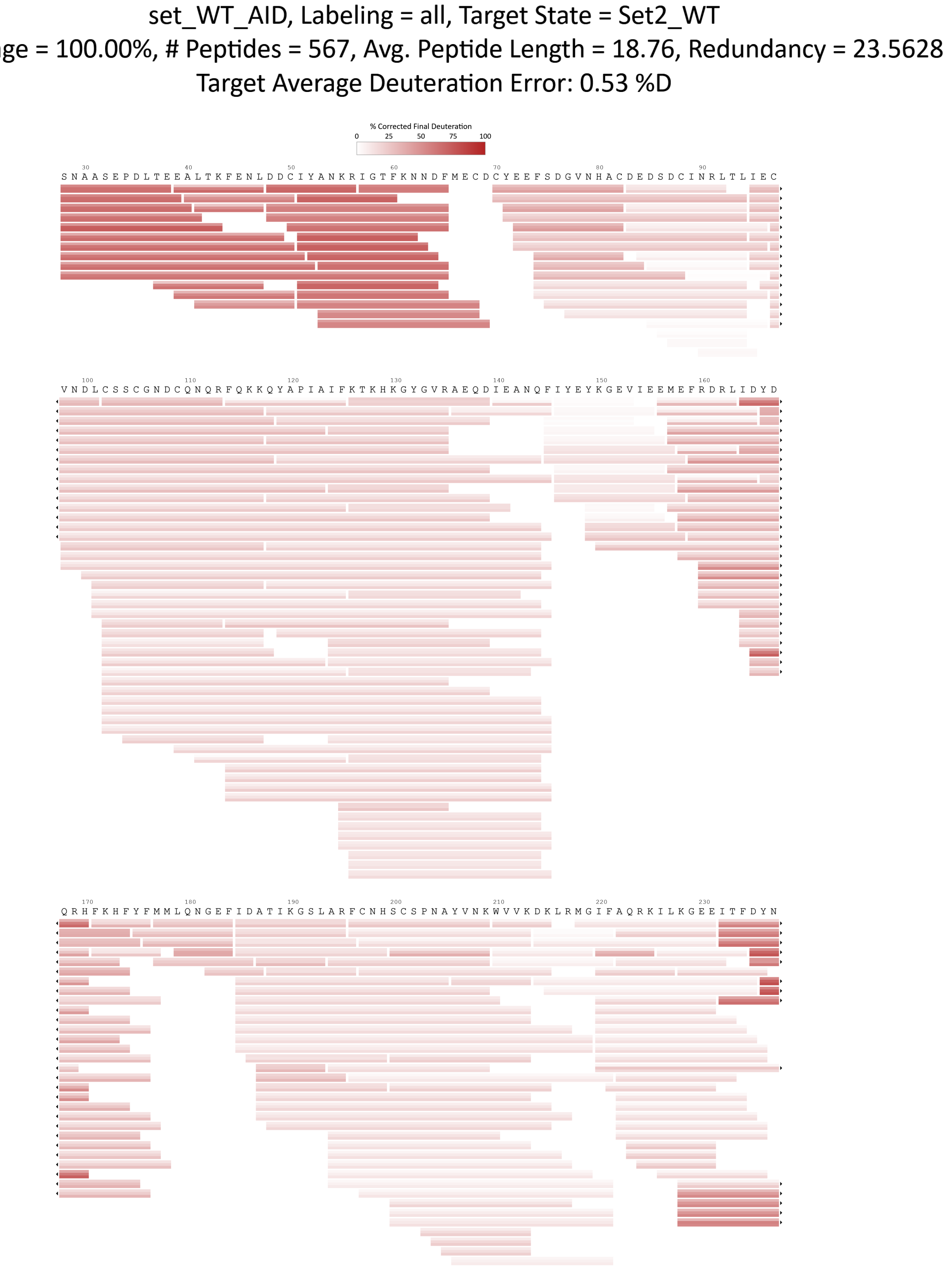

A. (continued)

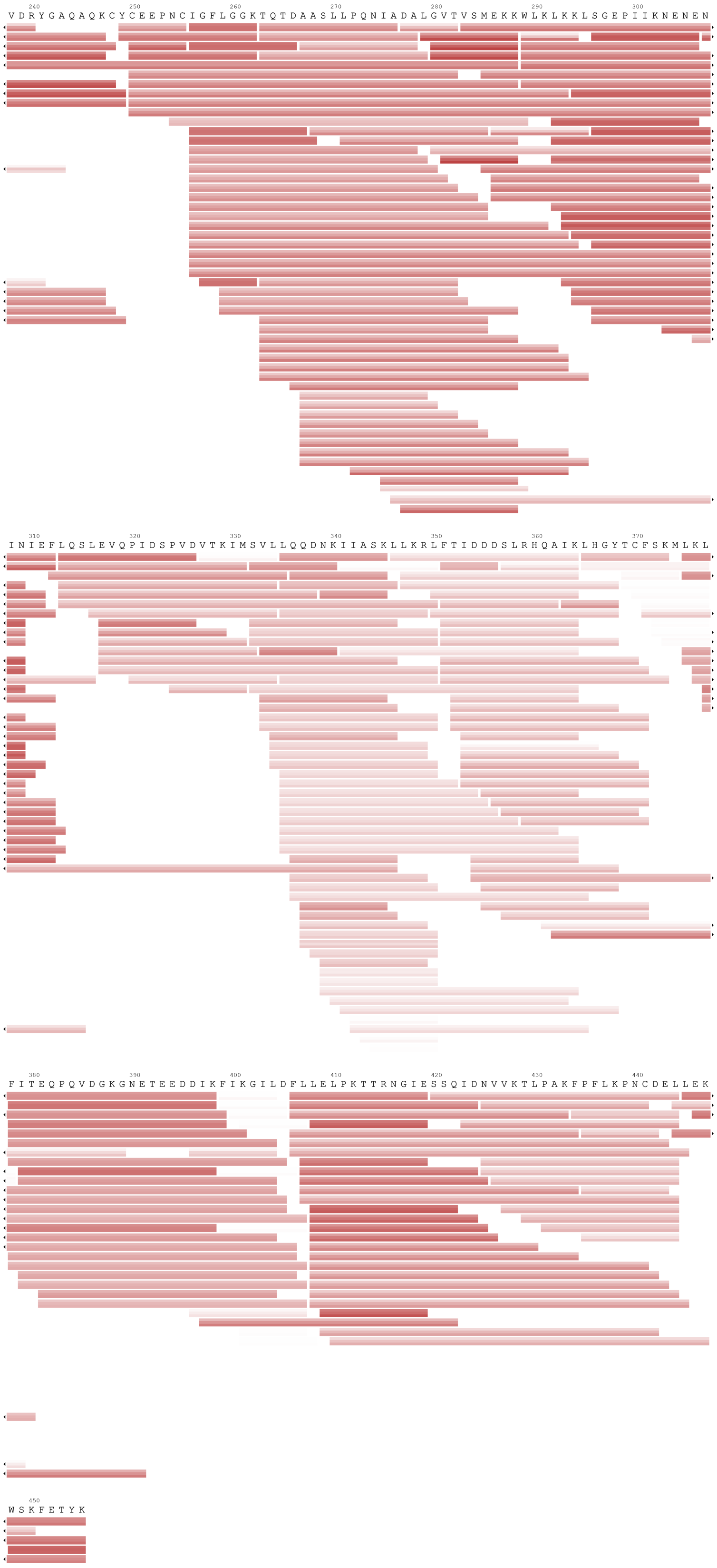

240 250 260 270 280 290 300
V D R Y G A Q A Q K C Y C E E P N C I G F L G G K T Q T D A A S L L P Q N I A D A L G V T V S M E K K W L K L K K L S G E P I I K N E N E N
310 320 330 340 350 360 370
I N I E F L Q S L E V Q P I D S P V D V T K I M S V L L Q Q D N K I I A S K L L K R L F T I D D D S L R H Q A I K L H G Y T C F S K M L K L
380 390 400 410 420 430 440
F I T E Q P Q V D G K G N E T E E D D I K F I K G I L D F L L E L P K T T R N G I E S S Q I D N V V K T L P A K F P F L K P N C D E L L E K
450
W S K F E T Y K

B.

set_MUT_G260D, Labeling = all, Target State = Set2_G260D
Coverage = 100.00%, # Peptides = 514, Avg. Peptide Length = 18.32, Redundancy = 20.8261
Target Average Deuteration Error: 0.62 %D

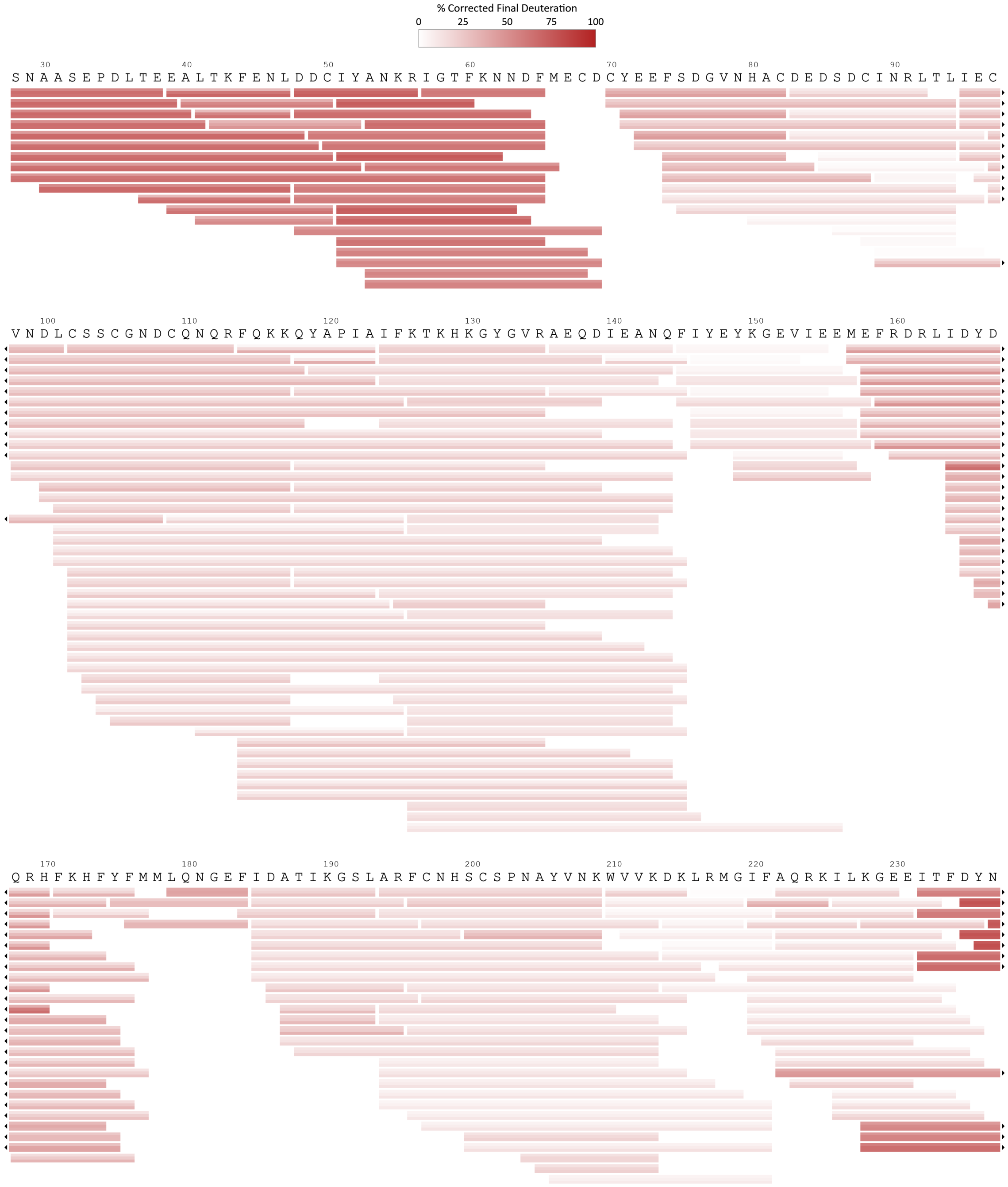

B. (continued)

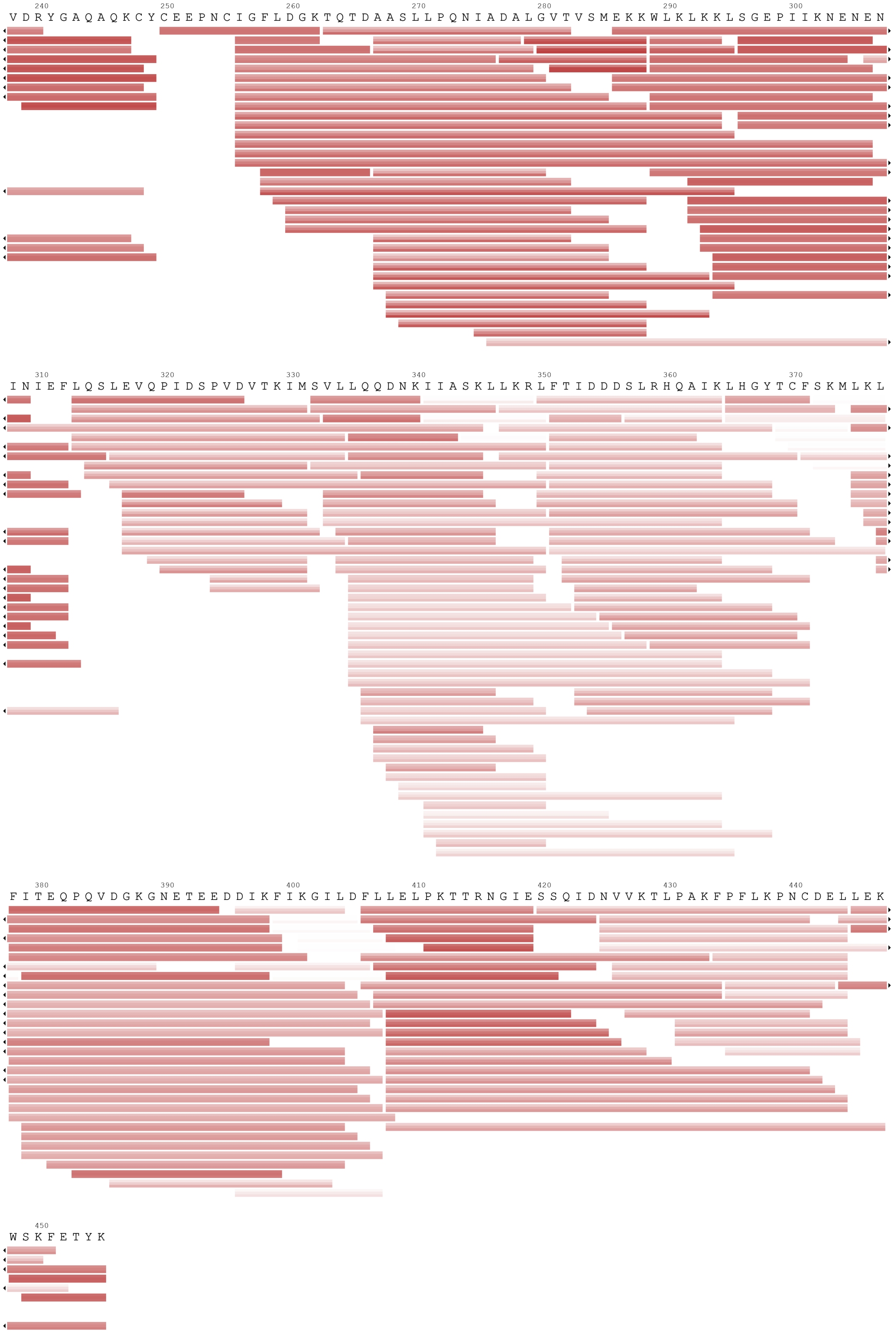

240 250 260 270 280 290 300
V D R Y G A Q A Q K C Y C E E P N C I G F L D G K T Q T D A A S L L P Q N I A D A L G V T V S M E K K W L K L K K L S G E P I I K N E N E N
310 320 330 340 350 360 370
I N I E F L Q S L E V Q P I D S P V D V T K I M S V L L Q Q D N K I I A S K L L K R L F T I D D D S L R H Q A I K L H G Y T C F S K M L K L
380 390 400 410 420 430 440
F I T E Q P Q V D G K G N E T E E D D I K F I K G I L D F L L E L P K T T R N G I E S S Q I D N V V K T L P A K F P F L K P N C D E L L E K
450
W S K F E T Y K

C.

set_MUT_I89L/G260D, Labeling = all, Target State = Set2_I89L_G260D
Coverage = 100.00%, # Peptides = 348, Avg. Peptide Length = 19.98, Redundancy = 15.4638
Target Average Deuteration Error: 0.50 %D

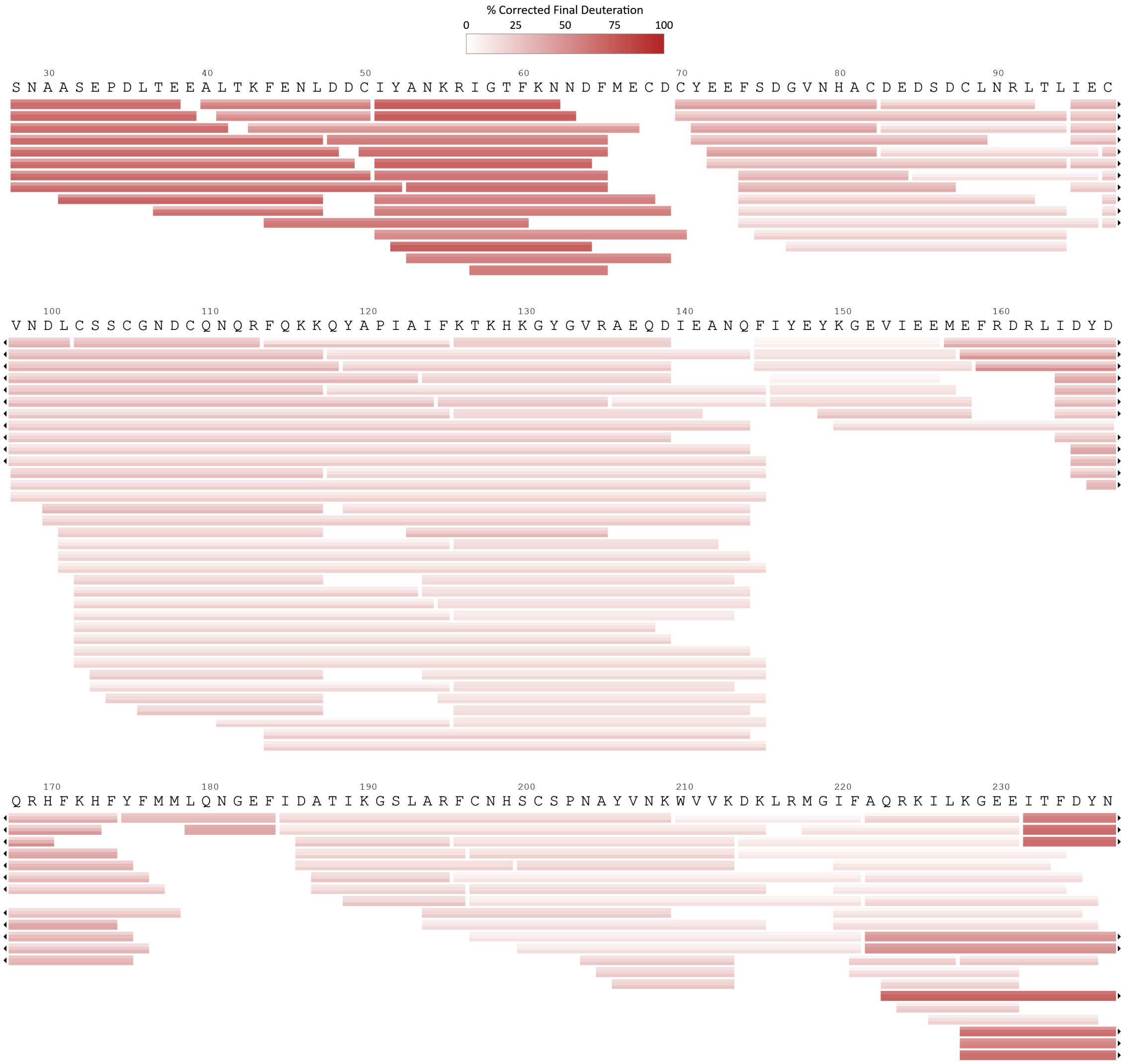

C. (continued)

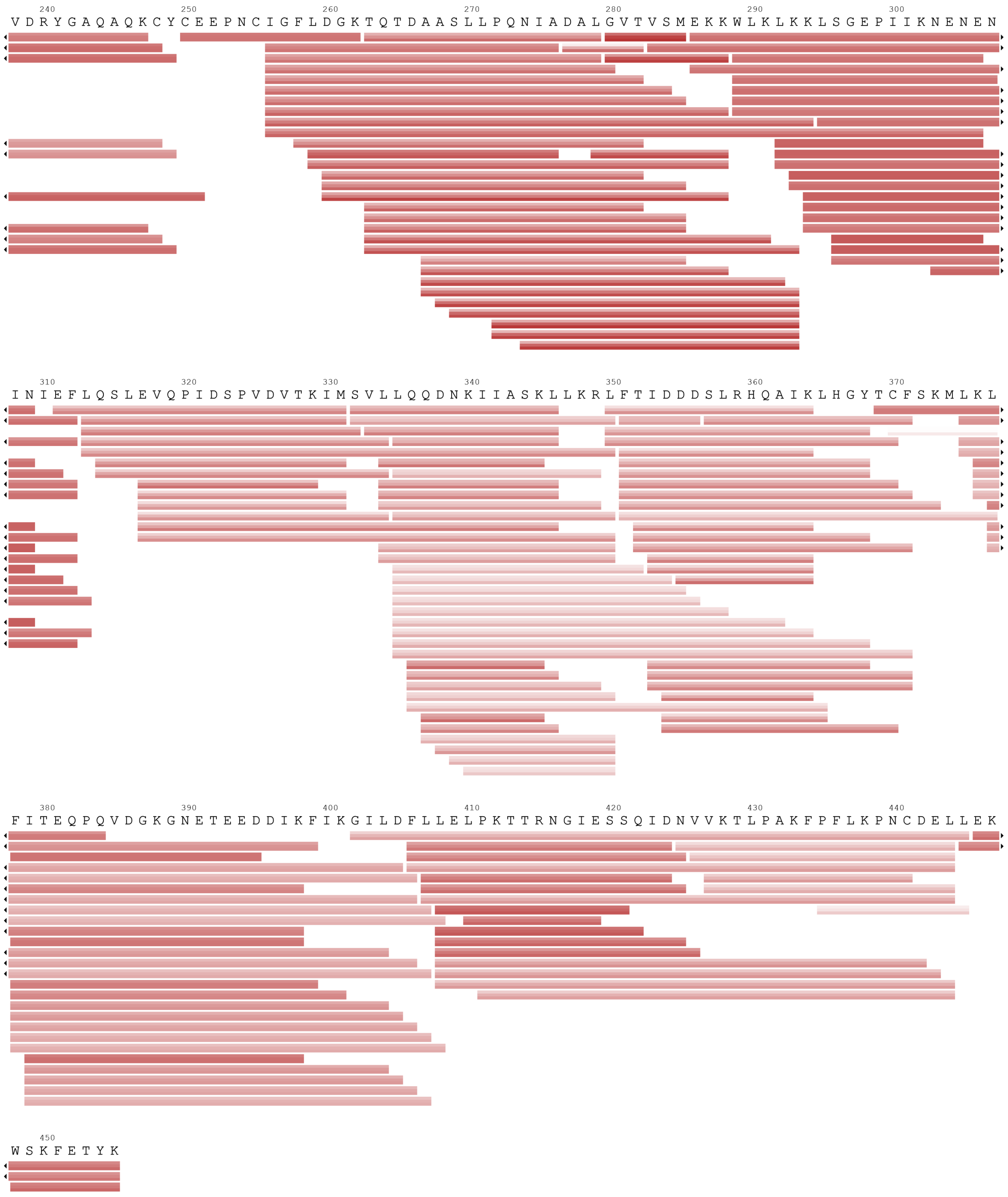

240 250 260 270 280 290 300
V D R Y G A Q A Q K C Y C E E P N C I G F L D G K T Q T D A A S L L P Q N I A D A L G V T V S M E K K W L K L K K L S G E P I I K N E N E N
310 320 330 340 350 360 370
I N I E F L Q S L E V Q P I D S P V D V T K I M S V L L Q Q D N K I I A S K L L K R L F T I D D D S L R H Q A I K L H G Y T C F S K M L K L
380 390 400 410 420 430 440
F I T E Q P Q V D G K G N E T E E D D I K F I K G I L D F L L E L P K T T R N G I E S S Q I D N V V K T L P A K F P F L K P N C D E L L E K
450
W S K F E T Y K

**Supplementary Figure S6.** HDX-MS peptide coverage maps for Set2 variants.
(A-C) Each bar represents an individual peptide (WT Set2 in (A), G260D-containing Set2 in (B), I89L/G260D-containing Set2 in (C)) that was valid at all three labeling time points and in at least three out of four replicates. The red shading indicates relative fractional deuterium uptake corrected for the maximum achievable deuteration level of 90%. Each peptide bar is divided into three horizontal sections corresponding to the labeling time points, shown from top to bottom: 15 s, 2 min and 20 min.

Supplementary Figure S7

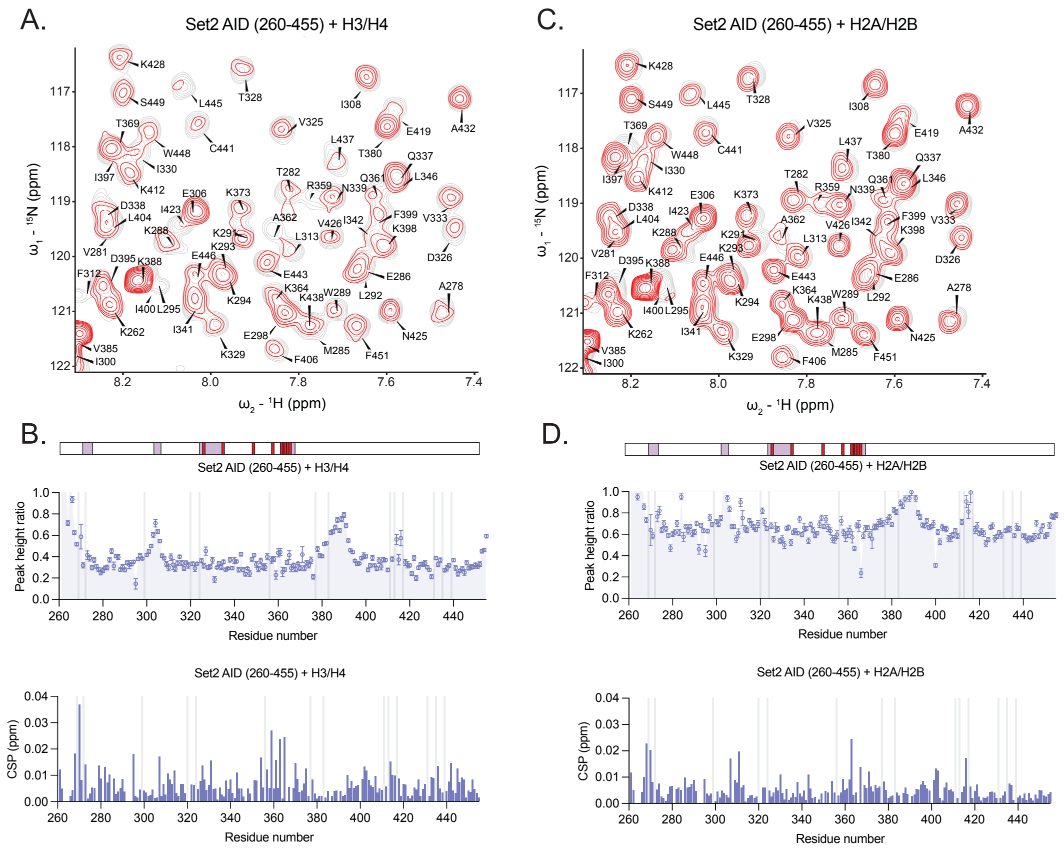


**Supplementary Figure S7. Analysis of Set2 AID interactions with core histones using NMR.**

(A) Overlaid 2D $^{15}N/^{1}H$ HSQC spectra obtained for free (gray) or histone H3/H4 (red) $^{13}C/^{15}N$ labeled Set2 AID.

(B) The color bar above the plots shows the location within the Set2 AID of previously identified suppressor mutations (red) and AlphaFold-predicted AID-CD interface residues (purple). The top plot of the NMR signal intensity ratios obtained for Set2 AID upon titration with the histone H3/H4 dimer. Residues that could not be assigned are indicated as gray bars. The data obtained using four-fold molar excess of histones are shown. A lower peak height ratio indicates evidence of binding. The bottom plot shows the quantification of backbone amide chemical shift perturbations (CSPs) in the Set2 AID induced upon addition of the histone H3/H4 dimer. A higher chemical shift perturbation indicates evidence of binding. The data obtained using four-fold molar excess of histone are shown. Residues that could not be assigned are shown as gray bars.

(C) As in (A) but with H2A/H2B dimer.

(D) As in (B) but with H2A/H2B dimer.

Supplementary Figure S8

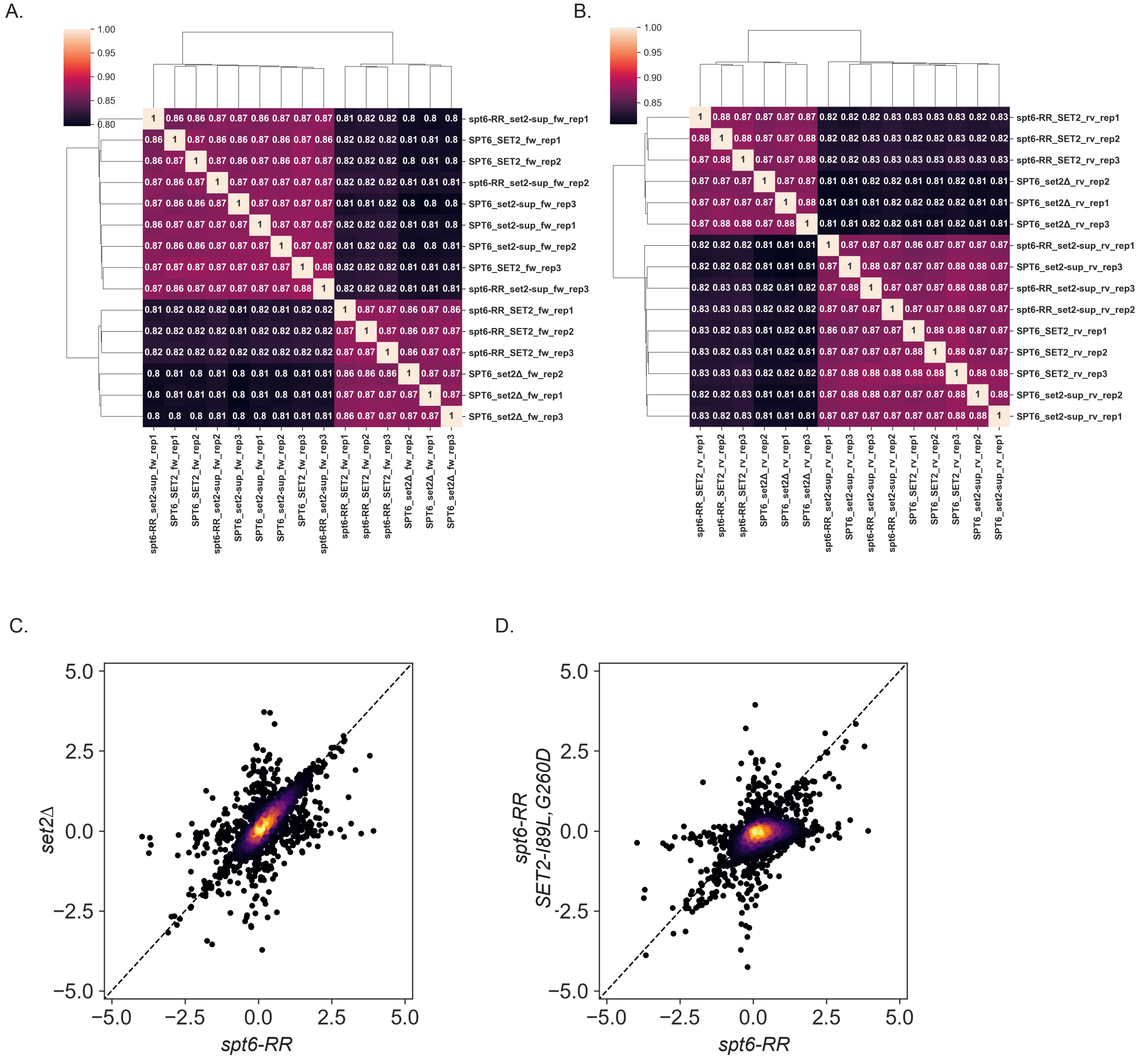


**Supplementary Figure S8. Analysis of antisense transcription.**

(A) Clustered heatmaps showing the Spearman correlation of the forward strand RNA-seq reads from 3,087 non-overlapping protein-coding genes between all conditions and biological replicates.

(B) As in (A), but for reverse strand reads.

(C) A scatterplot comparing the $\log_2$ fold change in antisense transcript abundance in the *set2Δ* and *spt6-RR* mutants compared to wild type. Each point represents one of 3,087 non-overlapping protein-coding genes. Points are colored using a kernel density estimate to illustrate the density of the data.

(D) A similar scatterplot comparing *spt6-RR SET2-I89L/G260D* to *spt6-RR*.

#### Supplementary Figure S9

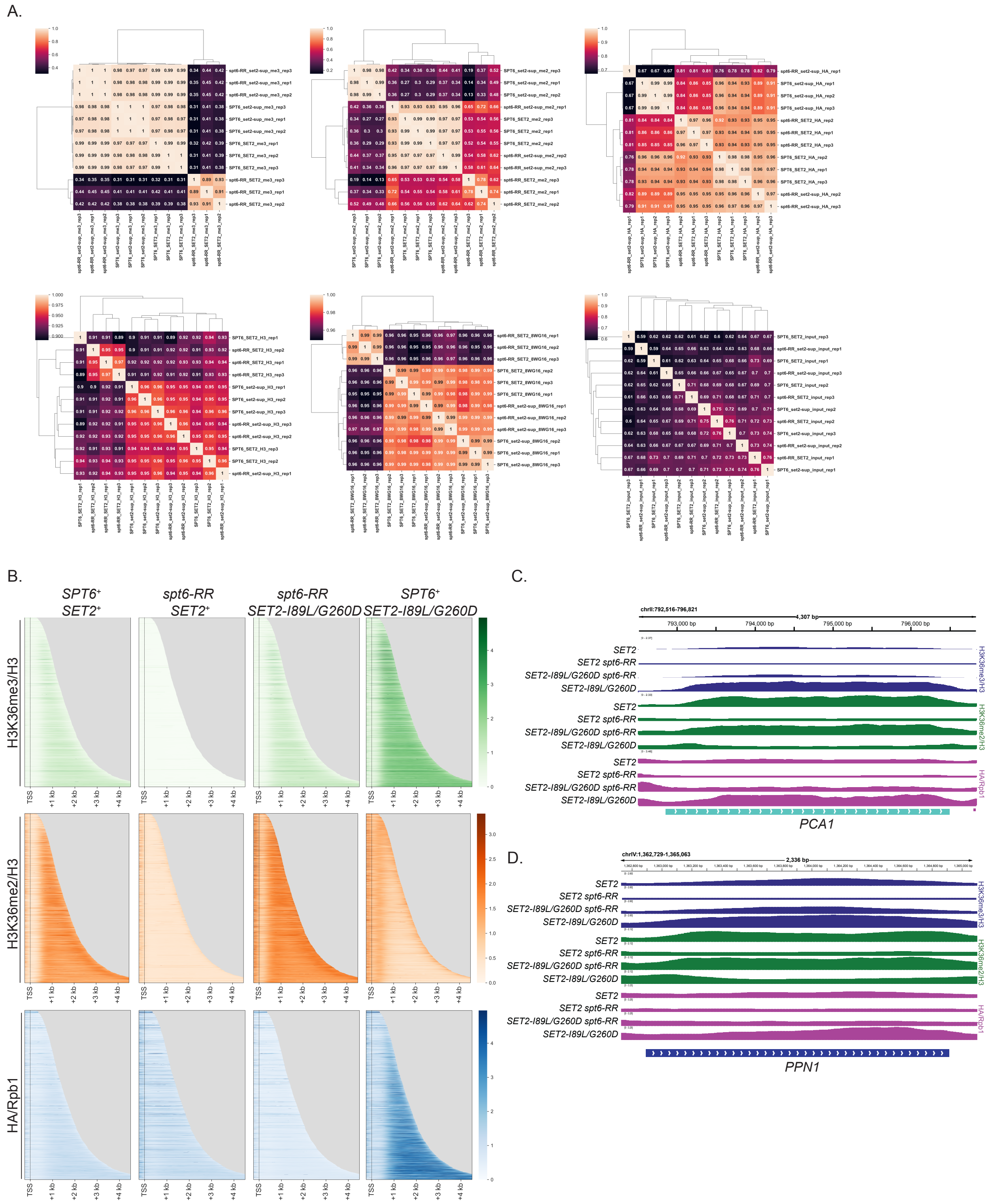

**Supplementary Figure S9.** ***SET2-I89L/G260D*** **restores H3K36 methylation genome-wide.**

(A) Clustered heatmaps showing the Spearman correlation of the indicated ChIP-seq enrichment between all conditions and biological replicates.

(B) ChIP-seq results, showing the H3- or Rpb1-normalized levels of H3K36me3, H3K36me2, or Set2 relative in the indicated strains *spt6-RR, spt6-RR SET2-I89L/G260D*, or *SET2-I89L/G260D*.

(C) Genome browser profile showing the distribution of H3K36me3, H3K36me2 and Set2 over *PCA1.* Values are the average of three biological replicates (see S8D).

(D) As in (A), but over *PPN1.*

**Supplementary Table S1.** Yeast strains

| Strain | Genotype | Species |
|---|---|---|
| FWP566 | SET2-3xHA-NatMX | S. pombe |
| FY87 | MATα lys2-128δ ura3-52 leu2Δ1 | S. cerevisiae |
| FY3137 | MATα his3Δ200 lys2-128δ ura3-52 leu2Δ1 FLO8-URA3 spt6-D1035R/E1038R set2-T93A | S. cerevisiae |
| FY3199 | MATα his3Δ200 lys2-128δ ura3-52 leu2Δ1 FLO8-URA3 set2Δ::NatMX | S. cerevisiae |
| FY3544 | MATα LEU2::TIR1, lys2-128δ, ura3-52, trp1Δ63, SPT6-V5-AID:KanMX | S. cerevisiae |
| FY3588 | MATα his3Δ200 lys2-128δ ura3-52 leu2Δ1 FLO8-URA3 set2-T93A | S. cerevisiae |
| FY3589 | MATα his3Δ200 lys2-128δ ura3-52 leu2Δ1 FLO8-URA3 spt6-D1035R/E1038R | S. cerevisiae |
| FY3590 | MATα his3Δ200 lys2-128δ ura3-52 FLO8-URA3 leu2Δ1 set2Δ::KanMX spt6-D1035R/E1038R | S. cerevisiae |
| FY3591 | MATα his3Δ200 lys2-128δ ura3-52 leu2Δ1 FLO8-URA3 spt6-D1035R/E1038R SET2-I89L | S. cerevisiae |
| FY3592 | MAT**a** his3Δ200 lys2-128δ ura3-52 leu2Δ1 FLO8-URA3 spt6-D1035R/E1038R SET2-N90A | S. cerevisiae |
| FY3593 | MATα his3Δ200 leu2Δ1 lys2-128δ ura3-52 FLO8-URA3 SET2-N90A | S. cerevisiae |
| FY3594 | MAT**a** his3Δ200 lys2-128δ leu2Δ1 ura3-52 FLO8-URA3 SET2-G260D | S. cerevisiae |
| FY3595 | MAT**a** his3Δ200 lys2-128δ ura3-52 FLO8-URA3 SET2-L92H | S. cerevisiae |
| FY3596 | MATα his3Δ200 leu2Δ1 lys2-128δ ura3-52 FLO8-URA3 SET2-I89L | S. cerevisiae |
| FY3597 | MATα his3Δ200 lys2-128δ ura3-52 FLO8-URA3 SET2-L92I | S. cerevisiae |
| FY3598 | MAT**a** his3Δ200 lys2-128δ ura3-52 FLO8-URA3 spt6-D1035R/E1038R SET2Δ3 | S. cerevisiae |
| FY3599 | MATα his3Δ200 lys2-128δ leu2Δ1 ura3-52 FLO8-URA3 spt6-D1035R/E1038R SET2-N208Y | S. cerevisiae |
| FY3600 | MAT**a** his3Δ200 lys2-128δ ura3-52 FLO8-URA3 spt6-D1035R/E1038R SET2-I89L | S. cerevisiae |
| FY3601 | MAT**a** his3Δ200 lys2-128δ ura3-52 FLO8-URA3 SET2-L92F | S. cerevisiae |
| FY3602 | MAT**a** his3Δ200 lys2-128δ ura3-52 FLO8-URA3 SET2-N208Y | S. cerevisiae |
| FY3603 | MATα his3Δ200 lys2-128δ ura3-52 leu2Δ1 FLO8-URA3 spt6-D1035R/E1038R SET2-I89V | S. cerevisiae |
| FY3604 | MATα his3Δ200 lys2-128δ ura3-52 leu2Δ1 FLO8-URA3 spt6-D1035R/E1038R SET2-I89Y | S. cerevisiae |
| FY3605 | MATα his3Δ200 leu2Δ1 lys2-128δ ura3-52 FLO8-URA3 SET2-N208D | S. cerevisiae |
| FY3606 | MATα his3Δ200 lys2-128δ ura3-52 leu2Δ1 FLO8-URA3 spt6-D1035R/E1038R SET2-I89W | S. cerevisiae |

| | | |
|---|---|---|
| FY3607 | *MATα his3Δ200 lys2-128δ ura3-52 FLO8-URA3 spt6-D1035R/E1038R SET2-G260D* | *S. cerevisiae* |
| FY3608 | *MAT**a** his3Δ200 lys2-128δ ura3-52 FLO8-URA3 spt6-D1035R/E1038R SET2-L92F* | *S. cerevisiae* |
| FY3609 | *MAT**a** his3Δ200 lys2-128δ ura3-52 FLO8-URA3 spt6-D1035R/E1038R SET2-N208D* | *S. cerevisiae* |
| FY3610 | *MATα his3Δ200 lys2-128δ ura3-52 FLO8-URA3 spt6-D1035R/E1038R SET2-L92H* | *S. cerevisiae* |
| FY3611 | *MAT**a** his3Δ200 lys2-128δ ura3-52 FLO8-URA3 spt6-D1035R/E1038R SET2-L92I* | *S. cerevisiae* |
| FY3612 | *MATα his3Δ200 lys2-128δ ura3-52 FLO8-URA3 spt6-D1035R/E1038R SET2-I89L/G260D* | *S. cerevisiae* |
| FY3613 | *MATα his3Δ200 lys2-128δ ura3-52 leu2Δ1 FLO8-URA3 spt6-D1035R/E1038R SET2-3xHA-KanMX* | *S. cerevisiae* |
| FY3614 | *MATα his3Δ200 lys2-128δ ura3-52 leu2Δ1 FLO8-URA3 SET2-3xHA-KanMX* | *S. cerevisiae* |
| FY3615 | *MAT**a** his3Δ200 lys2-128δ ura3-52 leu2Δ1 FLO8-URA3 spt6-D1035R/E1038R SET2-I89L/N208Y* | *S. cerevisiae* |
| FY3616 | *MATα his3Δ200 lys2-128δ ura3-52 FLO8-URA3 SET2-I89L/G260D* | *S. cerevisiae* |
| FY3617 | *MAT**a** his3Δ200 lys2-128δ ura3-52 FLO8-URA3 SET2-I89L/N208Y* | *S. cerevisiae* |
| FY3618 | *MATα his3Δ200 lys2-128δ leu2Δ1 ura3-52 FLO8-URA3 spt6-D1035R/E1038R* | *S. cerevisiae* |
| FY3619 | *MATα his3Δ200 lys2-128δ ura3-52 leu2Δ1 FLO8-URA3 spt6-D1035R/E1038R SET2-I89L/G260D-3xHA-KanMX* | *S. cerevisiae* |
| FY3620 | *MATα his3Δ200 lys2-128δ ura3-52 leu2Δ1 FLO8-URA3 SET2-I89L/G260D-3xHA-KanMX* | *S. cerevisiae* |
| FY3621 | *MATα his3Δ200 lys2-128δ ura3-52 FLO8-URA3 SET2-I89W* | *S. cerevisiae* |
| FY3622 | *MATα lys2-128δ ura3-52 SET2-I89L/G260D spt6-YW* | *S. cerevisiae* |
| FY3623 | *MAT**a** lys2-128δ ura3-52 SET2-I89L/G260D spt6-YW* | *S. cerevisiae* |
| FY3624 | *MAT**a** his3Δ200 lys2-128δ ura3-52 FLO8-URA3 SET2-I89V* | *S. cerevisiae* |
| FY3625 | *MAT**a** his3Δ200 lys2-128δ leu2Δ1 ura3-52 FLO8-URA3 SET2-I89Y* | *S. cerevisiae* |
| FY3626 | *MAT**a**/α his3Δ200/his3Δ200 lys2-128δ / lys2-128δ FLO8-URA3/FLO8-URA3 ura3-52/ura3-52 SET2-I89L/SET2+ spt6-D1035R/E1038R/spt6-D1035R/E1038R* | *S. cerevisiae* |
| FY3627 | *MAT**a**/α his3Δ200/his3Δ200 lys2-128δ / lys2-128δ FLO8-URA3/FLO8-URA3 ura3-52/ura3-52 SET2-G260D/SET2+ spt6-D1035R/E1038R/spt6-D1035R/E1038R* | *S. cerevisiae* |
| FY3628 | *MAT**a**/α his3Δ200/his3Δ200 lys2-128δ / lys2-128δ FLO8-URA3/FLO8-URA3 ura3-52/ura3-52 SET2-I89L/G260D/SET2+ spt6-D1035R/E1038R/spt6-D1035R/E1038R* | *S. cerevisiae* |

| | | |
|---|---|---|
| FY3629 | *MATα his3Δ200 lys2-128δ ura3-52 leu2Δ1 FLO8-URA3 spt6-D1035R/E1038R set2-T93A* | *S. cerevisiae* |
| FY3630 | *MAT**a**/α his3Δ200/his3Δ200 lys2-128δ / lys2-128δ FLO8-URA3/FLO8-URA3 ura3-52/ura3-52 SET2-I89L/SET2-I89L spt6-D1035R/E1038R/spt6-D1035R/E1038R* | *S. cerevisiae* |
| FY3631 | *MAT**a**/α his3Δ200/his3Δ200 lys2-128δ / lys2-128δ FLO8-URA3/FLO8-URA3 ura3-52/ura3-52 SET2-G260D/SET2-G260D spt6-D1035R/E1038R/spt6-D1035R/E1038R* | *S. cerevisiae* |
| FY3632 | *MAT**a**/α his3Δ200/his3Δ200 lys2-128δ / lys2-128δ FLO8-URA3/FLO8-URA3 ura3-52/ura3-52 SET2-I89L/G260D/SET2-I89L/G260D spt6-D1035R/E1038R/spt6-D1035R/E1038R* | *S. cerevisiae* |

**Supplementary Table S2.** Oligos

| Allele created | gRNA | Repair Template Fw | Repair Template Rv |
|---|---|---|---|
| *spt6ΔHtH* | GACGGTAAGCATATATAAAA | ATTCCAGAAAGATATCAAGAATTAAGAGCAGGTATTACTGACGGTGGTTCTGGTGGTTCT | TCTGATTTTTGAAGTATTCAGTGACAATAGGATCATCGATAGAACCACCAGAACCACCGT |
| *spt6ΔHhH* | TGGTTCATTTAGCCTTTGAA | CTTGAAACCGCTTTCGTTGATATTGTCAACCTGGTAAGTGGTGGTTCTGGTGGTTCTGAA | TAGTTGATCATGTTCCAAATCTTCGTATTTTTGTCTTTTTTCAGAACCACCAGAACCACC |
| *spt6ΔDLD* | TCTAGTTTAGCTCTACGGTC | ACAAAAATACGAAGATTTGGAACATGATCAACTAGATAGCGGTGGTTCTGGTGGTTCTTT | TCACCTTGCAAAGGATGAAAGTCATTTCTCAATTCTTCAAAAGAACCACCAGAACCACCG |
| *spt6ΔS1* | GCAAATGAAATATACGAAAT | TTGCAAGGTGATGAAATTTTCCAAAGTTTGACTGGTGAGTCTGGTGGTTCTGGTGGTTCT | CTTCCAGTTCTTGTTTCAAGTCCCAAATGGAAGGATCTTTAGAACCACCAGAACCACCAG |
| *spt6ΔtSH2* | GAAGATATTCTACAATGATC | GAGGAGAGGAAATTGATGATGGCAGAAGCCCGTGCAAAGAGAGGTGGTTCTGGTGGTTCT | GCATCTAACGGTAGTTGTTCATTCTATTCTTACTACTGTTAGAACCACCAGAACCACCTC |
| *spt6-D1035R/E1038R* | ATCGTATTCTAAAGCATCAG | TAGATAGCACTAGAATTCATCCAGAAGACTACCATTTGGCCACCAAGGTTGCTGCTAGAG | TGTTCTTCTTTTTCGGCAATAGTATCAGGATCGTATCTTAAAGCTCTAGCAGCAACCTTG |
| *spt6-D1035K/E1038K* | ATCGTATTCTAAAGCATCAG | TAGATAGCACTAGAATTCATCCAGAAGACTACCATTTGGCCACCAAGGTTGCTGCTAAAG | TGTTCTTCTTTTTCGGCAATAGTATCAGGATCGTATTTTAAAGCTTTAGCAGCAACCTTG |
| *spt6-D1035A/E1038A* | ATCGTATTCTAAAGCATCAG | TAGATAGCACTAGAATTCATCCAGAAGACTACCATTTGGCCACCAAGGTTGCTGCTGCTG | GTTCTTCTTTTTCGGCAATAGTATCAGGATCGTAGGCTAAAGCAGCAGCAGCAACCTTG |
| *spt6ΔDLD* | ATCGTATTCTAAAGCATCAG | ACAAAAATACGAAGATTTGGAACATGATCAACTAGATAGCGGTGGTTCTGGTGGTTCTTT | TCACCTTGCAAAGGATGAAAGTCATTTCTCAATTCTTCAAAAGAACCACCAGAACCACCG |
| *spt6-D1035R* | ATCGTATTCTAAAGCATCAG | CCCTGTTCTTCTTTTTCGGCAATAGTATCAGGATCGTATTCTAAAGCTCTAGCAGCAACC | GCACTAGAATTCATCCAGAAGACTACCATTTGGCCACCAAGGTTGCTGCTAGAGCTTTAG |
| *spt6-D1038R* | ATCGTATTCTAAAGCATCAG | TCACTCATAGTCCCCTGTTCTTCTTTTTCGGCAATAGTATCAGGATCGTATCTTAAAGCA | CCAGAAGACTACCATTTGGCCACCAAGGTTGCTGCTGATGCTTTAAGATACGATCCTGAT |
| *SET2-I89L* | CGTCAACCATGCCTGTGACG | AGAATTCTCAGATGGCGTCAACCATGCCTGTGACG | TGAACACAAATCGTTGACACATTCTATCAACGTAAGT |

| | | | |
|---|---|---|---|
| | | AAGATTCTGACTGTTTAAATAGACT | CTATTTAAACAGTCAGAATCTTC |
| *SET2-N90A* | CGTCAACCATGCCTGTGACG | GACACATTCTATCAACGTAAGTCTAGCGATACAGTCAGAATCTTCGTCACAGGCATGGTT | GATTTCATGGAATGTGACTGTTATGAAGAATTCTCAGATGGCGTCAACCATGCCTGTGAC |
| *SET2-L92F* | CGTCAACCATGCCTGTGACG | CTCAGATGGCGTCAACCATGCCTGTGACGAAGATTCTGACTGTATCAATAGATTCACGTT | ACCACAAGATGAACACAAATCGTTGACACATTCTATCAACGTGAATCTATTGATACAGTC |
| *SET2-L92H* | CGTCAACCATGCCTGTGACG | CGTCAACCATGCCTGTGACGAAGATTCTGACTGTATCAATAGACATACGTTGATAGAATG | ATCATTACCACAAGATGAACACAAATCGTTGACACATTCTATCAACGTATGTCTATTGAT |
| *SET2-L92I* | CGTCAACCATGCCTGTGACG | CGTCAACCATGCCTGTGACGAAGATTCTGACTGTATCAATAGAATCACGTTGATAGAATG | TCATTACCACAAGATGAACACAAATCGTTGACACATTCTATCAACGTGATTCTATTGATA |
| *SET2-N208D* | TGTTAATAAATGGGTTGTTA | TGGCCAGATTTTGCAATCACTCTTGCAGCCCCAATGCATACGTTGATAAATGGGTTGTTA | TTCTTTGAGCAAATATTCCCATGCGTAGCTTATCTTTAACAACCCATTTATCAACGTATG |
| *SET2-N208Y* | TGTTAATAAATGGGTTGTTA | CGTTGGCCAGATTTTGCAATCACTCTTGCAGCCCCAATGCATACGTTTATAAATGGGTTG | TTTGAGCAAATATTCCCATGCGTAGCTTATCTTTAACAACCCATTTATAAACGTATGCAT |
| *SET2-G260D* | TGGTAAGACTCAAACAGATG | TGCTACTGTGAGGAGCCAAATTGTATTGGGTTTCTCGACGGTAAGACCCAAACAGATGCC | ACTCCCAAGGCATCAGCGATGTTCTGGGGCAATAAAGATGCGGCATCTGTTTGGGTCTTA |
| *SET2-I89L/N208Y* | CGTCAACCATGCCTGTGACG | AGAATTCTCAGATGGCGTCAACCATGCCTGTGACGAAGATTCTGACTGTTTAAATAGACT | TGAACACAAATCGTTGACACATTCTATCAACGTAAGTCTATTTAAACAGTCAGAATCTTC |
| *SET2-I89L/G260D* | CGTCAACCATGCCTGTGACG | AGAATTCTCAGATGGCGTCAACCATGCCTGTGACGAAGATTCTGACTGTTTAAATAGACT | TGAACACAAATCGTTGACACATTCTATCAACGTAAGTCTATTTAAACAGTCAGAATCTTC |
| *SET2-I89Y* | CGTCAACCATGCCTGTGACG | GTGACTGTTATGAAGAGTTTTCAGATGGCGTCAACCATGCCTGTGACGAAGATTCTGACT | AAATCGTTGACACATTCTATCAACGTAAGTCTATTGTAACAGTCAGAATCTTCGTCACAG |
| *SET2-I89W* | CGTCAACCATGCCTGTGACG | GTGACTGTTATGAAGAGTTTTCAGATGGCGTCAACCATGCCTGTGACGAAGATTCTGACT | AAATCGTTGACACATTCTATCAACGTAAGTCTATTCCAACAGTCAGAATCTTCGTCACAG |
| *SET2-I89V* | CGTCAACCATGCCTGTGACG | TTGACACATTCTATCAACGTAAGTCTATTGACACAGTCAGAATCTTCGTCACAGGCATGG | GACTGTTATGAAGAGTTTTCAGATGGCGTCAACCATGCCTGTGACGAAGATTCTGACTGT |

| **Plasmid name and description** | | **Primer Fw** | **Primer Rv** |
|---|---|---|---|
| bARE45 | pRS414 [spt6ΔIWS1-3xFLAG, TRP1, CEN/ARS] | ACACAGATTACTGGTCTA<br>TCGGGTGGTTCTGGTGG<br>TTCTGGTAACGACAACAA<br>TGAAGCT | AGCTTCATTGTTGTCGTT<br>ACCAGAACCACCAGAAC<br>CACCCGATAGACCAGTAA<br>TCTGTGT |
| bARE46 | pRS414 [spt6ΔYqgF-3xFLAG, TRP1, CEN/ARS] | AAGTTTATGACAAAATTAG<br>ACGGTGGTTCTGGTGGT<br>TCTTTGTTGGAATATGCTA<br>ATTTA | TAAATTAGCATATTCCAAC<br>AAAGAACCACCAGAACC<br>ACCGTCTAATTTTGTCATA<br>AACTT |
| bARE50 | pRS414 [spt6ΔHtH-3xFLAG, TRP1, CEN/ARS] | TTAAGAGCAGGTATTACT<br>GACGGTGGTTCTGGTGG<br>TTCTATCGATGATCCTATT<br>GTCACT | AGTGACAATAGGATCATC<br>GATAGAACCACCAGAACC<br>ACCGTCAGTAATACCTGC<br>TCTTAA |
| bARE44 | pRS414 [spt6ΔHhH-3xFLAG, TRP1, CEN/ARS] | TTGATATTGTCAACCTGG<br>TAAGTGGTGGTTCTGGTG<br>GTTCTGAAAAAAGACAAA<br>AATACG | CGTATTTTTGTCTTTTTTC<br>AGAACCACCAGAACCAC<br>CACTTACCAGGTTGACAA<br>TATCAA |
| bARE52 | pRS414 [spt6ΔDLD-3xFLAG, TRP1, CEN/ARS] | TGGAACATGATCAACTAG<br>ATAGCGGTGGTTCTGGTG<br>GTTCTTTTGAAGAATTGA<br>GAAATG | CATTTCTCAATTCTTCAAA<br>AGAACCACCAGAACCAC<br>CGCTATCTAGTTGATCAT<br>GTTCCA |
| bAR53 | pRS414 [spt6ΔS1-3xFLAG, TRP1, CEN/ARS] | CAAAGTTTGACTGGTGAG<br>TCTGGTGGTTCTGGTGG<br>TTCTAAAGATCCTTCCATT<br>TGGGAC | GTCCCAAATGGAAGGATC<br>TTTAGAACCACCAGAACC<br>ACCAGACTCACCAGTCA<br>AACTTTG |
| bARE51 | pRS414 [spt6ΔtSH2-3xFLAG, TRP1, CEN/ARS] | GCAGAAGCCCGTGCAAA<br>GAGAGGTGGTTCTGGTG<br>GTTCTAACAGTAGTAAGA<br>ATAGAATG | CATTCTATTCTTACTACTG<br>TTAGAACCACCAGAACCA<br>CCTCTCTTTGCACGGGC<br>TTCTGC |
| **Purpose of primer** | | **Primer Fw** | **Primer Rv** |
| PCR Mutagenesis upstream of Set2 AID | | AAAACTGCATAGTCGTGC<br>TGTCAAACCTTTCTCCTT<br>TCCTGGTTGTTGTTTTGC<br>GTGATC | TCCCAAGGCATCAGCGAT<br>GTTCTGGGGCAATAAAGA<br>TGCCGCATCTGTTTGAGT<br>CTTACC |
| Tagging Set2 with 3xHA for ChIP-seq | | ATCAACAAGGATGTCTTC<br>TCCTCCACCTTCAAC<br>ATCATCACGGATCCCCG<br>GGTTAATTAA | GAAAACGTGAAACAAGC<br>CCCAAATATGCATGT<br>CTGGTTAAGAATTCGAGC<br>TCGTTTAAAC |

**Supplementary Table S3**. Plasmids.

| Plasmid | Plasmid description | Host | Source | Purpose |
|---|---|---|---|---|
| bARE10 | pRS415 [*SET2,* LEU2, CEN/ARS] | DH5α | This study | PCR-mutagenesis screen |
| pRS414 | pRS414 [TRP1, CEN/ARS] | DH5α | PMID: 2659436 | Depletion-complementation |
| pJLW93 | pRS414 [SPT6-3xFLAG, TRP1, CEN/ARS] | DH5α | PMID: 40972526 | Depletion-complementation |
| pJLW95 | pRS414 [spt6-Δ2-238-3xFLAG, TRP1, CEN/ARS] | DH5α | PMID: 40972526 | Depletion-complementation |
| bARE45 | pRS414 [*spt6ΔSPN1*-3xFLAG, TRP1, CEN/ARS] | DH5α | This study | Depletion-complementation |
| bARE46 | pRS414 [*spt6ΔYqgF*-3xFLAG, TRP1, CEN/ARS] | DH5α | This study | Depletion-complementation |
| bARE50 | pRS414 [*spt6ΔHtH*-3xFLAG, TRP1, CEN/ARS] | DH5α | This study | Depletion-complementation |
| bARE44 | pRS414 [*spt6ΔHhH*-3xFLAG, TRP1, CEN/ARS] | DH5α | This study | Depletion-complementation |
| bARE52 | pRS414 [*spt6ΔDLD*-3xFLAG, TRP1, CEN/ARS] | DH5α | This study | Depletion-complementation |
| bARE53 | pRS414 [*spt6ΔS1*-3xFLAG, TRP1, CEN/ARS] | DH5α | This study | Depletion-complementation |
| bARE51 | pRS414 [*spt6ΔtSH2*-3xFLAG, TRP1, CEN/ARS] | DH5α | This study | Depletion-complementation |
| pFA6-3xHA-KanMX6 | pFA6 [3xHA, KanMX6] | DH5α | PMID: 9717240 | Tagging SET2 for ChIP-seq |
| FB2721 | AmpR Ura3 CEN SET2 | DH5α | This study | Cloning |
| pML107 | Cas9, sgRNA, Leu2, 2μ | DH5α | PMID: 26305040 | CRISPR-Cas9 genome editing |
| pMCSG7 | His6, TEV cleavage site | DH5α | PMID: 12071693 | Protein expression (NMR) |
| Set2-pMCSG7 | His6, TEV cleavage site, Set2(33-260) | DH5α | This study | Protein expression (NMR and HDX) |
| Set2(33-260)-pMCSG7 | His6, TEV cleavage site, Set2(33-260) | DH5α | This study | Protein expression (NMR) |
| Set2(260-455)-pMCSG7 | His6, TEV cleavage site, Set2(260-455) | DH5α | This study | Protein expression (NMR) |

| | | | | |
|---|---|---|---|---|
| Set2(33-455)-pMCSG7 | His6, TEV cleavage site, Set2(33-455) | DH5α | This study | Protein expression (NMR) |
| Set2(I89L)-pMCSG7 | His6, TEV cleavage site, Set2(I89L) | DH5α | This study | Protein expression (NMR) |
| Set2(N90A)-pMCSG7 | His6, TEV cleavage site, Set2(N90A) | DH5α | This study | Protein expression (NMR) |
| Set2(L92F)-pMCSG7 | His6, TEV cleavage site, Set2(L92F) | DH5α | This study | Protein expression (NMR) |
| Set2(L92H)-pMCSG7 | His6, TEV cleavage site, Set2(L92H) | DH5α | This study | Protein expression (NMR) |
| Set2(L92I)-pMCSG7 | His6, TEV cleavage site, Set2(L92I) | DH5α | This study | Protein expression (NMR) |
| Set2(N208D)-pMCSG7 | His6, TEV cleavage site, Set2(N208D) | DH5α | This study | Protein expression (NMR) |
| Set2(N208Y)-pMCSG7 | His6, TEV cleavage site, Set2(N208Y) | DH5α | This study | Protein expression (NMR) |
| Set2(G260D)-pMCSG7 | His6, TEV cleavage site, Set2(G260D) | DH5α | This study | Protein expression (NMR and HDX) |
| Set2(I89L/N208Y)-pMCSG7 | His6, TEV cleavage site, Set2(I89L/N208Y) | DH5α | This study | Protein expression (NMR) |
| Set2(I89L/G260D)-pMCSG7 | His6, TEV cleavage site, Set2(I89L/G260D) | DH5α | This study | Protein expression (NMR) |
| H2A-pMCSG7 | histone H2A (*X. laevis*) | DH5α | PMID: 40972526 | Protein expression (NMR) |
| H2B-pMCSG7 | histone H2B (*X. laevis*) | DH5α | PMID: 40972526 | Protein expression (NMR) |
| H3-pMCSG7 | histone H3 (*X. laevis*) | DH5α | PMID: 40972526 | Protein expression (NMR) |
| H4-pMCSG7 | histone H4 (*X. laevis*) | DH5α | PMID: 40972526 | Protein expression (NMR) |
| pRARE2 | tRNA, chloramphenicol resistance | DH5α | PMID: 18289875 | Protein expression (NMR) |

**Supplementary Table S4.** *SET2* suppressor alleles of *spt6-RR*.

| Plasmid: | Whole plasmids sequencing results: | | | | | | Changes made in parental strain: |
|---|---|---|---|---|---|---|---|
| 1 | K225K | G248C | I275N | | | | |
| 2 | K288STOP | | | | | | |
| 3 | F30F | **L92I** | E137E | K228I | | | L92I (C>A) |
| 4 | I51L | E156E | **G260D** | | | | G260D (G>A) |
| 5 | K43E | **L92F** | D100V | Q273L | | | L92F (C>T) |
| 6 | **L92F** | | | | | | L92F (C>T) |
| 7 | L279S | | | | | | |
| 8 | **N90D** | Q112R | V153V | V281M | | | N90D (A>G) |
| 9 | S202S | K288STOP | | | | | |
| 10 | E33V | **L92H** | **N208D** | P272S | | | N208D (A>G) or L92H (T>A) |
| 11 | Y52Y | L279S | | | | | |
| 12 | **L92F** | | | | | | L92F (C>T) |
| 13 | L279S | | | | | | |
| 14 | F65L | W289STOP | | | | | |
| 15 | I16I | L295STOP | | | | | |
| 16 | D277STOP | | | | | | |
| 17 | S8R | F30I | G106R | G133G | A245P | P272P | |
| 18 | S10L | Q32E | **L92F** | Q138K | S284S | | L92F (C>T) |
| 19 | A40V | **I89L** | A123T | V212A | G219G | | I89L (A>C) |
| 20 | E22G | D64V | L92F | | | | |
| 21 | G251D | N274K | | | | | |
| 22 | D83N | **N208Y** | | | | | N208Y (A>T) |
| 23 | D100D | K291STOP | | | | | |

**Supplementary Table S5.** RNA-seq results, showing all genes with an average log2 fold change >1 or <-1 in at least one genotype for cryptic antisense transcript levels.

| **gene name** | **mean log2 fold-change** | | | |
|---|---|---|---|---|
| | ***set2*Δ/WT** | ***RR*/WT** | ***RRsup*/WT** | ***sup*/WT** |
| HSP26 | 3.717 | 0.1951 | -0.06573 | 3.518 |
| PTI1 | 3.693 | 0.4014 | 0.05014 | 0.1464 |
| RNQ1 | 3.342 | 0.553 | -0.2021 | 0.4636 |
| RAD55 | 2.969 | 2.906 | 0.2198 | 0.1338 |
| LIN1 | 2.836 | 2.945 | -0.2174 | 2.904 |
| YCK3 | 2.793 | 2.914 | 0.101 | 0.1488 |
| ARD1 | 2.668 | -0.07074 | 2.445 | 3.053 |
| VAM3 | 2.607 | 2.59 | -0.1664 | -0.0115 |
| GDH3 | 2.602 | 2.494 | 2.465 | -0.04242 |
| CCW12 | 2.588 | -0.04202 | -0.2625 | 0.1581 |
| YGR283C | 2.584 | 2.736 | 1.625 | 0.2185 |
| HOS1 | 2.525 | 0.1633 | -0.2812 | 1.213 |
| CRP1 | 2.494 | 2.844 | -0.0741 | 2.922 |
| YIF1 | 2.396 | 2.459 | 3.053 | 2 |
| YCH1 | 2.383 | 0.3818 | -0.05246 | 2.805 |
| GAS2 | 2.352 | 3.803 | 2.643 | -0.1172 |
| YHR127W | 2.324 | 2.6 | 1.79 | 0.5576 |
| MSL1 | 2.322 | 2.537 | 2.441 | 2.855 |
| NCE102 | 2.305 | 0.3308 | 2.178 | 2.725 |
| FLX1 | 2.254 | 0.37 | -0.1752 | 2.316 |
| FMC1 | 2.215 | -0.1699 | -0.1766 | 2.863 |
| SUB1 | 2.209 | 0.728 | 1.813 | 2.049 |
| ALG14 | 2.2 | 0.00299 | -0.0253 | 2.932 |
| HEH2 | 2.2 | 2.055 | -0.0756 | 0.2163 |
| MSR1 | 2.184 | 0.2551 | 1.447 | 0.0474 |
| PPH21 | 2.145 | 2.207 | 0.1388 | 1.568 |
| ERG20 | 2.113 | 0.6543 | 0.2695 | 0.573 |
| RXT2 | 2.098 | 1.874 | -0.1343 | 0.52 |
| PSY4 | 2.088 | 2.27 | 1.147 | -0.02048 |
| YLR118C | 2.053 | -0.1656 | -0.10364 | -0.2036 |
| SAM35 | 1.989 | 2.012 | -0.09656 | -0.09393 |
| YKR011C | 1.985 | 2.2 | -0.2917 | 2.438 |
| FES1 | 1.983 | -0.1649 | 0.0983 | 1.842 |
| SBA1 | 1.982 | -0.4304 | 0.8564 | 2.248 |

log2 fold-change > 1

log2 fold-change < -1

*sup=SET2-I89L/G260D*
*RR=spt6-RR*

| | | | | |
|---|---|---|---|---|
| YRR1 | 1.972 | 2.064 | 1.17 | 0.1241 |
| COS12 | 1.959 | 1.759 | -0.06476 | -0.03665 |
| CTO1 | 1.95 | 1.854 | 0.914 | 0.4587 |
| URA3 | 1.948 | -0.4844 | 1.623 | 0.075 |
| ARG80 | 1.9375 | 0.1665 | 1.948 | 2.842 |
| HST3 | 1.935 | 0.437 | -0.08875 | 0.10864 |
| THI72 | 1.923 | 0.651 | -0.0387 | 0.04193 |
| ERV29 | 1.902 | 3.314 | 0.337 | 3.68 |
| THP1 | 1.891 | 1.952 | 0.919 | 0.951 |
| LMO1 | 1.881 | 1.62 | -0.08514 | 0.0923 |
| THI11 | 1.87 | -0.1554 | 1.353 | -0.1968 |
| FMP30 | 1.853 | 0.002306 | 0.01224 | 0.1205 |
| KRS1 | 1.844 | 2.125 | 0.03543 | 0.746 |
| BMH1 | 1.841 | 0.2429 | -0.2664 | 0.5054 |
| CAP2 | 1.835 | 2.03 | 1.553 | 1.997 |
| YKL069W | 1.829 | 1.754 | 1.46 | 1.575 |
| NCE101 | 1.828 | 1.643 | 0 | 0 |
| ADH3 | 1.797 | 0.1223 | -0.2551 | 1.969 |
| HXT9 | 1.794 | -0.2534 | -0.1676 | 0.5513 |
| YMR226C | 1.791 | 0.5625 | 0.03046 | 0 |
| YER134C | 1.769 | 1.411 | 1.33 | -0.0996 |
| SGF29 | 1.767 | 0.1647 | 0.1436 | 2.053 |
| MSS11 | 1.765 | 1.566 | -0.3477 | -0.301 |
| LAM1 | 1.753 | 1.701 | 0.0328 | 0.01894 |
| THI12 | 1.746 | -0.3115 | -0.339 | -0.06366 |
| GRX7 | 1.728 | 1.795 | 1.331 | -0.3103 |
| REV3 | 1.724 | 1.781 | 0.02187 | 0.1552 |
| APD1 | 1.722 | 1.514 | 1.141 | 0.1 |
| SMC2 | 1.72 | 1.377 | 0.01108 | 0.03073 |
| APP1 | 1.713 | 2.32 | 0.008995 | 0.518 |
| SPR2 | 1.704 | 0.1844 | 0.4019 | 3.1 |
| CTT1 | 1.704 | 1.423 | 0.1111 | 0.2045 |
| SIT4 | 1.702 | 0.1289 | 1.45 | 0.1414 |
| DPH2 | 1.698 | 0.1592 | 1.148 | 0.1685 |
| CDC3 | 1.675 | 1.689 | -0.4478 | 1.579 |
| BRR2 | 1.658 | 1.627 | -0.1306 | 0.03513 |
| RIM15 | 1.651 | 1.631 | -0.1224 | -0.1957 |
| BUD9 | 1.647 | 0.5205 | 1.535 | 1.626 |
| RAD61 | 1.645 | 1.318 | 0.004326 | 0.3235 |

| | | | | |
|---|---|---|---|---|
| ZRC1 | 1.622 | 0.1887 | 1.415 | 1.1875 |
| IML1 | 1.62 | 1.634 | -0.05664 | -0.03232 |
| MDL1 | 1.617 | 1.762 | -0.03333 | -0.03552 |
| RKM3 | 1.606 | 0.6694 | -0.2399 | 1.976 |
| ESBP6 | 1.605 | 1.001 | 0.07135 | 0.09937 |
| SSK1 | 1.605 | 1.432 | -0.2869 | -0.28 |
| PET127 | 1.591 | 1.562 | -0.04825 | 0.354 |
| VPS62 | 1.59 | 1.765 | 0.355 | 0.307 |
| GIP4 | 1.581 | 1.699 | -0.2141 | 0.0733 |
| TDA7 | 1.579 | 1.768 | -0.01901 | 0.02129 |
| SPT20 | 1.5625 | 1.524 | 0.05072 | 0.1501 |
| PSP2 | 1.557 | 1.618 | 0.00678 | 0.3274 |
| TAT1 | 1.543 | 0.6514 | 0.774 | 1.057 |
| AVO1 | 1.537 | 1.416 | -0.0362 | 0.000616 |
| KIN4 | 1.521 | 1.109 | 0.666 | 0.8877 |
| YKL050C | 1.52 | 1.304 | 0.01663 | 0.01508 |
| YKL091C | 1.518 | 1.474 | 1.161 | -0.12103 |
| RTT102 | 1.493 | 0.241 | 0.10364 | 1.667 |
| SKP2 | 1.492 | 1.437 | 0.1896 | 0.03021 |
| SSH1 | 1.491 | -0.02756 | -0.0373 | 1.541 |
| MPH1 | 1.488 | 1.464 | -0.01971 | -0.05685 |
| RFC4 | 1.481 | 0.01233 | -0.2754 | 0.8716 |
| SYC1 | 1.476 | 0.31 | 1.426 | 1.784 |
| MUD2 | 1.472 | 1.345 | -0.0934 | 0.17 |
| JHD1 | 1.471 | 1.591 | 0.2668 | 0.0661 |
| IES2 | 1.47 | 0.0577 | 0.03564 | 0.1836 |
| IZH4 | 1.466 | 0.1267 | 0.03058 | -0.1859 |
| SOG2 | 1.462 | 1.717 | -0.0342 | -0.01703 |
| TED1 | 1.46 | 0.396 | -0.1726 | -0.05347 |
| TUS1 | 1.457 | 1.478 | 0.07513 | -0.1171 |
| GYL1 | 1.454 | 1.3955 | -0.1632 | 0.09454 |
| HBS1 | 1.452 | 1.137 | -0.0447 | 0.1569 |
| YKU80 | 1.452 | 1.24 | 1.052 | 0.2976 |
| CLP1 | 1.449 | 1.416 | -0.1914 | -0.1192 |
| RPS4B | 1.447 | -0.1255 | -0.0876 | 0.1622 |
| SPS1 | 1.444 | -0.149 | 0.5024 | -0.11444 |
| CCL1 | 1.431 | 1.516 | 0.288 | 0.3757 |
| MRPL31 | 1.419 | -0.10065 | -0.04532 | -0.0781 |
| MRPL24 | 1.415 | 1.771 | 0.9478 | -0.2072 |

| | | | | |
|---|---|---|---|---|
| POL31 | 1.412 | 1.333 | 0.3445 | 0.3025 |
| BIR1 | 1.41 | 1.537 | -0.1126 | 0.0358 |
| ORC3 | 1.396 | 1.494 | 0.7944 | 0.2444 |
| ABZ1 | 1.3955 | 0.5273 | 0.793 | 1.226 |
| RTT101 | 1.395 | 1.074 | -0.04062 | 0.1909 |
| RAD30 | 1.391 | 1.562 | 0.1694 | 0.2399 |
| TIM23 | 1.383 | -0.2134 | -0.06836 | 0.5522 |
| CDC26 | 1.382 | 1.32 | -0.07935 | 0.2281 |
| AAP1 | 1.382 | 1.277 | 0.07166 | 0.1849 |
| GUT1 | 1.38 | 1.595 | -0.0633 | 0.1147 |
| VMA10 | 1.364 | 2.021 | 0.11426 | 1.822 |
| SGV1 | 1.363 | 1.408 | 0.01135 | -0.0626 |
| PEX5 | 1.36 | 1.284 | -0.0379 | -0.10004 |
| SSY1 | 1.355 | 1.282 | 0.08704 | 0.073 |
| MDV1 | 1.342 | 0.1436 | -0.0879 | -0.0143 |
| UAF30 | 1.341 | 0.01778 | 1.511 | 0.1166 |
| PCA1 | 1.34 | 1.065 | -0.013504 | 0.05252 |
| MDM36 | 1.334 | 1.33 | 0.204 | 0.3452 |
| GPN3 | 1.333 | 0.10895 | 0.5454 | 0.02611 |
| ATG2 | 1.329 | 1.43 | -0.1469 | -0.0991 |
| RGA1 | 1.318 | 1.05 | -0.006023 | 0.1393 |
| DCC1 | 1.315 | 0.551 | 1.194 | -0.014626 |
| AIM21 | 1.315 | 1.023 | -0.2583 | -0.3528 |
| MON1 | 1.311 | 1.469 | -0.01828 | -0.3608 |
| LSP1 | 1.304 | 0.2139 | 0.869 | 1.2 |
| ARO80 | 1.303 | 1.245 | -0.01399 | 0.3645 |
| VPS35 | 1.303 | 1.59 | 0.09045 | 0.2299 |
| CCH1 | 1.303 | 1.172 | 0.01726 | 0.02385 |
| MET30 | 1.299 | 0.824 | -0.2023 | -0.0862 |
| RIO2 | 1.297 | 1.804 | 0.5483 | 0.4673 |
| DSS1 | 1.291 | 1.335 | -0.04385 | 0.03473 |
| MTF1 | 1.288 | 0.89 | 0.0795 | 1.272 |
| IMA1 | 1.286 | -0.4233 | 0.2546 | -0.57 |
| BPT1 | 1.284 | 1.382 | 0.06137 | 0.163 |
| FMP16 | 1.282 | -0.2004 | 0.5293 | 1.407 |
| ZRG8 | 1.282 | 1.3 | -0.004154 | 0.00796 |
| NPR3 | 1.282 | 1.288 | -0.03317 | 0.093 |
| AIM46 | 1.277 | 0.854 | 0.3667 | 0.275 |
| RRT2 | 1.267 | 1.42 | 0.2343 | 0.547 |

| | | | | |
|---|---|---|---|---|
| RIF1 | 1.264 | 1.215 | -0.041 | 0.1705 |
| BNR1 | 1.262 | 0.9565 | -0.04276 | -0.00783 |
| SPC105 | 1.258 | 1.125 | 0.0491 | 0.0674 |
| RUB1 | 1.255 | -1.778 | -2.166 | -1.886 |
| BDP1 | 1.253 | 0.7847 | 1.131 | 1.512 |
| BSP1 | 1.251 | 1.081 | -0.1326 | -0.02625 |
| RAD2 | 1.25 | 1.113 | -0.135 | -0.1028 |
| PPH22 | 1.248 | 0.03017 | 0.00906 | 0.0901 |
| TAF4 | 1.24 | 1.258 | 0.01944 | 0.4333 |
| YBL111C | 1.235 | 0.03134 | 0.7314 | 0.4863 |
| ATG4 | 1.234 | 0.00626 | 1.268 | 1.773 |
| UBX7 | 1.231 | 1.286 | 1.021 | 0.04825 |
| BOI2 | 1.222 | 1.033 | -0.0804 | -0.0678 |
| PEF1 | 1.219 | 1.02 | -0.05463 | -0.0816 |
| HCR1 | 1.217 | 0.2837 | 0.001059 | 0.3035 |
| SER33 | 1.217 | 0.3198 | -0.0195 | 0.3667 |
| YNL122C | 1.215 | 1.226 | 1.191 | 1.224 |
| DNF3 | 1.214 | 1.162 | -0.06033 | -0.03268 |
| EFG1 | 1.211 | -0.1359 | 1.13 | 0.00825 |
| YCP4 | 1.211 | 0.9897 | 0.9575 | 1.418 |
| ZDS2 | 1.211 | 1.097 | 0.1761 | 0.2314 |
| LTE1 | 1.208 | 1.071 | -0.01308 | 0.02939 |
| RTC1 | 1.207 | 0.939 | 0.08844 | 0.0966 |
| PRP9 | 1.206 | 1.396 | -0.07104 | 0.0562 |
| ESP1 | 1.202 | 1.096 | 0.00943 | 0.001273 |
| ORM1 | 1.2 | 1.551 | 1.332 | -0.1827 |
| YHR182W | 1.2 | 0.941 | 0.01846 | 0.4402 |
| MEC1 | 1.199 | 1.128 | 0.03065 | 0.0124 |
| VPS41 | 1.194 | 1.244 | -0.2217 | -0.10144 |
| GLE1 | 1.193 | 0.851 | 0.04483 | -0.03647 |
| TMA108 | 1.191 | 1.127 | 0.002188 | 0.0799 |
| MF(ALPHA)2 | 1.19 | -0.2954 | -0.3567 | 0.004047 |
| NCA2 | 1.19 | 1.271 | 0.1111 | 0.1383 |
| CEM1 | 1.189 | 1.533 | -0.0489 | 1.593 |
| CHA4 | 1.189 | 1.158 | -0.265 | -0.1186 |
| AAD10 | 1.187 | 0.532 | -0.03964 | 0.9307 |
| MNT3 | 1.185 | 0.965 | 0.038 | -0.06586 |
| DSL1 | 1.184 | 1.103 | 0.2119 | -0.0453 |
| BNI4 | 1.184 | 0.958 | 0.0494 | 0.1225 |

| | | | | |
|---|---|---|---|---|
| SSL2 | 1.183 | 1.118 | -0.0001268 | 0.1655 |
| ESC1 | 1.178 | 1.108 | 0.01344 | 0.1959 |
| TGL1 | 1.177 | 1.038 | 0.01371 | -0.0313 |
| AAD4 | 1.176 | 1.216 | -0.1165 | -0.1326 |
| ISA1 | 1.172 | 0.4226 | 0.968 | 1.355 |
| BCS1 | 1.166 | -0.7466 | -0.2676 | 0.287 |
| PMD1 | 1.164 | 1.117 | 0.014206 | 0.09735 |
| SYF1 | 1.163 | 1.591 | -0.2183 | -0.08136 |
| SKI2 | 1.163 | 0.987 | -0.008125 | 0.1175 |
| HSP33 | 1.162 | 0.1388 | 0.7476 | 0.004345 |
| ABP1 | 1.162 | 0.6626 | -0.1439 | 0.01338 |
| VPS34 | 1.161 | 1.19 | 0.0666 | 0.1525 |
| ARC18 | 1.159 | 1.33 | 0.17 | 0.02637 |
| NTO1 | 1.156 | 0.9424 | 0.0986 | 0.2468 |
| SPO77 | 1.155 | 1.07 | -0.597 | 1.396 |
| MSC3 | 1.15 | 1.221 | 0.06775 | 0.01063 |
| CYK3 | 1.149 | 1.169 | 0.1036 | 0.1647 |
| VPS8 | 1.147 | 1.324 | 0.02573 | -0.04025 |
| SSL1 | 1.145 | 1.23 | -0.07294 | 0.3364 |
| RAD9 | 1.139 | 1.047 | 0.01782 | 0.001546 |
| DAK2 | 1.138 | 0.636 | 0.8184 | 0.7075 |
| NST1 | 1.136 | 0.823 | 0.1287 | 0.1098 |
| OXA1 | 1.134 | 1.287 | -0.1779 | 0.135 |
| INP2 | 1.133 | 1.224 | 0.1873 | 0.2233 |
| RHB1 | 1.132 | 0.191 | 0.8755 | -0.06158 |
| PCL8 | 1.128 | 1.003 | -0.1329 | 0.08936 |
| YIL108W | 1.128 | 0.9663 | -0.1526 | -0.11774 |
| MET1 | 1.125 | 1.17 | -0.2852 | -0.2932 |
| SAC3 | 1.125 | 1.067 | 0.0376 | 0.1271 |
| PLB2 | 1.122 | 1.047 | -0.0765 | 0.2605 |
| KSS1 | 1.117 | 0.932 | 1.02 | 0.4595 |
| SCD5 | 1.116 | 1.112 | -0.3875 | -0.343 |
| RKR1 | 1.116 | 1.138 | -0.095 | 0.08777 |
| SAC1 | 1.113 | 1.23 | 0.03018 | 0.2317 |
| ATG1 | 1.113 | 1.221 | -0.08307 | -0.09503 |
| RAD5 | 1.105 | 1.184 | -0.0472 | 0.0971 |
| PIS1 | 1.101 | 1.008 | 0.196 | 1.199 |
| ARE2 | 1.1 | 1.421 | 0.00406 | 1.149 |
| USV1 | 1.098 | 1.265 | 1.192 | 0.631 |

| | | | | |
|---|---|---|---|---|
| STB4 | 1.098 | 1.025 | 0.0576 | -0.0643 |
| APL5 | 1.097 | 0.8896 | -0.03174 | 0.02266 |
| ECM25 | 1.092 | 1.412 | -0.2408 | 0.00887 |
| YLR179C | 1.091 | 0.91 | 0.663 | 1.286 |
| AVL9 | 1.091 | 0.8804 | -0.02716 | 0.010574 |
| RPL38 | 1.087 | 1.1455 | 1.148 | -0.1903 |
| ASI1 | 1.083 | 0.831 | 0.09375 | 0.2112 |
| FAB1 | 1.079 | 0.94 | 0.02074 | 0 |
| TBS1 | 1.073 | 1.199 | 0.10834 | 0.1771 |
| SIR4 | 1.073 | 0.977 | -0.02779 | 0.04184 |
| FRK1 | 1.072 | 0.895 | -0.1724 | 0.1852 |
| POL1 | 1.071 | 1.066 | -0.10834 | -0.0915 |
| HRD3 | 1.067 | 1.003 | -0.06396 | -0.2496 |
| RGC1 | 1.066 | 1.092 | -0.02605 | 0.03906 |
| LHS1 | 1.064 | 0.8525 | 0.03214 | -0.03433 |
| DAK1 | 1.063 | 1 | 0.08826 | 0.08417 |
| HUL5 | 1.0625 | 0.914 | -0.004906 | 0.001252 |
| VAC7 | 1.061 | 1.023 | -0.01094 | 0.2974 |
| UBR2 | 1.06 | 0.675 | 0.06915 | 0.12244 |
| MDR1 | 1.059 | 1.097 | 0.002848 | 0.1058 |
| RAD23 | 1.055 | 3.076 | 2.607 | 0.4028 |
| BOI1 | 1.05 | 0.862 | -0.03053 | -0.03867 |
| QDR3 | 1.046 | 0.809 | 0.146 | 0.2424 |
| SPO75 | 1.046 | 1.018 | -0.0958 | 0.2351 |
| VAM6 | 1.044 | 1.045 | -0.0915 | 0.1344 |
| SWH1 | 1.044 | 0.4644 | -0.05652 | -0.02371 |
| DYN1 | 1.043 | 1.065 | -0.002974 | 0.002466 |
| AVT4 | 1.039 | 1.084 | 0.02724 | -0.1688 |
| SPT7 | 1.037 | 0.7036 | -0.071 | -0.01732 |
| TRS65 | 1.035 | 1.257 | -0.1475 | -0.2242 |
| ATG13 | 1.034 | 1.25 | 0.0481 | 0.0794 |
| IME1 | 1.032 | 0.1771 | 0.513 | 0.01907 |
| GAS3 | 1.031 | 0.965 | -0.03607 | -0.131 |
| DUO1 | 1.028 | -1.558 | -0.1587 | -0.1316 |
| ECM22 | 1.028 | 0.3835 | -0.1154 | -0.1625 |
| LAM4 | 1.028 | 1.107 | 0.0836 | 0.03806 |
| ARP4 | 1.027 | 0.8164 | -0.3164 | 0.2399 |
| TEL1 | 1.026 | 0.8564 | 0.00792 | 0.0597 |
| NFI1 | 1.025 | 0.9346 | -0.01491 | 0.09424 |

| | | | | |
|---|---|---|---|---|
| COQ6 | 1.024 | 0.8906 | 0.2286 | 0.11896 |
| AKL1 | 1.022 | 0.8804 | -0.00489 | 0.1438 |
| YJR084W | 1.021 | 1.046 | -0.1643 | -0.0629 |
| APM2 | 1.0205 | 0.9688 | 0.05734 | 0.08563 |
| CSS2 | 1.016 | 1.046 | -0.3608 | 0.1 |
| PHO8 | 1.016 | 0.8643 | 0.2349 | 0.26 |
| GEA1 | 1.013 | 0.982 | 0.10516 | 0.1917 |
| SLH1 | 1.011 | 0.954 | -0.003593 | -0.00996 |
| SSQ1 | 1.009 | 1.171 | 0.04733 | 0.079 |
| MNT2 | 1.005 | 1.037 | 0.2421 | 0.3257 |
| LYP1 | 1.005 | 1.058 | 0.1663 | 0.1663 |
| NTG1 | 1.004 | 1.243 | 0.2925 | 0.01585 |
| NAM2 | 1.002 | 0.6987 | -0.0425 | -0.02396 |
| SEC3 | 1.002 | 0.717 | 0.09204 | 0.02762 |
| GTB1 | 1.001 | -0.1536 | 0.659 | 0.6064 |
| DIE2 | 1 | 0.8486 | 0.0474 | 0.1671 |
| PPZ2 | 1 | 1.09 | 0.0966 | 0.0818 |
| TOK1 | 0.9937 | 1.129 | 0.0357 | 0.1849 |
| MHP1 | 0.985 | 1.054 | -0.02235 | 0.1329 |
| AFG1 | 0.977 | 1.036 | -0.0159 | 0.4272 |
| MEP1 | 0.975 | 0.8584 | 2.057 | 2.441 |
| CLN1 | 0.9717 | 1.247 | -0.02534 | 0.1342 |
| RIB2 | 0.968 | 1.593 | -0.1312 | 1.509 |
| GAL80 | 0.9653 | 1.113 | 0.1709 | 0.2278 |
| LAM5 | 0.962 | 1.146 | 0.02145 | -0.0696 |
| BYE1 | 0.9614 | 1.038 | 0.2305 | 0.3447 |
| REC104 | 0.958 | 1.068 | 0.255 | 1.039 |
| IRC22 | 0.944 | 1.121 | -0.0344 | 0.2036 |
| YRO2 | 0.9414 | 1.186 | 0.22 | 0.3323 |
| HXT16 | 0.9375 | 0.0872 | 2.115 | 2.047 |
| MIG1 | 0.933 | 1.183 | -0.1486 | 1.101 |
| PCL2 | 0.927 | 1.133 | 0.3936 | 0.5093 |
| COS10 | 0.927 | 0.1394 | -0.0713 | 1.012 |
| TRF5 | 0.9253 | 1.069 | 0.1378 | 0.349 |
| DIG2 | 0.924 | 1.277 | -0.3572 | 0.02913 |
| NCS6 | 0.92 | 1.102 | 0.151 | 0.546 |
| ATG9 | 0.92 | 1.195 | -0.2211 | -0.0882 |
| PAU22 | 0.919 | 1.936 | 1.083 | 0 |
| RFA2 | 0.9062 | 1.022 | 0.1421 | 0.2007 |

| | | | | |
|---|---|---|---|---|
| YCT1 | 0.9053 | 1.03 | -0.05304 | -0.08484 |
| KRE5 | 0.8984 | 1.009 | 0.01953 | -0.035 |
| SIS1 | 0.8965 | 1.208 | 1.063 | 0.45 |
| ORC4 | 0.891 | 1.109 | -0.0201 | 0.1708 |
| EPO1 | 0.886 | 1.093 | 0.02805 | 0.1292 |
| MAK11 | 0.877 | 2.096 | 1.539 | 1.885 |
| ATP25 | 0.867 | 1.572 | 0.1632 | 0.05463 |
| RCE1 | 0.8647 | 1.153 | 0.325 | 0.905 |
| SEC62 | 0.8555 | 0.6724 | 0.1302 | 1.358 |
| CHS6 | 0.833 | 1.019 | -0.03772 | -6.65E-05 |
| NRT1 | 0.83 | 1.096 | -0.1503 | -0.422 |
| YPR148C | 0.8296 | 0.4153 | 1.376 | -0.1143 |
| IOC2 | 0.814 | 1.022 | -0.04053 | 0.6025 |
| MET6 | 0.8105 | 1.018 | -0.07764 | -0.04306 |
| SET5 | 0.785 | 1.524 | -0.343 | -0.1521 |
| LSB3 | 0.7715 | 1.074 | 0.1854 | 0.01605 |
| TPC1 | 0.751 | 1.143 | 0.0742 | 0.4993 |
| ATG23 | 0.751 | 1.195 | 0.171 | -0.05234 |
| GLR1 | 0.739 | 1.1045 | 0.2146 | 0.396 |
| MCP1 | 0.7334 | 1.347 | 0.3135 | 0.2191 |
| FLO5 | 0.7334 | 1.086 | 0.02068 | -0.02615 |
| DBF2 | 0.73 | 1.054 | 0.0003114 | -0.04 |
| NSI1 | 0.729 | 1.021 | -0.1569 | -0.2725 |
| PDE1 | 0.7256 | 1.3125 | -0.04517 | 1.887 |
| GAL4 | 0.715 | 2.123 | 0.2896 | 0.3467 |
| PET20 | 0.697 | 1.129 | 0.4285 | 0.543 |
| GFD1 | 0.6963 | 0.8105 | 0.655 | 1.047 |
| FAR7 | 0.6855 | 0.539 | -0.04047 | 1.616 |
| RML2 | 0.6836 | 1.348 | -0.2742 | -0.1307 |
| HEM1 | 0.6743 | 1.505 | 1.499 | 1.844 |
| SNF6 | 0.671 | 1.13 | -0.00534 | 1.169 |
| SNF12 | 0.6523 | 0.846 | 1.412 | -0.02757 |
| RRM3 | 0.6323 | 2.332 | 0.0594 | 1.505 |
| PZF1 | 0.6045 | 1.602 | -0.006836 | -0.00895 |
| ATS1 | 0.5938 | 1.08 | 0.1212 | 0.979 |
| LAS17 | 0.5737 | 1.41 | 0.05817 | 0.2612 |
| RPS30B | 0.5674 | 0.2323 | -0.02728 | 1.004 |
| YJR056C | 0.5415 | 0.373 | -0.02316 | 1.478 |
| STB5 | 0.5376 | 1.676 | 0.163 | 1.149 |

| | | | | |
|---|---|---|---|---|
| DAM1 | 0.506 | 0.8853 | 1.404 | 1.771 |
| SWM1 | 0.5054 | 0.2091 | 1.039 | 1.6455 |
| PRP6 | 0.4912 | 0.448 | 1.652 | 2.195 |
| JLP1 | 0.4812 | 1.614 | 1.116 | 0.04254 |
| THG1 | 0.4536 | 0.426 | 1.131 | -0.009735 |
| ARL1 | 0.4473 | 1.174 | 0.0856 | 1.473 |
| TTI2 | 0.429 | 1.437 | 0.1451 | 0.632 |
| ZIP2 | 0.4272 | 1.223 | 0.05884 | 0.1985 |
| RPL19B | 0.4165 | 1.495 | 0.5137 | 1.21 |
| HMG2 | 0.4133 | 1.967 | 0.6724 | 0.01817 |
| STP1 | 0.4102 | 0.335 | 0.005024 | 2.223 |
| GEP3 | 0.4011 | 0.08026 | 0.3425 | 1.289 |
| RPL20A | 0.3962 | 0.6353 | 0.4048 | 1.004 |
| SKO1 | 0.395 | 0.26 | 0.6274 | 1.272 |
| ADH2 | 0.3945 | 2.076 | 0.2361 | 3.027 |
| RSM25 | 0.3894 | 0.5503 | 0.861 | 1.05 |
| WWM1 | 0.3755 | 1.354 | 0.8667 | 0.6157 |
| RPS5 | 0.3696 | 0.3198 | 2.223 | 0.4978 |
| THP3 | 0.3538 | 1.438 | -0.04807 | 0.7153 |
| MED6 | 0.3528 | 1.005 | -0.0695 | 0.1346 |
| VMA13 | 0.343 | 0.1713 | -0.0274 | 2.371 |
| NUP60 | 0.3303 | 1.525 | 0.656 | 0.646 |
| YSR3 | 0.3184 | 0.3972 | -0.09924 | 3.482 |
| PEX8 | 0.2969 | 1.287 | -0.1692 | 1.189 |
| VAM7 | 0.2903 | 0.318 | 0.01648 | 2.64 |
| SNP1 | 0.2803 | 1.76 | 1.591 | 0.0643 |
| POM33 | 0.2751 | 1.142 | 0.2812 | 0.54 |
| RNH70 | 0.2454 | 1.351 | 0.2993 | 0.6685 |
| HMF1 | 0.2318 | 0.09357 | 2.223 | -0.109 |
| TRM10 | 0.228 | 3.164 | 2.797 | 3.525 |
| SIP2 | 0.2266 | 1.58 | -0.005764 | 1.572 |
| PRE6 | 0.2161 | 1.237 | 0.0754 | -0.0704 |
| PXP2 | 0.2139 | 1.77 | 1.537 | -0.3757 |
| AVT5 | 0.211 | 1.8125 | 0.836 | 0.1675 |
| COX6 | 0.2108 | 2.28 | 2.271 | 2.732 |
| TPN1 | 0.2083 | 1.553 | 0.08624 | 0.0705 |
| POL4 | 0.2074 | 1.16 | 0.2489 | 0.428 |
| FEN2 | 0.2046 | 1.231 | -0.04376 | 0.10583 |
| TFB6 | 0.1973 | 1.703 | -0.327 | 1.288 |

| | | | | |
|---|---|---|---|---|
| PET10 | 0.1915 | 0.0727 | 3.941 | 4.56 |
| RRP3 | 0.1869 | 2.082 | 0.0869 | 0.06207 |
| RPT1 | 0.1804 | 1.244 | 1.164 | 1.8125 |
| MND1 | 0.1732 | 0.2019 | 1.495 | 0.2505 |
| NAT4 | 0.1653 | -0.09827 | 0.2925 | 1.35 |
| DUT1 | 0.1543 | 1.1875 | 1.176 | -0.3303 |
| AIR1 | 0.1481 | 1.506 | 0.2435 | 0.227 |
| SWA2 | 0.1442 | 1.648 | -0.1346 | 1.146 |
| LCB1 | 0.1437 | 1.378 | -0.275 | 0.899 |
| YRB1 | 0.1109 | 0.531 | 0.1536 | 1.048 |
| MSW1 | 0.10425 | 0.03662 | -0.4688 | 1.057 |
| MSS116 | 0.1029 | 1.783 | 0.621 | 0.759 |
| ECM11 | 0.0785 | 1.664 | 0.2798 | 0.003416 |
| MRPL44 | 0.0762 | 1.388 | 1.343 | 1.212 |
| MRPL23 | 0.07556 | 0.84 | -0.786 | 1.218 |
| NMT1 | 0.06537 | 0.3188 | -0.5596 | 1.199 |
| THI13 | 0.06174 | 0.2323 | 0.01242 | 1.799 |
| KAP95 | 0.0511 | 1.133 | -0.087 | -0.02585 |
| GET2 | 0.04434 | 2.793 | -0.121 | 0.3833 |
| UTR4 | 0.02669 | 1.031 | 0.9424 | 0.1569 |
| NAT2 | 0.025 | -0.2056 | -0.0984 | 1.385 |
| PAC10 | 0.02237 | 1.222 | 0.1306 | -0.1011 |
| YPS6 | 0.02126 | 1.477 | 0.5938 | 0.908 |
| NNR1 | 0.01648 | 0.3994 | 1.003 | 0.8555 |
| TIM22 | 0.003069 | 1.19 | 0.854 | -0.0923 |
| RPS29B | 0 | 3.928 | 0 | 3.92 |
| YHM2 | 0 | 0.05954 | 1.403 | 0 |
| YFH7 | 0 | 0 | 0.0066 | 2.104 |
| COS4 | 0 | 2.146 | 0 | 0 |
| SPO13 | -0.000713 | 0.2112 | 0.5063 | 1.898 |
| PIC2 | -0.000957 | 1.927 | -0.0641 | 2.43 |
| MKS1 | -0.004738 | 0.512 | -0.4634 | 1.371 |
| HSP32 | -0.01949 | 1.841 | -0.010544 | 0.00815 |
| RRN11 | -0.02711 | 1.032 | -0.02078 | 0.2837 |
| ATG41 | -0.02736 | 1.337 | -0.1472 | 1.298 |
| SHE3 | -0.0531 | 1.038 | -0.2068 | -0.1304 |
| BAG7 | -0.05475 | 1.849 | 1.508 | 2.139 |
| PET122 | -0.05527 | 1.673 | 1.523 | 2.262 |
| SUI1 | -0.0687 | 2.473 | 1.698 | 2.318 |

| | | | | |
|---|---|---|---|---|
| RPL5 | -0.0724 | -0.207 | -0.6084 | 1.343 |
| TOA2 | -0.07733 | 1.803 | 1.111 | 0.225 |
| MRPL13 | -0.08466 | 3.508 | 3.344 | 3.945 |
| HTA2 | -0.0925 | 0.06064 | 1.374 | 1.492 |
| THS1 | -0.09503 | 1.315 | -0.2302 | -0.1578 |
| PNC1 | -0.1018 | 0.03854 | 1.668 | -0.07916 |
| ERV1 | -0.1345 | 1.302 | 1.218 | 0.589 |
| SAS2 | -0.1366 | 1.332 | 0.716 | -0.1127 |
| ATG18 | -0.1497 | 1.892 | -0.02213 | 0.1571 |
| ARR1 | -0.1528 | 2.033 | -0.06305 | 0.01064 |
| GUD1 | -0.1605 | 1.598 | -0.41 | -0.3606 |
| FUB1 | -0.1718 | 1.204 | 1.179 | 1.161 |
| TOS8 | -0.1786 | 1.643 | -0.416 | 0.948 |
| TIM11 | -0.1852 | 2.926 | 1.381 | 1.975 |
| PRM9 | -0.1873 | -0.3784 | 0.5713 | 1.054 |
| REE1 | -0.1951 | 1.502 | 1.322 | 0.373 |
| RPS23A | -0.1968 | 0.7617 | -0.05704 | 1.785 |
| FYV6 | -0.2036 | -0.05942 | 0.08563 | 1.285 |
| POR2 | -0.2084 | 0.1798 | 1.413 | -0.07904 |
| FMP45 | -0.2279 | 1.534 | 1.046 | 1.223 |
| PDX3 | -0.2306 | 0.01863 | 1.606 | -0.0477 |
| MRT4 | -0.2327 | -0.3596 | -0.2086 | 2.371 |
| RKM2 | -0.2732 | 1.357 | -0.01194 | 0.0766 |
| RPS8B | -0.289 | -0.1403 | 1.387 | 1.608 |
| ADH7 | -0.3096 | 0.337 | -0.1226 | 1.139 |
| PGU1 | -0.3118 | 1.096 | -0.3628 | -0.2197 |
| YJR085C | -0.316 | -0.02956 | -0.2324 | 1.367 |
| SNO4 | -0.3323 | 0.9814 | -0.3323 | 1.329 |
| SWM2 | -0.3513 | 1.329 | 0.1075 | 1.806 |
| NDE2 | -0.3618 | 1.149 | -0.4866 | 0.02556 |
| RPL17A | -0.3735 | -0.267 | -0.2722 | 1.48 |
| KAR1 | -0.3796 | -0.3882 | -0.5005 | 1.128 |
| SVP26 | -0.427 | 1.726 | 1.3955 | -0.4846 |
| RBG1 | -0.4377 | 0.0922 | -0.5347 | 2.053 |
| ZIM17 | -0.4536 | 1.183 | 1.02 | -0.06055 |
| CSE4 | -0.4844 | 2.025 | -0.5435 | -1.159 |
| DCV1 | -0.522 | -0.2507 | 3.207 | 0.236 |
| PRO2 | -0.5464 | 1.027 | -0.4673 | 0.3708 |
| NNR2 | -0.571 | -0.4045 | 2.11 | 2.709 |

| | | | | |
|---|---|---|---|---|
| MAM1 | -0.6333 | 1.491 | 1.145 | -0.7305 |
| MAS1 | -0.6636 | 1.091 | 0.2988 | -0.8867 |
| RRI1 | -0.6777 | 1.505 | 0.3232 | 0.1357 |
| CRG1 | -1.157 | -0.0958 | 0.2167 | 1.178 |
| SDS3 | -1.745 | -1.711 | 1.521 | -1.318 |
| RIF2 | -2.012 | -1.896 | -0.3599 | 1.533 |
